# Liver Protein Expression in Nash Mice on a High-Fat Diet

**DOI:** 10.1101/2022.01.29.478332

**Authors:** James Varani, Shannon D McClintock, Randall N Knibbs, Isabelle Harber, Dania Zeidan, Mohamed Ali H Jawad-Makki, Muhammad N Aslam

## Abstract

Male MS-NASH mice were maintained on a high-fat diet for 16 weeks with and without red algae-derived minerals. Obeticholic acid (OCA) was used as a comparator in the same strain and diet. C57BL/6 mice maintained on a standard (low-fat) rodent chow diet were used as a control. At the end of the in-life portion of the study, body weight, liver weight, liver enzyme levels and liver histology were assessed. Samples obtained from individual livers were subjected to Tandem Mass Tag labeling / mass spectroscopy for protein profile determination. As compared to mice maintained on the low-fat diet, all high-fat-fed mice had increased whole body and liver weight, increased liver enzyme (aminotransferases) levels and widespread steatosis / ballooning hepatocyte degeneration. Histological evidence for liver inflammation and collagen deposition was also present, but changes were to a lesser extent. A moderate reduction in ballooning degeneration and collagen deposition was observed with mineral supplementation. Control mice on the high-fat diet alone demonstrated multiple protein changes associated with dysregulated fat and carbohydrate metabolism, lipotoxicity and oxidative stress. Cholesterol metabolism and bile acid formation were especially sensitive to diet. In mice receiving multi-mineral supplementation along with the high-fat diet, there was reduced liver toxicity as evidenced by a decrease in levels of several cytochrome P450 enzymes and other oxidant-generating moieties. Additionally, elevated expression of several keratins was also detected in mineral-supplemented mice. The protein changes observed with mineral supplementation were not seen with OCA. Our previous studies have shown that mice maintained on a high-fat diet for up to 18 months develop end-stage liver injury including hepatocellular carcinoma. Mineral-supplemented mice were substantially protected against tumor formation and other end-state consequences of high-fat feeding. The present study identifies early (16-week) protein changes occurring in the livers of the high-fat diet-fed mice, and how the expression of these proteins is influenced by mineral supplementation. These findings help elucidate early protein changes that contribute to end-stage liver injury and potential mechanisms by which dietary minerals may mitigate such damage.

## INTRODUCTION

Non-alcoholic fatty liver disease (NAFLD) is rapidly becoming the most common cause of liver injury in Western society and throughout the world (1–3). It is estimated that up to 25% of adults worldwide have some degree of hepatic fat accumulation (4). When fat accumulation is the extent of damage, it is well-tolerated by most people. However, in a subset of individuals, disease progression leads to liver inflammation and hepatocyte injury. Cycles of injury and repair result in measurable collagen deposition in the liver parenchyma, leading eventually to detectable fibrotic changes and, in some cases, to cirrhosis. When steatosis is followed by the tissue damage, this state is referred to as non-alcoholic steatohepatitis (NASH) (5). Formation of liver tumors, including hepatocellular carcinoma, is another common and the most devastating late-stage consequence. Why some individuals with extensive steatosis progress to more serious disease including liver cancer and others do not, remains a critical unanswered question.

The high content of saturated fat and sugar in the typical Western diet is thought to underlie fat accumulation (6–8). Experimental studies in rodents have shown that providing a high-fat or high-fat and -sugar diet leads rapidly to fatty liver infiltration – observable as early as after 8 weeks of feeding (9–11). With animals maintained for longer periods (20-25 weeks), evidence of inflammatory changes, ballooning hepatocyte degeneration and collagen deposition may also be seen. Even after 25 weeks, however, these changes are generally mild. Some studies have reported widespread liver damage (beyond steatosis) at earlier time-points but, typically, liver toxins such as carbon tetrachloride, thioacetamide or alcohol are administered along with the high-fat diet (12,13). Animals with severe genetic anomalies (14) or animals maintained on a toxic, nutrient-restricted diet such as the methionine- and choline-deficient diet (15) may develop severe liver anomalies more rapidly. When a high-fat or high-fat and -sugar diet alone is utilized in “genetically normal” strains, overt liver damage (beyond steatosis) is not routinely observed before 36 weeks of age (16) and a year or more is required for terminal damage – including hepatocellular carcinoma – to be widely manifested (17,18).

In our own previous studies (19,20), C57BL/6 mice were maintained on a high-fat Western-style diet (HFWD) for up to 18 months. The long feeding period allowed for the development of extensive liver damage in a majority of animals, especially males. In addition to wide-spread steatosis, HFWD-fed mice demonstrated extensive perivascular and parenchymal liver inflammation, ballooning hepatocyte injury and collagen deposition. Large fibrotic nodules and areas of infarct were observed in many of the animals. Liver tumors including hepatic adenomas and hepatocellular carcinomas were identified grossly and histologically in many of the animals. Of most interest, our studies (19,20) demonstrated that providing an adequate level of calcium (estimated to be 20-25 mg/day consumed) along with multiple other trace elements in a mineral supplement (Aquamin^®^) dramatically reduced tumor formation (20). Inflammation, hepatocyte injury and collagen deposition were also reduced while steatosis, itself, was largely unaffected.

While an extended feeding period is essential to see the full spectrum of liver disease develop in rodents, long-term studies are not optimal for identifying the early events that lead to steatosis and disease progression in high-fat-fed mice. Likewise, long-term studies are not ideal for understanding how dietary minerals (or other interventions) might mitigate disease progression. As a way to help understand the early events that drive liver disease progression and its prevention, cohorts of mice (MS-NASH strain) were placed on a high-fat diet with fructose added to the drinking water (high-fat diet) with and without mineral supplementation (Aquamin^®^) for a period of 16 weeks. An additional cohort of high-fat diet mice was treated with obeticholic acid (OCA), a drug that acts as a Farnesoid X Receptor (FXR) agonist, that has been shown to reduce liver damage in both humans (21–24) and mice (25). C57BL/6 mice fed a low-fat rodent chow diet served as an additional control. At the end of the in-life portion of the study, whole body weight and live weight were obtained from each animal along with liver enzyme levels and metabolic parameters. Livers were assessed histologically for steatosis and for other changes (hepatocyte degeneration, inflammation, and collagen deposition) indicative of more serious damage. Finally, liver tissue samples were evaluated using a tandem mass tag (TMT) mass spectrometry-based proteomic approach for protein expression levels in individual animals. Findings from these relatively short-term studies (especially the proteomic analyses) allow us to identify early liver changes that reflect diet differences (i.e., that distinguish low-fat and high-fat feeding) and how the effects of high-fat feeding may be modified in response to intervention. Findings from this study are described here.

## MATERIALS AND METHODS

### Experimental diets and interventions

Diet D12079B (Research Diets Inc., New Brunswick, NJ) with 5% fructose added to the drinking water was used as the high-fat Western diet for this study. This diet was designed to provide 500 gm/kg of carbohydrates, 198 gm/kg of proteins and 210 gm/kg of fat. The percentage of calories from carbohydrates, proteins and fat in this diet was 43, 17, and 40% respectively. This diet also contained 39 gm/kg of a standard mineral mix (S10001A) containing calcium carbonate and calcium phosphate and other essential trace elements. Diet 5008 (Purina) was used as the low-fat control diet. The caloric distribution from this low-fat diet was approximately 57, 27, and 16% from carbohydrates, proteins and fat, respectively. Diet 5008 was supplemented with standard minerals (2.5%) as part of the diet-formula. High-fat diet alone was considered as the control for comparison with the low-fat diet or with Aquamin^®^ and OCA as interventions in the high-fat diet.

Aquamin^®^ is a product rich in calcium, magnesium, and trace elements, obtained from the skeletal remains of red marine algae (26) (Marigot Ltd, Cork, Ireland). Aquamin^®^ contains calcium and magnesium in an approximately 14:1 ratio, along with measurable levels of 72 additional trace minerals including minerals with calcimimetic activity (for example, magnesium, strontium, and trace elements from the lanthanide family [27-29]), and essentially all of the minerals concentrated from ocean water in the algal fronds. Mineral composition was established by an independent laboratory (Advanced Laboratories; Salt Lake City, Utah) using Inductively Coupled Plasma Optical Emission Spectrometry (*ICP-OES*). Supplement Table 1 provides a complete list of elements detected in Aquamin^®^ and their relative amounts. Aquamin^®^ is sold as a dietary supplement (GRAS 000028) and is used in various products for human consumption in Europe, Asia, Australia, and North America. A single batch of Aquamin®-Soluble (citrate-malate salt) was used for this study. Aquamin^®^-Soluble was added to the drinking water (18 mg/mL) throughout the entire in-life portion of the study. We have used Aquamin^®^ in previously conducted long term mouse studies (20,30,31)

Obeticholic acid (OCA) was prepared as a 3 mg/mL suspension in 0.5% methylcellulose. Animals received 0.25 mL OCA (approximately 30 mg/kg) by oral gavage five times per week beginning on week-8 of high-fat feeding.

### Mouse model and in-life study protocol

The study was conducted in a mouse model (MS-NASH mouse, formally known as FATZO [32,33]) by Crown Bioscience, Inc. Three cohorts of male MS-NASH mice (9 animals per group) were maintained on the high-fat diet, beginning at 6-7 weeks of age (at weaning) and after one week of acclimation. One group was kept as control (high-fat diet alone) while the other groups received Aquamin^®^ or OCA as interventions along with the high-fat Western diet. Since mineral supplementation (Aquamin®) is proposed as a preventive agent, it was included along with high-fat feeding. OCA is envisioned as a therapeutic agent for existing disease, and animals were started on OCA at week-8 (16 weeks of age). A low-fat diet control group consisted of male C57BL/6 (Envigo) mice (9 animals per group) maintained on Diet 5008. Food and water were provided *ad libitum.* The animal environment was maintained at 70-74°F with a 12-hour light-dark cycle. During the in-life portion of the study, health-checks were done daily. At termination (after 16 weeks on diet), animals were euthanized by carbon dioxide asphyxiation. The entire in-life portion of the study and termination (euthanasia and tissue harvesting) was carried out by Crown Bio, Inc., at their Lafayette, LA facility under the approved Standard Operating Procedures at the testing site. All procedures involving live animals were approved by the Institutional Animal Care and Use Committee (IACUC). Crown Bio, Inc., is an AAALAC-accredited institution.

At euthanasia, blood was obtained by cardiac puncture and used to assess liver enzyme (alanine aminotransferase [ALT] and aspartate aminotransferase [AST]) levels as a measure of injury and triglyceride (TG) levels as an indicator of metabolic function. Livers were removed and weighed. After gross examination, a piece of tissue from the left lobe was immediately frozen in liquid nitrogen and used for proteomic analysis. Another tissue piece from the left lobe was fixed in 10% buffered formalin and used for histology.

### AST, ALT and triglyceride assessment

Serum levels of AST, ALT and triglycerides were assessed using a Beckman chemical analyzer. All blood chemistry measurements were made following a standard operating procedure at Crown Bioscience.

### Histology and assessment scoring

Formalin-fixed liver tissue samples were processed for histology and 3-4 μm-thick sections were stained with hematoxylin and eosin (H&E) and picrosirius red (PSR), independently. H&E-stained tissue sections were evaluated under light microscopy for evidence of steatosis, ballooning hepatocyte degeneration and inflammation using the Kleiner scoring system (34). Kleiner et al has described the standardized histological scoring assessments of these NASH-related parameters in liver sections (34). These scoring methods have previously been applied to the histological scoring of mouse models of steatohepatitis by others (35,36) and by us (19). Under this scoring system, a score of (0–3) was given for steatosis and inflammation separately, and a score of (0–2) for ballooning degeneration of hepatocytes. From these evaluations, a NAFLD activity score (NAS) was calculated by adding these individual scores. Finally, PSR-stained liver sections were used to assess collagen deposition and scored using a scale from 0–4 as defined by Kleiner et al (34).

After microscopic evaluation, slides were digitized using the Aperio AT2 whole slide scanner (Leica Biosystems) at 40x with a resolution of 0.5μm per pixel with 20x objective. The scanned images were housed on a server and accessed using Leica Aperio eSlide Manager (Version 12.3.2.5030), a digital pathology management software. These digitized histological sections were viewed and analyzed using Aperio ImageScope (Version 12.3.3.5048), a slide viewing software by Leica.

### Proteomic assessment

Proteomic experiments were carried out in the Proteomics Resource Facility (PRF), a core laboratory in the Department of Pathology at the University of Michigan. For protein mass spectrometry analysis, we employed liver tissue from five mice from each group and assessed each separately. A cryopreserved tissue piece from the left lobe (of each mouse liver) was weighed and homogenized in Radioimmuno-Precipitation Assay (RIPA) – lysis and extraction buffer (Pierce, # 89901; ThermoFisher Scientific) for protein isolation, as described in our previous publications (37–39). Fifty micrograms of sample protein from each liver were digested separately with trypsin and individual samples labeled with one of 11 isobaric mass tags following the manufacturer’s protocol. Tandem Mass Tag (TMT)11plex Isobaric Label Reagent Set (ThermoFisher Scientific) was utilized for this application. After labeling, equal amounts of sample (peptide) from each liver sample were mixed together. In order to achieve in-depth characterization of the proteome, the labeled peptides were fractionated using 2D-LC (basic pH reverse-phase separation followed by acidic pH reverse phase) and analyzed on a high-resolution, tribrid mass spectrometer (Orbitrap Fusion Tribrid, ThermoFisher Scientific) using conditions optimized at the PRF. MultiNotch MS3 analysis (40) was used to accurately quantify identified proteins/peptides. Data analysis was performed using Proteome Discoverer (v 2.4, ThermoFisher Scientific). MS2 spectra were searched against UniProt mouse protein database (17011 sequences as reviewed entries; downloaded on 05/06/2020) using the following search parameters: MS1 and MS2 tolerance were set to 10 ppm and 0.6 Da, respectively; carbamidomethylation of cysteines (57.02146 Da) and TMT labeling of lysine and N-termini of peptides (229.16293 Da) were considered static modifications; oxidation of methionine (15.9949 Da) and deamidation of asparagine and glutamine (0.98401 Da) were considered variable. Identified proteins and peptides were filtered to retain only those that passed ≤2% false discovery rate (FDR) threshold of detection. Quantitation was performed using high-quality MS3 spectra (Average signal-to-noise ratio of 10 and <40% isolation interference). Differential protein expression in each treatment cohort was normalized to the high-fat alone cohort. Protein names retrieved from UniProt.org, and Reactome V78 (reactome.org) was used for pathway enrichment analyses (41) for species *Mus musculus.* STRING database-v11 (string-db.org) was used to detect protein-protein interactions and additional enrichment analyses provide information related to cellular components, molecular functions, and biological processes by Gene Ontology (GO) annotation. It also offered WikiPathways and Kyoto Encyclopedia of Genes and Genomes (KEGG) databases to curate pathways. Only proteins with a ≤2% FDR confidence of detection were included in the analyses. The individual differential protein expression was established by calculating the abundance ratio of normalized abundances of each intervention sample to the high-fat alone samples. For the group analysis, individual data points were merged by groups to obtain means and standard deviations. The initial analysis was an unbiased search of differentially-expressed proteins and significantly altered proteins. Subsequently, we searched the data base, specifically, for changes in the expression of keratins and other differentiation-related proteins. Mass spectrometry-based proteomics data were placed in a data repository known as ProteomeXchange Consortium via the PRIDE partner repository with the dataset identifier PXD030954.

Statistical analysis. Group means and standard deviations were obtained for discrete gross and histochemical features as well as for individual biochemical values (ALT, AST and TG) and individual protein values obtained in the proteomic analysis. Data generated in this way were analyzed by analysis of variance (ANOVA) followed by unpaired t-test (two-tailed) for comparison using GraphPad Prism, version 9. Pathways enrichment data reflect Reactome-generated *p*-values based on the number of entities identified in a given pathway as compared to total proteins responsible for that pathway. The significance (*p*-value) is calculated by the overrepresentation analysis (hypergeometric distribution). For STRING enrichment analysis, the whole genome statistical background was assumed and FDR stringency was ≤1%. Data were considered significant at *p*<0.05.

## RESULTS

### Body weight, liver weight comparisons, liver enzyme levels and triglycerides

Animal weights and liver weights were assessed at euthanasia (Figure 1A). As expected, there were large differences between mice fed the low-fat diet and those on the high-fat diet irrespective of intervention. Inclusion of Aquamin^®^ along with high-fat feeding had no effect on either total body weight or liver weight, but there was a small reduction in both parameters in the cohort of mice receiving OCA as compared to control mice on the high-fat diet. Liver enzyme (ALT and AST) levels and triglyceride levels are shown in Figure 1B. Levels of both ALT and AST increased substantially with high-fat feeding as did triglyceride levels. Both liver enzymes were also increased in mice receiving either Aquamin^®^ or OCA along with high-fat feeding. With OCA, there was a modest reduction compared to levels seen in untreated mice on the same diet. Differences between the interventions and control (high-fat diet-fed mice) were not statistically different for either ALT or AST but all three high-fat groups were different from the low-fat control group. Serum triglyceride levels (evidence of metabolic dysfunction) were significantly elevated with high-fat feeding but neither Aquamin^®^ nor OCA had a significant effect on this parameter.

**Figure 1.**
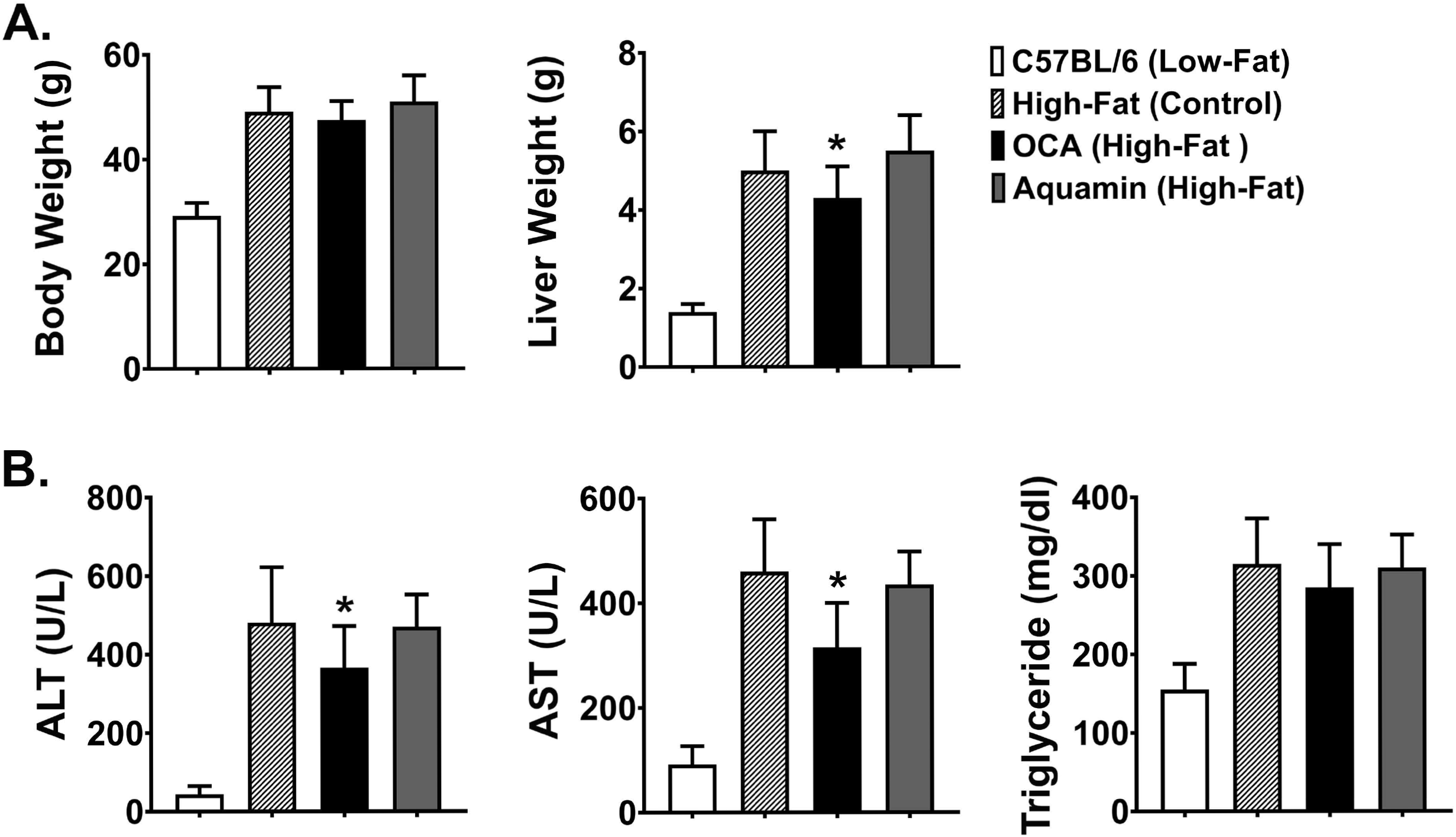
Whole body weight, liver weight and biochemical markers. **A**. Whole body weight and liver weight were obtained for each animal at euthanasia. **B**. Blood was obtained by cardiac puncture at euthanasia and assessed for ALT, AST and triglyceride levels. Values shown are means and standard deviations based on 9 animals per group. Statistical significance was assessed by ANOVA followed by two-tailed t-test. Asterisks (*) indicate statistical difference from high-fat control at p<0.05. With all 5 parameters, low-fat control values were statistically different from high-fat control at p<0.05.

### Histological features

Sections of formalin-fixed liver tissue were stained with hematoxylin and eosin and evaluated for steatosis, ballooning hepatocyte degeneration and inflammation. Additional sections were stained with PSR and assessed for collagen deposition. Data are shown in Figure 2. Steatosis was virtually undetectable in mice on the low-fat control diet but widespread in every animal on the high-fat diet after 16 weeks of feeding. Neither Aquamin^®^ nor OCA reduced steatosis significantly compared to the high-fat control, but both interventions reduced ballooning degeneration and inflammation to some extent. When scores from the three parameters were added together (as in citation 34) to give a NAFLD activity score, both Aquamin^®^ and OCA were different from high-fat alone. However, the degree of elevation was small to begin with (especially with inflammation) and the overall significance of the modest reduction is difficult to gauge. PSR staining demonstrated collagen deposition within the liver parenchyma of mice on the high-fat diet while sections from low-fat control mice did not show staining except around vascular elements and in relation to the capsule. Some reduction in the collagen deposition score was noted with both interventions. With Aquamin^®^, the reduction in the collagen deposition reached a level of statistical significance. However, as with inflammation, the overall PSR staining score was low (<1.5 out of 4) after 16 weeks on the high-fat diet.

**Figure 2.**
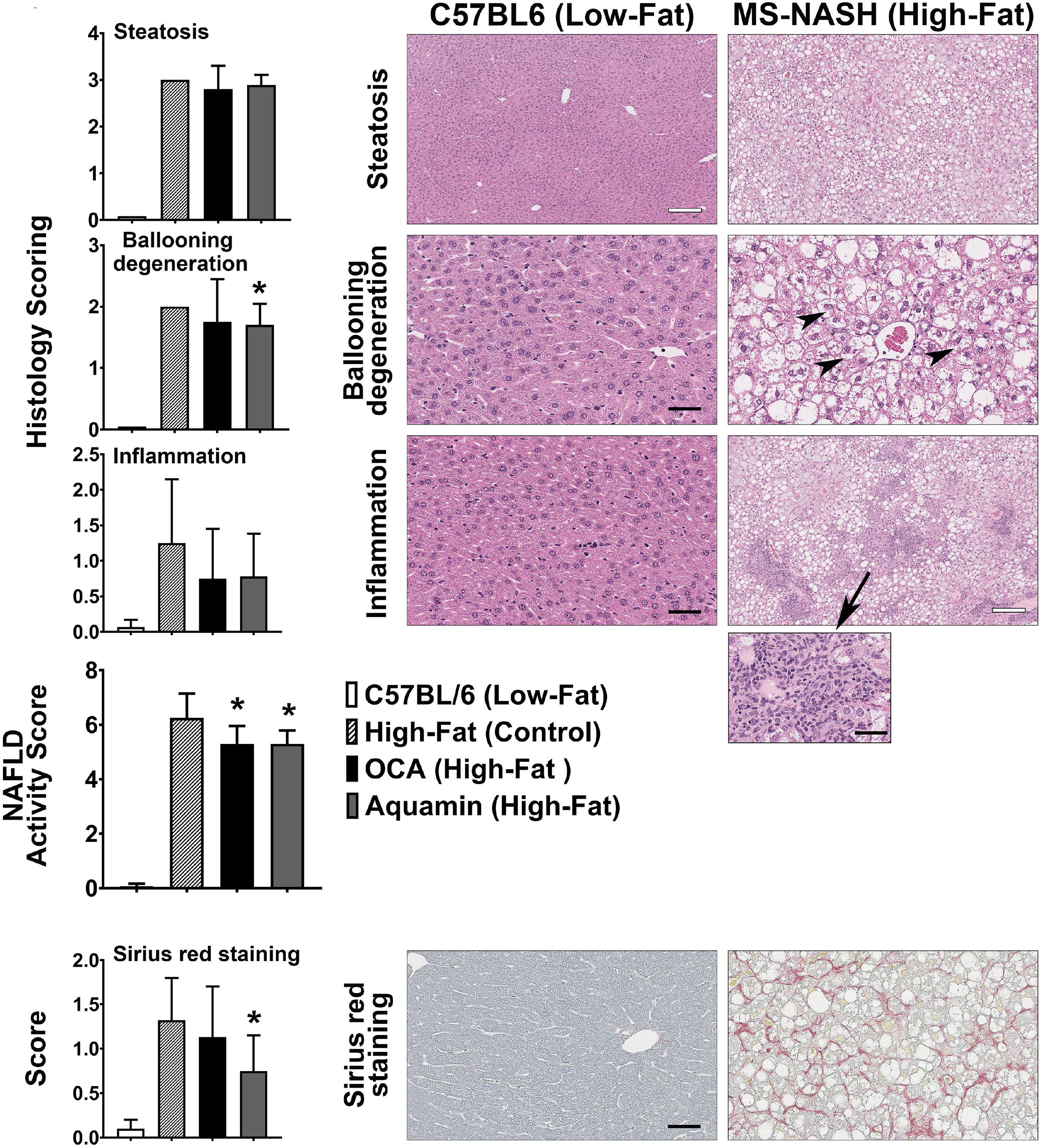
Histological features. H&E-stained sections were evaluated for steatosis, ballooning hepatocyte degeneration and inflammation by two trained individuals who did not know the treatment group from which the section was obtained. Scoring of each parameter was as described in the Materials and Methods Section and followed the Kleiner system (34). The NAFLD activity score (NAS) is a summation of the three individual parameters (steatosis, ballooning hepatocyte degeneration and inflammation), calculated in each mouse. Picrosirius red (PSR)-stained sections were evaluated for collagen deposition. Scoring of each parameter was as described in the Materials and Methods Section. Values shown are means and standard deviations based on 9 animals per group. Statistical significance was assessed by ANOVA followed by two-tailed t-test. Asterisks (*) indicate statistical difference from high-fat control at p<0.05. For steatosis panels, bar =200 μm, for ballooning degeneration and PSR panels, bar=50 μm and for inflammation in low-fat, bar=50 μm, and in high-fat, bar=200 μm. For inflammation in high-fat, a small panel is showing an inflammatory lesion at a higher resolution, bar=50 μm.

### Proteomic analysis: Effects of low-fat diet, Aquamin^®^ and OCA compared to high-fat alone

The number of upregulated and downregulated proteins in C57BL/6 mice on the low-fat diet (relative to high-fat control mice) and the number of up- and downregulated proteins with Aquamin^®^ or OCA in comparison to the high-fat control mice is presented as Figure 3A (2-fold cutoff) and 3B (1.5-fold cutoff). It can be seen from the bar graphs that the low-fat control diet was responsible for a substantial number of protein changes. With the 2-fold cutoff and <2% FDR, 182 proteins were upregulated, and 126 proteins were downregulated with the low-fat control diet. With Aquamin^®^ and OCA, a total of 86 and 76 proteins, respectively, were upregulated to the same extent, while only five and eight, respectively, were downregulated. When the less stringent cut-off (1.5-fold) was used, more proteins were altered with each intervention but the trends were the same.

**Figure 3.**
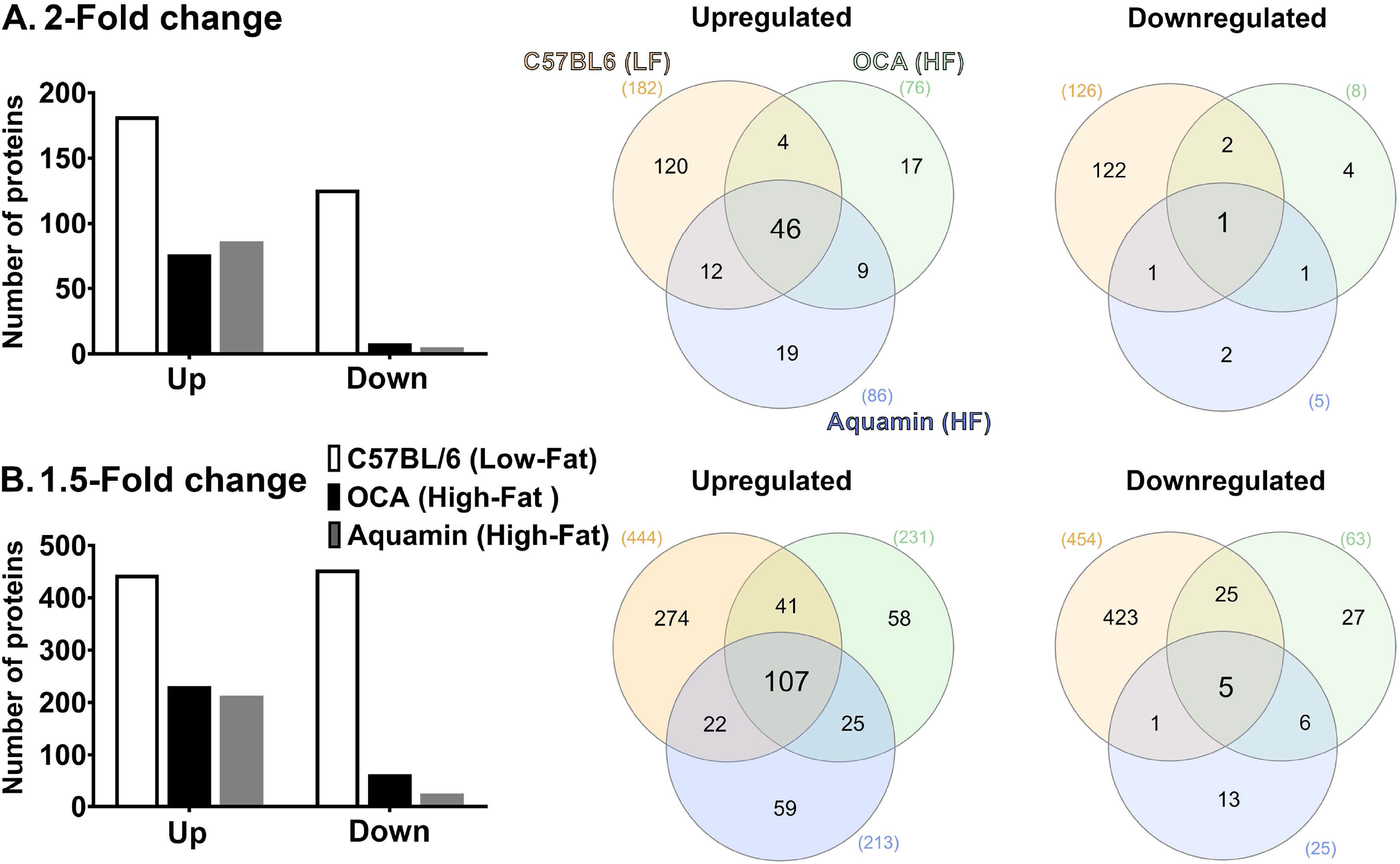
Proteomic analysis. Lysates were prepared from liver tissue of individual mice (5 per group) and subjected to Tandem Mass Tag (TMT) labeling / mass spectroscopy as described in the Materials and Methods Section. Data from individual animals were merged by treatment group for analysis. **Left panels**: Abundance ratios of all proteins were calculated by comparing the normalized abundance of each group to the high-fat control group. The upper panel indicates the number of proteins that was increased or decreased by 2-fold (**A**) or greater (with <2% FDR) and the lower panel indicates the number of proteins that was increased or decreased by 1.5-fold (**B**) or greater (with <2% FDR). **Right panels**: Venn diagrams showing the overlap in the number of proteins altered (increased or decreased) by an average of 2-fold (**A**) or greater (upper panel) or 1.5-fold (**B**) or greater (lower panel) among the three treatment groups relative to the high-fat control and unique to each group.

Venn diagrams (Figure 3A and 3B) show the overlap between protein alterations with low-fat feeding and the other two interventions. Among the proteins elevated with either Aquamin^®^ or OCA, approximately 50% (at either cut-off) were in common with proteins upregulated in the low-fat control diet. In contrast, the number of downregulated proteins with either intervention that overlapped with proteins downregulated with low-fat feeding was lower on a percentage basis. This was seen with either cutoff but was more extreme with 2-fold cutoff. Overall, 57 upregulated proteins were common between the two interventions (OCA and Aquamin^®^), while only two downregulated proteins were common between these two groups. Supplement Table 2 and Supplement Table 3 provide lists of upregulated and downregulated proteins common at 2-fold among three groups (low-fat, OCA and Aquamin^®^ on high-fat diet) across five mice per group.

### Proteomic analysis: Individual protein changes in response to Aquamin^®^

Figure 4 shows the foldchange distribution of individual proteins responsive to Aquamin^®^ in the presence of high-fat diet. Upregulated proteins that met the criterion of statistical significance at p<0.05 are shown in blue while downregulated proteins are shown in red. The bold colors (blue and red) indicate proteins that were statistically different and also at least 1.5-fold up- or downregulated.

**Figure 4.**
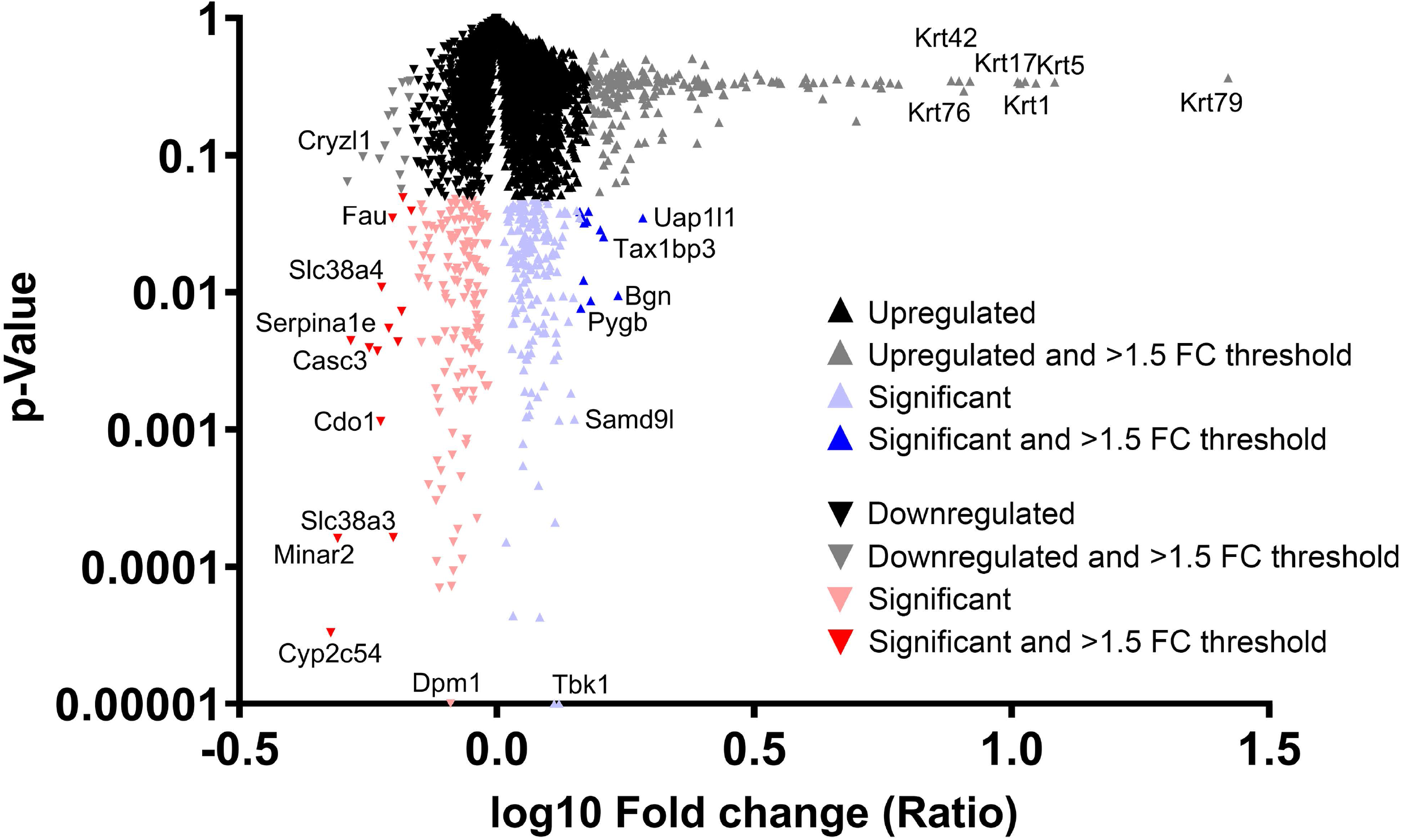
Distribution of proteins upregulated or downregulated in response to Aquamin^®^ compared to high-fat control. Values for each protein (group average) in the high-fat control was set to 1.0 and values in the Aquamin^®^ group compared to this. The x-axis shows the fold-change (log10) of individual proteins and the y-axis reflects individual protein *p*-values (N=5 individual liver samples per group). The mass of proteins shown in black represent proteins that were less than 1.5-fold different from control (up or down) and not statistically significant. Proteins depicted with gray were different from control by 1.5-fold or greater but not statistically different. Red triangles represent down-regulated proteins and blue triangles represent up-regulated proteins that were statistically significant as compared to control. Bold color (red or blue) represents proteins that were both statistically different from control and different from control by 1.5-fold or greater. Certain individual proteins are identified by UniProt protein (gene) symbol.

Table 1A identifies the upregulated proteins that met the dual criteria (statistical difference from the high-fat control at p<0.05 and at least 1.5-fold increase) in response to Aquamin®. Supplement Table 4 provide a comprehensive list of upregulated proteins that met the criterion of statistical difference from control, irrespective of fold-change. While only a small number of proteins (12 in total) met both criteria, a total of 183 upregulated proteins were statistically increased. Reactome pathway analysis was utilized as a way to help delineate how altered protein expression in response to Aquamin^®^ influences biological function. As can be seen from the altered pathways, the upregulated proteins impact lipid metabolism and trafficking of lipids (Table 1B). Supplement Table 5 lists top pathways affected by the altered (upregulated) proteins presented in Supplement Table 4.

**Table 1A.**
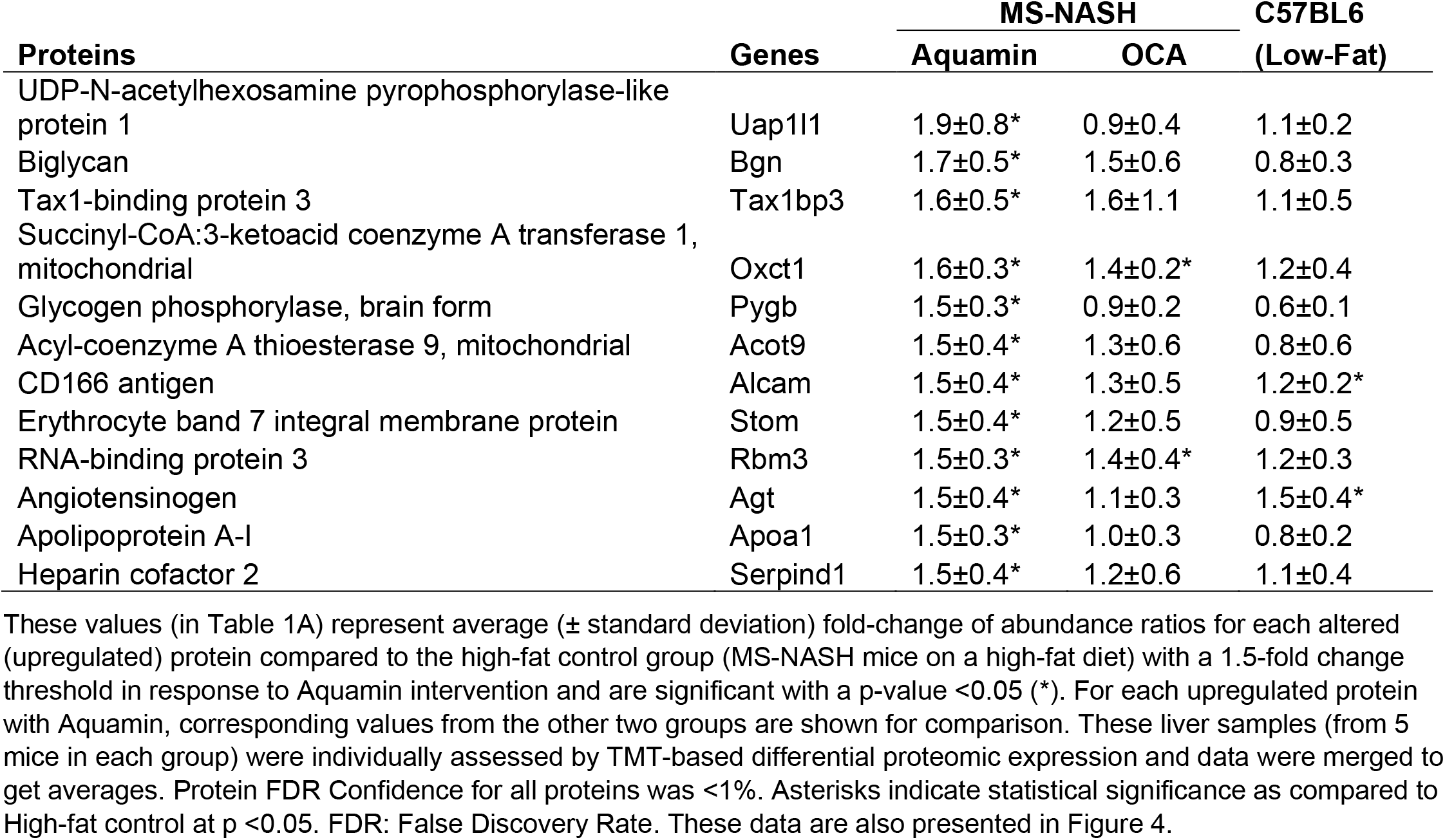
Significantly upregulated proteins with Aquamin in high-fat mice.

**Table 1B.**
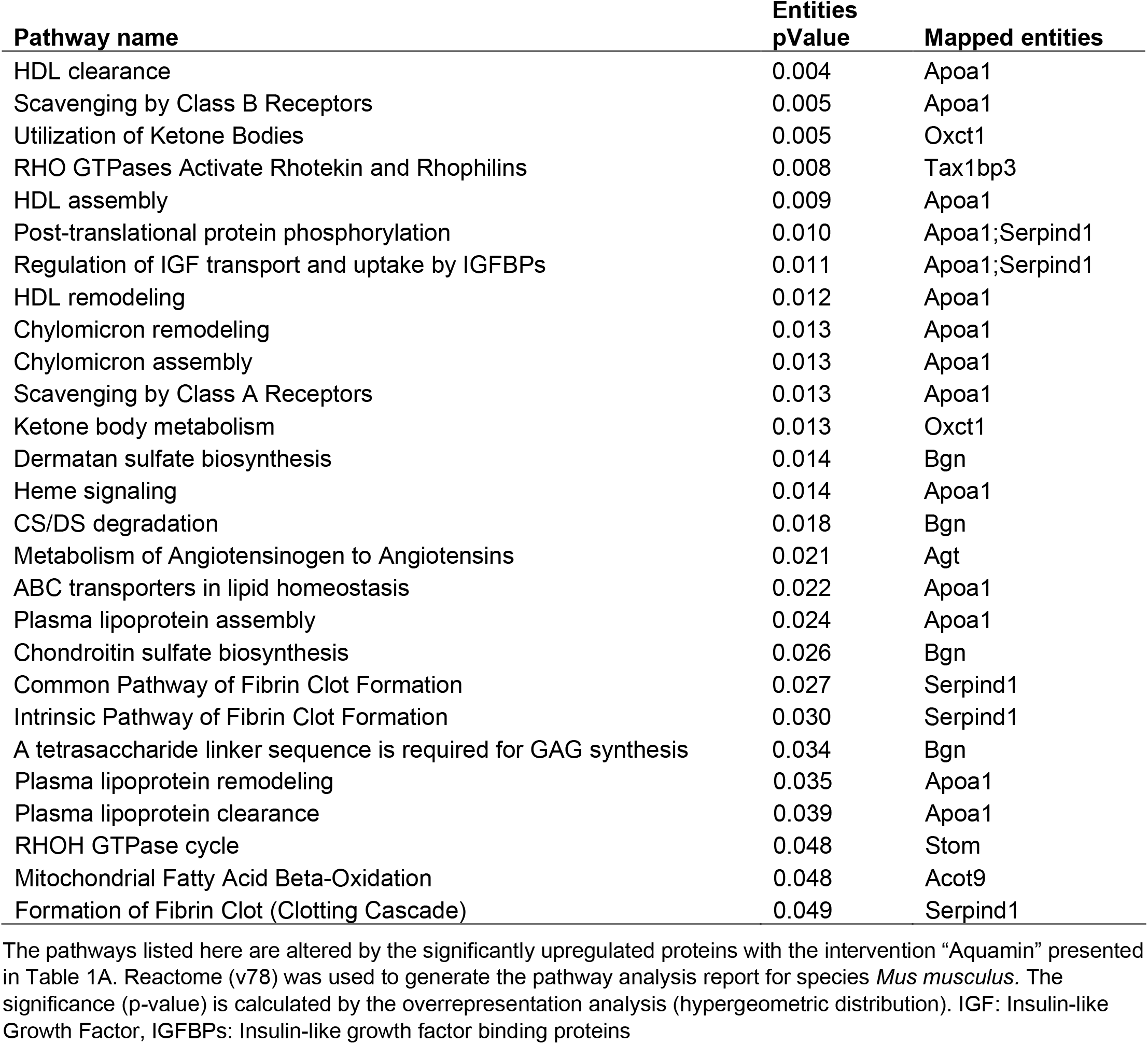
Top pathways associated with upregulated proteins (listed in 1A) altered with Aquamin.

| Pathway name | Entities | Mapped entities |
| --- | --- | --- |
|  | pValue |  |
| HDL clearance | 0.004 | Apoa1 |
| Scavenging by Class B Receptors | 0.005 | Apoa1 |
| Utilization of Ketone Bodies | 0.005 | Oxct1 |
| RHO GTPases Activate Rhotekin and Rhophilins | 0.008 | Tax1bp3 |
| HDL assembly | 0.009 | Apoa1 |
| Post-translational protein phosphorylation | 0.010 | Apoa1;Serpind1 |
| Regulation of IGF transport and uptake by IGFBPs | 0.011 | Apoa1;Serpind1 |
| HDL remodeling | 0.012 | Apoa1 |
| Chylomicron remodeling | 0.013 | Apoa1 |
| Chylomicron assembly | 0.013 | Apoa1 |
| Scavenging by Class A Receptors | 0.013 | Apoa1 |
| Ketone body metabolism | 0.013 | Oxct1 |
| Dermatan sulfate biosynthesis | 0.014 | Bgn |
| Heme signaling | 0.014 | Apoa1 |
| CS/DS degradation | 0.018 | Bgn |
| Metabolism of Angiotensinogen to Angiotensins | 0.021 | Agt |
| ABC transporters in lipid homeostasis | 0.022 | Apoa1 |
| Plasma lipoprotein assembly | 0.024 | Apoa1 |
| Chondroitin sulfate biosynthesis | 0.026 | Bgn |
| Common Pathway of Fibrin Clot Formation | 0.027 | Serpind1 |
| Intrinsic Pathway of Fibrin Clot Formation | 0.030 | Serpind1 |
| A tetrasaccharide linker sequence is required for GAG synthesis | 0.034 | Bgn |
| Plasma lipoprotein remodeling | 0.035 | Apoa1 |
| Plasma lipoprotein clearance | 0.039 | Apoa1 |
| RHOH GTPase cycle | 0.048 | Stom |
| Mitochondrial Fatty Acid Beta-Oxidation | 0.048 | Acot9 |
| Formation of Fibrin Clot (Clotting Cascade) | 0.049 | Serpind1 |

Aquamin®-responsive proteins the met the dual criteria of 1.5-fold decrease and statistical significance are shown in Table 1C and those that were statistically-different (and downregulated) from high-fat control, regardless of fold change, are shown in Supplement Table 6. A total of 14 proteins met both criteria while a total of 130 were statistically different from the high-fat control, independent of foldchange. As with upregulated proteins, Reactome pathway analysis was used to help identify potential mechanisms of action (Table 1D). Proteins whose expression was reduced by Aquamin^®^ impact, primarily, amino acid metabolism, although purine and pyrimidine metabolism also appear to be targets. Among the downregulated proteins were multiple cytochrome P450 family members (7 in all) including three (i.e., 2C54, 2C50, 2C29) that were among the most highly downregulated. Cytochrome P450 family members are highly-expressed in liver and serve to metabolize and detoxify numerous chemicals, including fats (42). Supplement Table 7 lists top pathways affected by all of the significantly downregulated proteins presented in Supplement Table 6. Amino acid metabolism, bile acid and bile salt metabolism and recycling of bile acids and salts are among the significantly downregulated pathways identified.

**Table 1C.**
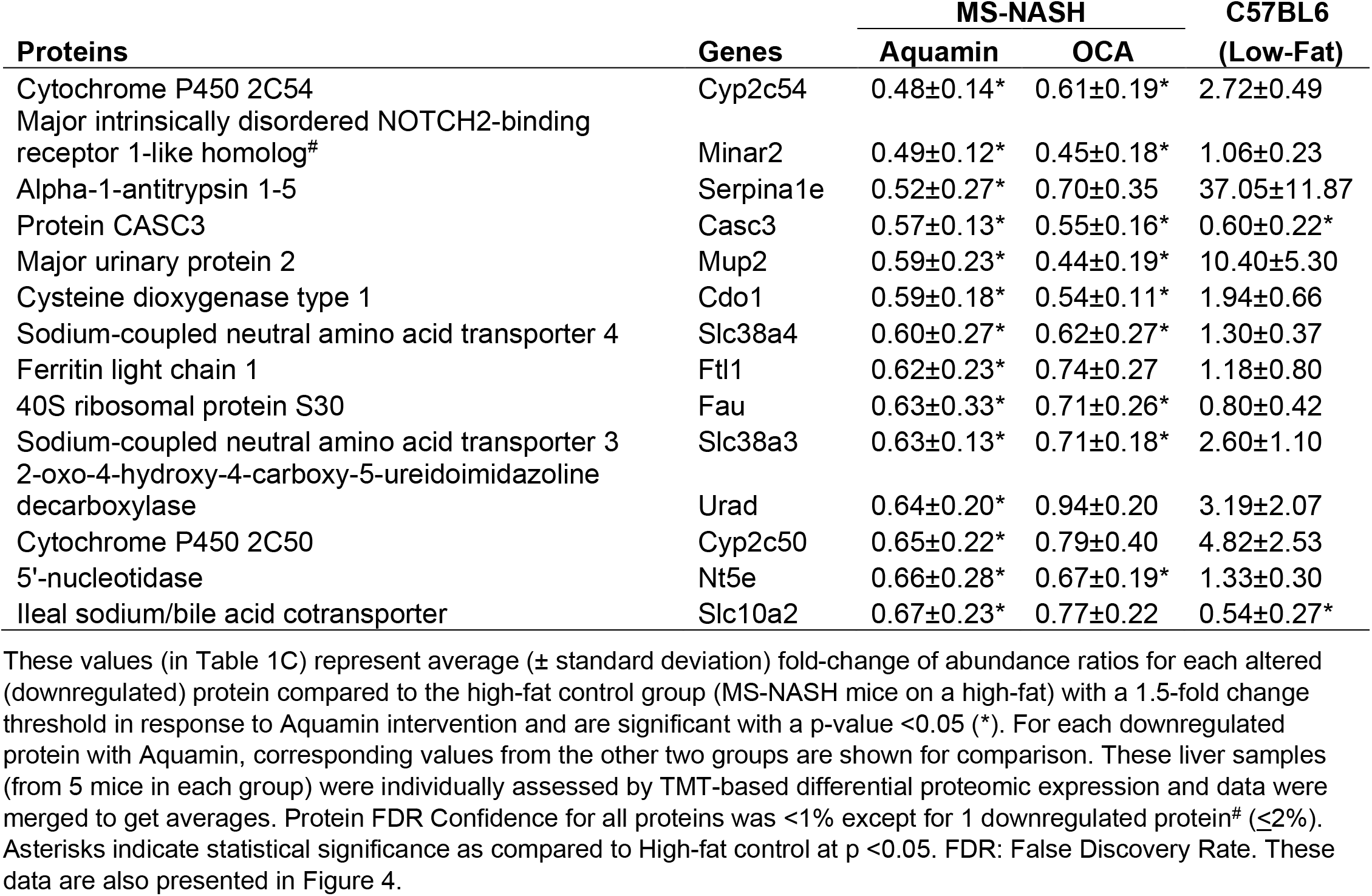
Significantly downregulated proteins with Aquamin in high-fat mice.

**Table 1D.**
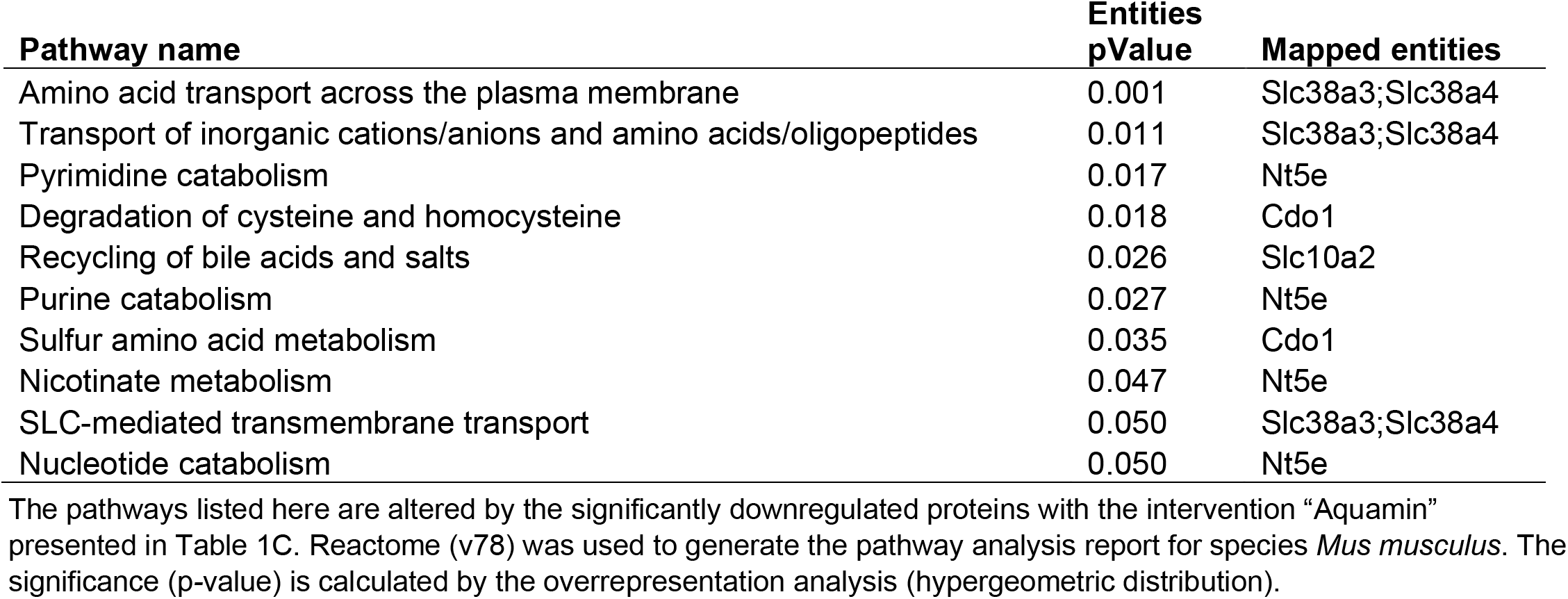
Top pathways associated with downregulated proteins (listed in 1C) altered with Aquamin.

| Pathway name | Entities<br>pValue | Mapped entities |
| --- | --- | --- |
| Amino acid transport across the plasma membrane | 0.001 | Slc38a3;Slc38a4 |
| Transport of inorganic cations/anions and amino acids/oligopeptides | 0.011 | Slc38a3;Slc38a4 |
| Pyrimidine catabolism | 0.017 | Nt5e |
| Degradation of cysteine and homocysteine | 0.018 | Cdo1 |
| Recycling of bile acids and salts | 0.026 | Slc10a2 |
| Purine catabolism | 0.027 | Nt5e |
| Sulfur amino acid metabolism | 0.035 | Cdo1 |
| Nicotinate metabolism | 0.047 | Nt5e |
| SLC-mediated transmembrane transport | 0.050 | Slc38a3;Slc38a4 |
| Nucleotide catabolism | 0.050 | Nt5e |

Tables 1A and 1C also show for comparison how the Aquamin®-sensitive proteins responded to low-fat feeding and to OCA. It can be seen that while there was overlap between Aquamin^®^ and OCA (as shown in Figure 3) in the many of the highly upregulated proteins, the response to OCA was not as robust as the response to Aquamin^®^ (i.e., fold-change was smaller and only two were statistically different from control). In regard to downregulated moieties, of the 14 Aquamin®-sensitive proteins that met the dual criteria, nine were also responsive to OCA by the same criteria. In contrast, there was little overlap between Aquamin®-responsiveness and response to the low-fat diet. In fact, many of the protein changes were in the opposite direction (Tables 1A and 1C).

Supplement Tables 8 and 9 present proteomic enrichment data curated by the STRING database with 183 significantly upregulated proteins and 130 significantly downregulated proteins, respectively, in response to Aquamin®. These datasheets provide information related to biological process, molecular function and cellular components along with additional pathways curated by KEGG and Wiki (as part of STRING enrichment analysis).

After analyzing data based on statistical significance, the proteomic database was searched for the most highly up- and downregulated Aquamin^®^-responsive proteins based on fold-change independent of statistical significance. Table 2 presents a list of the most highly upregulated proteins (2-fold or greater) with Aquamin^®^. As a group, epithelial cell keratins were prominent in the protein list. Of the top 22 most highly upregulated proteins, 10 were keratins (Table 2). Also, nine of these keratins were common at 2-fold-change with the low-fat group (Supplement Table 2). Of interest, the major keratins found in the liver – i.e., keratin 8, and 18 (43) – were not altered (up or down) with Aquamin^®^ (Figure 5). Keratins are structural components of epithelial cells and play key roles in differentiation, morphogenesis, barrier formation and tissue integrity. They are also important signaling intermediates (44). How each of the keratins identified here functions in the liver, specifically, is not known. The effects of low-fat feeding and the effects of treatment with OCA on these same proteins are included in Table 2 (complete protein list) and Figure 5 (keratins). As can be seen, some of the same keratins were increased with low-fat feeding but the degree of upregulation was less than that observed with Aquamin®. While the low-fat diet does not provide the same trace elements as does Aquamin®, D5008 chow contains a mineral supplement as part of its formulation. Thus, it is not unreasonable to observe some change in the keratin expression profile in livers of mice on the low-fat diet. In contrast, with the exception of keratin 79, upregulation of individual keratins with OCA was almost non-existent.

**Table 2.**
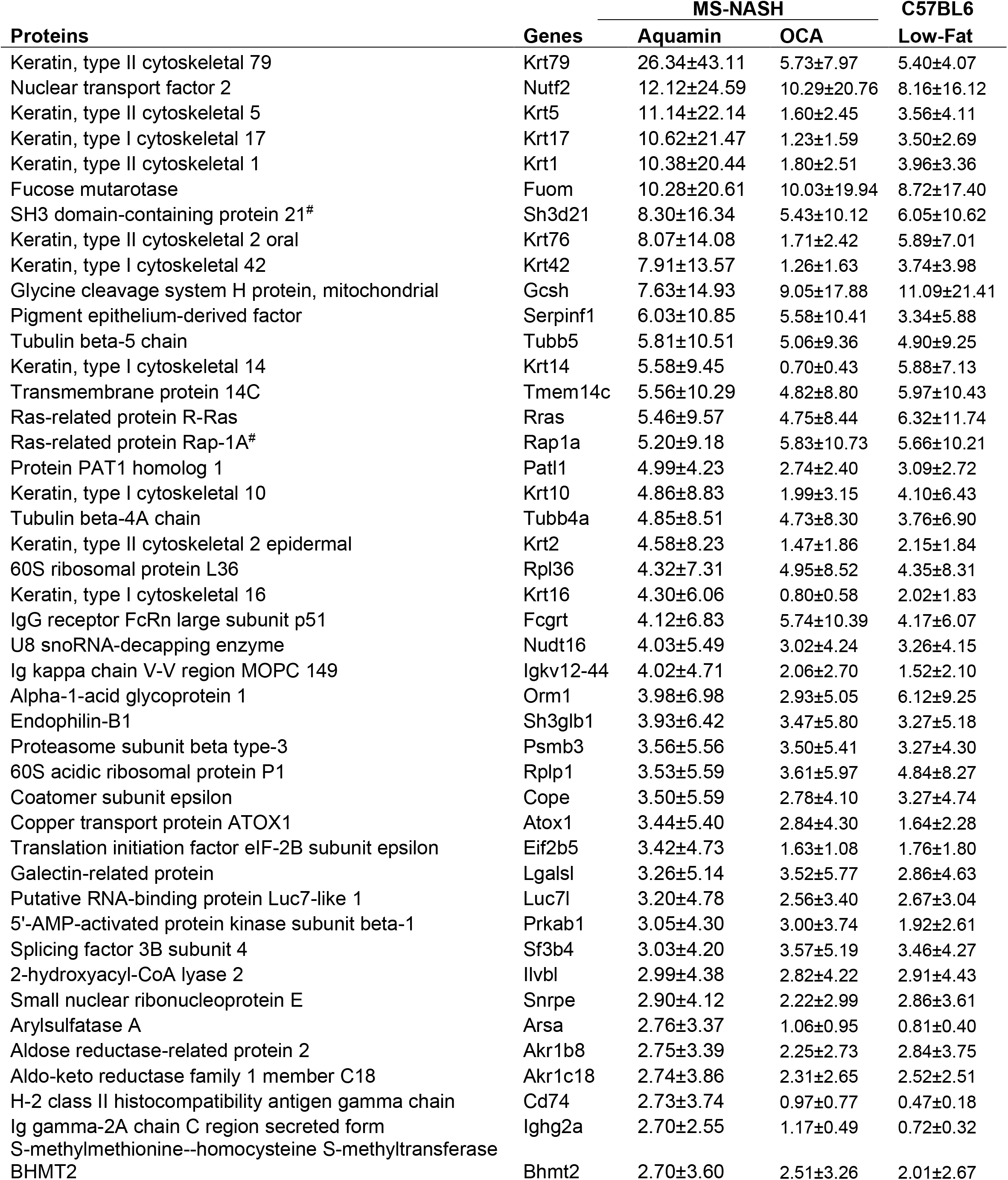
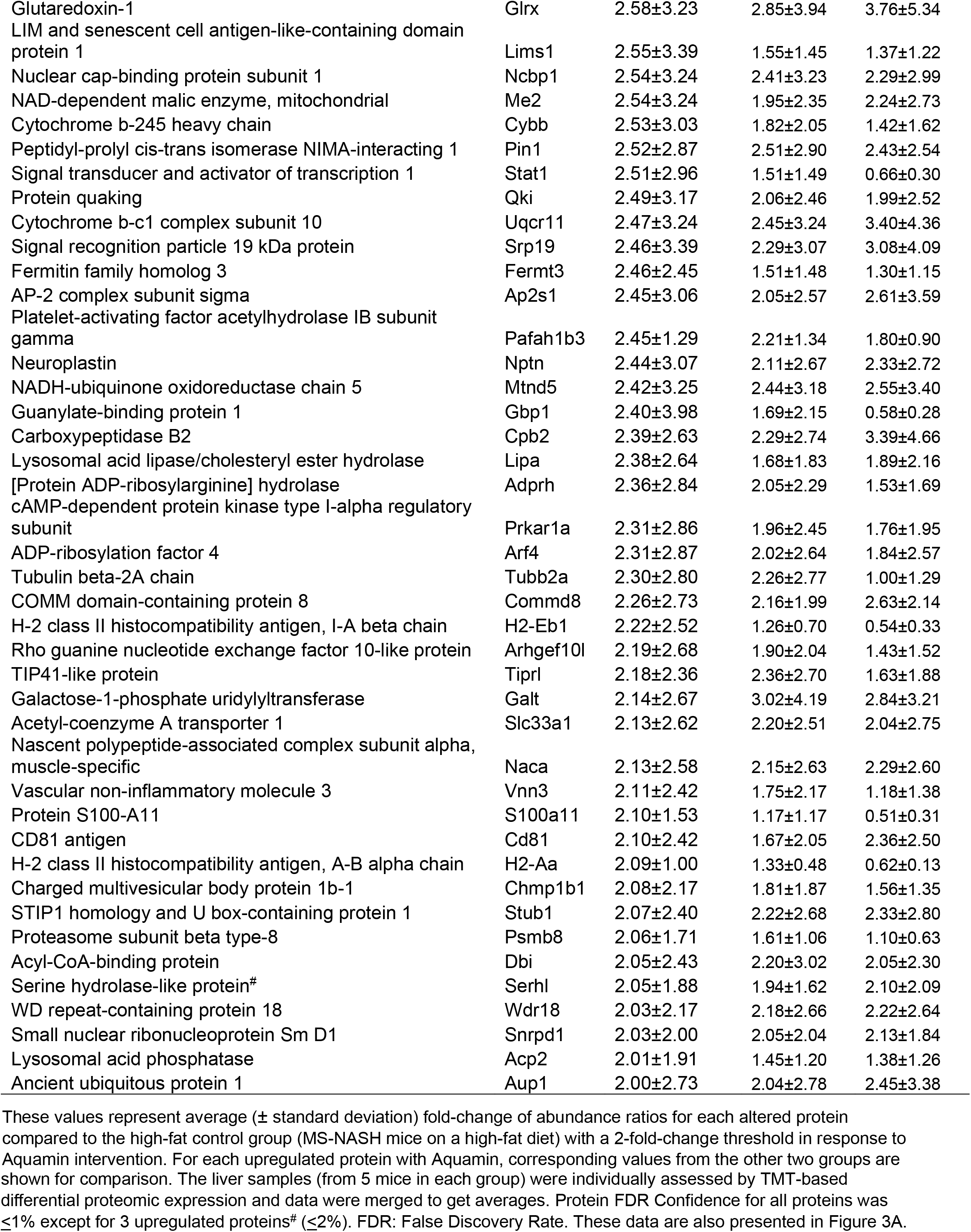
Upregulated proteins by an unbiased proteomic screening with Aquamin in high-fat mice.

**Figure 5.**
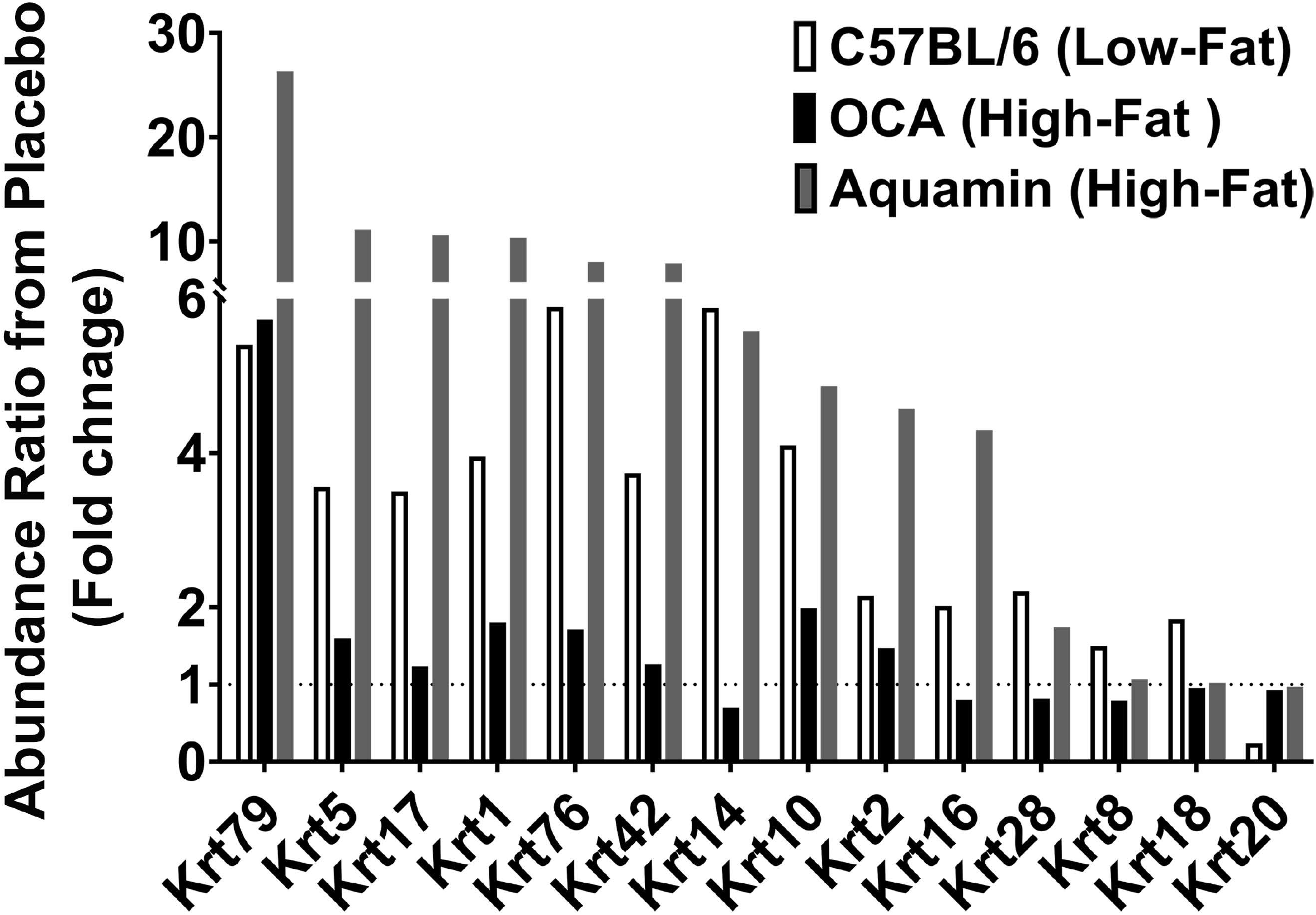
Keratins upregulated in response to Aquamin®. Values represent average fold-change from high-fat control based on n=5 liver samples per treatment group. Corresponding values for response to low-fat diet and response to OCA are shown for comparison.

In addition to keratins, three members of the tubulin family were also upregulated in response to Aquamin^®^ (Table 2). Tubulins constitute another set of important structural proteins within cells, as they are the major constituents of the microtubule system of the cytoskeleton (45). Of interest, the tubulin proteins were responsive to both low-fat feeding and OCA treatment as well as to Aquamin^®^ (Table 2).

Reactome pathway analysis was utilized to help identify the various pathways influenced by the protein changes noted in Table 2 with Aquamin®. Not surprisingly, pathways related to differentiation were the most highly affected (Table 3). Among these were keratinization, cornified envelope formation, hemidesmosome assembly, gap junction formation and gap junction assembly. In addition, pathways involved in intracellular trafficking of proteins as well as protein trafficking between the cytoplasm and plasma membrane were significantly influenced by the protein changes occurring in response to Aquamin®. Protein trafficking events are well-known to depend on cations in the environment (46,47). Another pathway of interest was *hedgehog signaling in the off-state.* Signaling through the hedgehog pathway is associated with chronic liver injury (48,49) and hedgehog signaling may have implications in hepatocellular carcinomas. All of the Aquamin®-responsive Reactome pathways that met the criterial of having an entities *p* value of <0.05 are presented in Table 3.

**Table 3.** Top pathways associated with upregulated proteins altered (at 2-fold change) with Aquamin.

| Pathway name | Entities<br>pValue | Mapped entities |
| --- | --- | --- |
| Formation of the cornified envelope | $1.7 \times 10^{-7}$ | Krt2;Krt1;Krt79;Krt16;Krt5;Krt76;Krt17;Krt10;Krt14 |
| Keratinization | $3.8 \times 10^{-5}$ | Krt2;Krt1;Krt79;Krt16;Krt5;Krt76;Krt17;Krt10;Krt14 |
| Developmental Biology | $8.5 \times 10^{-4}$ | Krt1;Krt16;Rras;Krt76;Krt14;Tubb2a;Krt2;Krt79;<br>Krt5;Tubb4a;Krt17;Krt10;Ap2s1 |
| mRNA Splicing - Minor Pathway | 0.002 | Sf3b4;Ncbp1;Snrpe;Snrpd1 |
| Hedgehog 'off' state | 0.004 | Prkar1a;Tubb4a;Psmb8;Psmb3;Tubb2a |
| SLBP independent Processing of Histone Pre-mRNAs | 0.004 | Ncbp1;Snrpe |
| Recycling pathway of L1 | 0.01 | Tubb4a;Tubb2a;Ap2s1 |
| SLBP Dependent Processing of Replication-Dependent Histone Pre-mRNAs | 0.01 | Ncbp1;Snrpe |
| Type I hemidesmosome assembly | 0.01 | Krt5;Krt14 |
| The role of GTSE1 in G2/M progression after G2 checkpoint | 0.01 | Tubb4a;Psmb8;Psmb3;Tubb2a |
| Golgi-to-ER retrograde transport | 0.01 | Pafah1b3;Arf4;Tubb4a;Cope;Tubb2a |
| Microtubule-dependent trafficking of connexons from Golgi to the plasma membrane | 0.01 | Tubb4a;Tubb2a |
| Transport of connexons to the plasma membrane | 0.01 | Tubb4a;Tubb2a |
| Signaling by Hedgehog | 0.01 | Prkar1a;Tubb4a;Psmb8;Psmb3;Tubb2a |
| COPI-independent Golgi-to-ER retrograde traffic | 0.01 | Pafah1b3;Tubb4a;Tubb2a |
| Regulation of RUNX2 expression and activity | 0.01 | Stub1;Psmb8;Psmb3 |
| COPI-mediated anterograde transport | 0.01 | Arf4;Tubb4a;Cope;Tubb2a |
| COPI-dependent Golgi-to-ER retrograde traffic | 0.01 | Arf4;Tubb4a;Cope;Tubb2a |
| LDL clearance | 0.01 | Lipa;Ap2s1 |
| Transcriptional regulation by RUNX2 | 0.02 | Stub1;Psmb8;Psmb3 |
| RHO GTPases Activate NADPH Oxidases | 0.02 | Pin1;Cybb |
| MHC class II antigen presentation | 0.02 | Tubb4a;Cd74;Tubb2a;Ap2s1 |
| Carboxyterminal post-translational modifications of tubulin | 0.03 | Tubb4a;Tubb2a |
| Sealing of the nuclear envelope (NE) by ESCRT-III | 0.03 | Tubb4a;Tubb2a |
| Regulation of PTEN stability and activity | 0.03 | Stub1;Psmb8;Psmb3 |
| L1CAM interactions | 0.03 | Tubb4a;Tubb2a;Ap2s1 |
| Cell junction organization | 0.03 | Lims1;Krt5;Krt14 |
| G2/M Transition | 0.03 | Tubb5;Tubb4a;Psmb8;Psmb3;Tubb2a |
| Processing of Capped Intronless Pre-mRNA | 0.03 | Ncbp1;Snrpe |
| Mitotic G2-G2/M phases | 0.03 | Tubb5;Tubb4a;Psmb8;Psmb3;Tubb2a |
| Intra-Golgi and retrograde Golgi-to-ER traffic | 0.03 | Pafah1b3;Arf4;Tubb4a;Cope;Tubb2a |
| Plasma lipoprotein clearance | 0.03 | Lipa;Ap2s1 |
| Gap junction assembly | 0.04 | Tubb4a;Tubb2a |
| Glycine degradation | 0.04 | Gcsh |
| Ion influx/efflux at host-pathogen interface | 0.04 | Atox1 |
| Metabolism of RNA | 0.04 | Sf3b4;Ncbp1;Snrpe;Psmb8;Psmb3;Rplp1;<br>Patl1;Rpl36;Snrpd1;Wdr18 |
| Aggrephagy | 0.04 | Tubb4a;Tubb2a |
| ARMS-mediated activation | 0.05 | Rap1a |
| Galactose catabolism | 0.05 | Galt |
| PCP/CE pathway | 0.05 | Psmb8;Psmb3;Ap2s1 |
| ER to Golgi Anterograde Transport | 0.05 | Arf4;Tubb4a;Cope;Tubb2a |
| Recruitment of NuMA to mitotic centrosomes | 0.05 | Tubb5;Tubb4a;Tubb2a |
| SRP-dependent cotranslational protein targeting to membrane | 0.05 | Srp19;Rplp1;Rpl36 |
| CREB1 phosphorylation through the activation of Adenylate Cyclase | 0.06 | Prkar1a |
| PKA activation in glucagon signalling | 0.06 | Prkar1a |
| GDP-fucose biosynthesis | 0.06 | Fuom |
| Cell-Cell communication | 0.06 | Lims1;Krt5;Krt14 |
| Gap junction trafficking | 0.06 | Tubb4a;Tubb2a |
| Gap junction trafficking and regulation | 0.06 | Tubb4a;Tubb2a |

The database was also searched for proteins that were downregulated by 2-fold or greater with Aquamin^®^. A total of five individual proteins met this criterion. Of the proteins in the list, four of the five were also statistically significant and presented as part of Table 1.

### Proteomic changes in response to low-fat feeding and to OCA

While the response to Aquamin^®^ was the focus of this study, we also assessed protein changes driven by low-fat feeding or by OCA in high-fat mice. Supplement Tables 10 and 11 show proteins upregulated and downregulated, respectively, by 2-fold or greater with low-fat feeding. Reactome pathways influenced by these protein changes are shown in Supplement Table 12 and 13. A large majority of the protein changes observed with low-fat feeding (both up- and downregulated) are related to fat and carbohydrate metabolism. This is consistent with previous proteomic assessment of diet-influenced protein profile in livers of both rats (50) and mice (51).

Supplement Tables 14 and 15 show protein changes in mice on high-fat diet treated with OCA. Supplement Tables 16 and 17 show pathways responsive to OCA. The most interesting (if not entirely unexpected) finding was the prominent downregulation of pathways related to cholesterol metabolism and bile acid formation. This finding provides a sharp distinction between the likely mechanistic events driven by Aquamin^®^ and those responsive to OCA. This finding helps validate the utility of the proteomic screening approach employed here.

## DISCUSSION

A wide range of studies (9–11) has demonstrated the susceptibility of rodents to diet-induced NAFLD. Steatosis develops rapidly (within 4-8 weeks) in rats and mice placed on a high-fat or high-fat and -sugar diet. In contrast, end-stage liver disease – i.e., fibrosis/cirrhosis and tumor formation – does not manifest until much later. Tumor formation has been reported to occur as early as after 36 weeks of feeding, but typically 12 or more months is required for widespread neoplastic disease to be seen (9–11). In our own previous studies (19,20), tumors were widespread in male C57BL/6 mice fed a high-fat diet after 12-18 months but almost non-existent in a cohort of mice euthanized at the 5-month timepoint. Extensive inflammation, parenchymal necrosis and fibrotic nodules were also seen at the later timepoints. Most importantly, our studies demonstrated that providing an adequate supply of dietary minerals along with high-fat feeding substantially reduced the incidence and severity of liver damage. A reduction in tumor formation was especially prominent.

Even though end-stage liver injury in rodents is fully manifested only after an extended feeding period, the cellular and molecular events that bring about damage undoubtedly begin to occur much earlier. It is also likely that interventions that mitigate long-term injury have effects that can be detected much earlier. With this in mind, the present study was carried out to identify hepatic alterations detectable in mice after 16 weeks on a high-fat and -sugar diet compared to control mice on a low-fat diet. As a major part of this study, mineral supplementation along with a different intervention (OCA) with a, presumably, different mechanisms of action were also separately evaluated.

In comparison with low-fat-fed control mice, livers were substantially enlarged in high-fat mice. However, there was no gross evidence of more serious liver damage (i.e., fibrotic nodules or tumors) at the 16-week time-point. At the histological level, steatosis was widespread in virtually every mouse on the high-fat diet (not observed in low-fat controls). Ballooning hepatocyte degeneration was also prominent, but inflammation and collagen deposition were minimal. These effects were accompanied by an increase in serum triglycerides and an increase in serum ALT and AST levels. Consistent with previous findings in both rat (50) and mouse (51) models, proteomic assessment demonstrated substantial differences in the protein expression profile between control (low-fat) animals and those on the high-fat diet. The most dramatic differences were in proteins that regulate fat and carbohydrate metabolism. Increases in both lipid storage and lipid oxidation [breakdown] were seen with high-fat feeding, but moieties that regulate protein and nucleic acid turnover were also diet-sensitive. Proteins involved in cholesterol metabolism and bile acid formation were especially sensitive to the high-fat diet. Together, these findings demonstrate that extensive metabolic dysregulation occurs early in response to high-fat-feeding – in parallel with widespread steatosis but while other structural changes in the liver are minimal. The metabolic changes described here should be seen as a “normal” liver’s attempt to cope with excessive fat and sugar consumption rather than as evidence of impending liver failure in MS-NASH mouse model. Even the rise in ALT and AST levels noted here in MS-NASH mice (reaching 300-500 U/mL), while substantial, are not necessarily evidence of serious liver damage. Acute chemical-induced liver toxicity in mice is associated with ALT / AST levels in the range of 3000-6000 U/mL (12).

In contrast to the substantial differences distinguishing low-fat and high-fat feeding, mineral supplementation – along with the high-fat diet – had no effect on whole body weight, gross liver weight or steatosis. Likewise, there was no improvement in serum triglycerides and no decrease in AST and ALT values as compared to what was seen in high-fat control mice. This is consistent with findings reported earlier that mineral supplementation does not prevent steatosis from developing (19,20) and points to downstream events as targets of Aquamin^®^.

The protein profile observed in response to intervention with Aquamin^®^ supports this view. While a select few proteins involved directly in fat metabolism / lipid transport were modulated with Aquamin^®^, the most striking Aquamin^®^-induced protein changes relate to decreased amino acid metabolism as well as a reduction in purine and pyrimidine catabolism. Also prominent was the

increased expression of moieties that are part of the epithelial cell differentiation process. Reactome pathway analysis identified keratinization, cornified envelope formation, hemidesmosome formation, junction formation and gap junction assembly as significantly altered pathways. Proteins involved in microtubule assembly were also upregulated in response to Aquamin^®^ and several pathways related to protein trafficking within the cell or between interior and cell surface were identified.

How the protein changes observed in response to Aquamin^®^ in the current study and the reduction in liver tumor formation seen previously in our earlier long-term studies (19,20) are related is not fully understood. In the long-term studies with C57BL/6 mice on a HFWD, formation of multiple neoplastic lesions including aberrant foci, both non-regenerative and regenerative hyperplastic nodules, hepatic adenomas and hepatocellular carcinomas were seen. The majority of the tumors occurred in male mice though females were not completely protected. Mice fed a healthy rodent chow diet had many fewer tumors, but even these were not completely protected after 18 months. Hepatic tumor formation in the context of a high-fat diet and steatosis has been extensively studied and multiple contributing mechanisms are thought to be involved (3,11,16,17). Perhaps among them, the most important is lipotoxicity-driven oxidative stress. Oxidant burden, leading to a highly “mutagenic environment” concurrent with high cell turn over resulting from repeated cycles of injury and repair, is a fertile ground for tumor initiation and progression. The formation of toxic and carcinogenic bile acids (a manifestation of lipid over-feeding) may also contribute to tumor formation. The present study identified Aquamin®-sensitive protein changes that suggest reduced catabolism of fats (as well as reduced amino acid, purine and pyrimidine metabolism); all of which could be expected to lower oxidant burden. The reduction in multiple cytochrome P450 enzymes is supportive. The cytochrome P450 enzymes are strongly expressed in liver and rise as part of the organ’s response to a toxic environment. Of interest, previous studies have demonstrated that strontium (52), selenium (53) and members of the lanthanide family of rare earth elements (54) – all components of Aquamin^®^ – protect the liver against oxidative stress. While liver tumor formation, *per se,* was not addressed in these earlier studies, reduced oxidative stress and reduced tumor formation are, certainly, consistent.

Aquamin^®^-induced epithelial cell differentiation provides another potential mechanism to counteract cancer-promotion. A wide range of empirical information supports this view. Epidemiological studies have clearly demonstrated the inverse relationship between differentiation in epithelial cells and tumor incidence in many tissues including those of the gastrointestinal tract (55–59). Animal studies have confirmed this relationship (60,29,30) and cell culture studies have provided mechanistic insight into the inverse relationship between differentiation and reduced tumor formation (61,62). In addition to a reduction in tumor incidence, the expression of differentiation features (assessed histologically) and improved prognosis is correlated with both pre-malignant and malignant epithelial cell tumors (63–67). Calcium, which is the quintessential “driver” of differentiation in epithelial cells (68), is the most abundant mineral in Aquamin^®^. Thus, epithelial cell differentiation induced by calcium in the multi-mineral product may underlie liver tumor suppression by Aquamin®. It is well-documented, furthermore, that several of the trace elements in the multi-mineral product can act like calcimimetic agonists, promoting a “left-shift” in the calcium dose-response curve. A higher affinity for the extracellular calcium-sensing receptor than calcium itself has been documented with several of the cationic trace elements in Aquamin^®^ (29, 69–71). Similarly, extracellular calcium-sensing receptor has also been expressed in rat hepatocytes (72). Also consistent with this idea is the fact that keratin upregulation was also seen in low-fat mice, though the degree of induction was lower than that observed with Aquamin^®^ itself.

In both Aquamin®-treated animals and low-fat fed mice, the strong keratin upregulation reflected a response in a subset of animals. Why only some animals demonstrated this protein signature is not known, but the finding raises an interesting question: Does the variability in response from animal-to-animal help explain why virtually every mouse fed a high-fat diet rapidly develops steatosis but only a subset progresses to more serious disease? This question cannot be addressed with the presently available data but we have recently initiated a large linear study in which liver biopsies are obtained from mice on a high-fat diet after 16-weeks on diet (as in the present study) following which the animals are maintained for a total of 18-months in order to determine if there is an inverse correlation between keratin expression at the early time-point and tumor formation later.

Although the focus of this study was on mineral supplementation, we also included OCA as a separate intervention. OCA has been shown to inhibit features of NAFLD / NASH in human patients (21–24) and in mice (25). In the current study, OCA produced a modest reduction in steatosis (histology) and a corresponding reduction in serum triglyceride, AST and ALT levels. In the proteomic screen, there was some overlap in upregulated proteins between Aquamin^®^ and OCA, but OCA did not have the same effect on epithelial differentiation proteins as was seen with Aquamin®. In regard to downregulated proteins, the changes seen with OCA were related to cholesterol metabolism and bile acid synthesis, consistent with OCA’s function as a Farnesoid X Receptor agonist (21–24). Given these observations, it is unlikely that the two agents – Aquamin^®^ and OCA – would overlap mechanistically in regard to tumor suppression in a meaningful way. Whether they might produce synergistic anti-tumor activity is not known but is a question worth addressing.

NAFLD is a rising public health challenge. Currently, public health measures focus on “lifestyle” changes, especially diet, to prevent the development of steatosis. Such an approach will not work with everyone. Perhaps efforts directed at reducing the downstream consequences of fatty liver infiltration would be of value. It should be noted in this regard that a majority of individuals in Western society do not meet USDA daily intake guidelines for calcium (73). Many individuals are also deficient in magnesium (74) and, presumably, in other minerals that are nutritionally associated with calcium and magnesium. Inadequate mineral intake is not limited to individuals consuming a western style diet. A recent study demonstrated that a majority of individuals in many developing regions of the world also lacked an adequate amount of calcium in their diet (75,76). Whether the mineral supplement used here or some other formulation might provide a way to insure adequate mineral intake remains to be seen.

We have recently completed a 90-day pilot phase trial in which 30 healthy human subjects were randomized to receive Aquamin^®^ formulated to provide 800 mg of calcium per day, calcium carbonate at the same level or placebo (37,77). To summarize, no safety or tolerability issues were seen with Aquamin^®^. At the same time, colon biopsies obtained before and after treatment demonstrated upregulation of several differentiation-related proteins in the colonic mucosa. In the calcium-alone group, differentiation proteins were also induced, but the levels of increase were lower than seen with Aquamin®. Finally, a decrease in the levels of certain primary and secondary bile acids was also observed in subjects receiving Aquamin^®^ in conjunction with altered gut microbial profile. These metabolomic and microbial changes were not observed with calcium alone. While the focus of these clinical studies has been colonic health, the same approach may provide benefit in the liver.

In summary, our past studies have clearly demonstrated the importance of adequate mineral intake for preventing the downstream consequences of fatty liver accumulation in a murine model. The present studies provide mechanistic insight into how mineral supplementation may contribute to reduction in liver tumor formation, one of the most devastating consequences of fatty liver disease in the face of steatosis.

## DATA AVAILABILITY STATEMENT

All relevant data are within the manuscript and its Supporting Information files. The mass spectrometry proteomics data are available on ProteomeXchange Consortium (PRIDE partner repository) – dataset identifier PXD030954.

## ETHICS STATEMENT

For this murine study we utilized NASH mouse model known as MS-NASH. The entire in-life portion of the study and euthanasia was carried out by Crown Bio, Inc., at their Lafayette, LA (USA) facility under the approved Standard Operating Procedures at the testing site. All procedures involving live animals were reviewed and approved by the Institutional Animal Care and Use Committee (IACUC). Crown Bio, Inc., is an American Association for Accreditation of Laboratory Animal Care (AAALAC)-accredited institution.

## AUTHOR CONTRIBUTIONS

All the listed authors have contributed toward this project to attain authorship status according to ICMJE criteria. Their specific contributions are as follows; Conceptualization: MNA, JV; Methodology, MNA, JV; Software, MNA; Validation, MNA, SDM, RNK, JV; Formal Analysis, MNA, SDM, RNK, IH, DZ, MAHJ-M; Investigation, MNA, SDM, RNK, IH, DZ, MAHJ-M, JV; Resources, MNA, JV; Data Curation, MNA, SDM, RNK, IH, DZ, MAHJ-M; Writing-Original Draft Preparation, MNA, JV; Writing-Review & Editing, MNA, SDM, RNK, IH, DZ, MAHJ-M, JV; Visualization, MNA; Supervision, MNA, JV; Project Administration, MNA, JV; Funding Acquisition, JV. All authors have read and agreed to this version of the manuscript.

## ACKNOWLEDGMENTS

We thank Marigot LTD (Cork, Ireland) for providing Aquamin^®^ Soluble as a gift. Marigot LTD had no role or influence in study design, data collection and analysis, decision to publish, or preparation of the manuscript. We also thank the Proteomics Resource Facility (Pathology Department) for assistance with proteomic data acquisition.

## SUPPLEMENTARY MATERIAL

### The following files are available as part of supplemental information

S Table 1: Mineral Composition of Aquamin^®^ Soluble; S Table 2: Common and unique upregulated proteins (at 2-fold change threshold); S Table 3: Common and unique downregulated proteins (at 2fold change threshold); S Table 4: Comprehensive list of upregulated (and significantly altered) proteins with Aquamin in high-fat mice; S Table 5: Top pathways associated with significantly upregulated proteins (shown in S Table 4) altered with Aquamin; S Table 6: Comprehensive list of downregulated (and significantly altered) proteins with Aquamin in high-fat mice; S Table 7: Top pathways associated with significantly downregulated proteins (shown in S Table 5) altered with Aquamin; S Table 8: String enrichment analysis of upregulated proteins (presented in S Table 4); S Table 9: String enrichment analysis of downregulated proteins (presented in S Table 6); S Table 10: Upregulated Proteins by an unbiased proteomic screening of C57BL6 mice on low-fat diet; S Table 11: Downregulated Proteins by an unbiased proteomic screening of C57BL6 mice on low-fat diet; S Table 12: Top pathways associated with upregulated proteins altered with low-fat diet in C57BL6 mice; S Table 13: Top pathways associated with downregulated proteins altered with low-fat diet in C57BL6 mice; S Table 14: Upregulated Proteins by an unbiased proteomic screening with Obeticholic acid (OCA) in high-fat mice; S Table 15: Downregulated Proteins by an unbiased proteomic screening with Obeticholic acid (OCA) in high-fat mice; S Table 16: Top pathways associated with upregulated proteins altered with Obeticholic acid (OCA); S Table 17: Top pathways associated with downregulated proteins altered with Obeticholic acid (OCA)

