## Supplemental Table 13 for "Liver Protein Expression in Nash Mice on a High-Fat Diet"

**Supplement Table 13. Top pathways associated with downregulated proteins altered with low-fat diet in C57BL6 mice**

| Pathway name | Entities pValue | Mapped entities |
|---|---|---|
| NoRC negatively regulates rRNA expression | $1.27\times10^{-9}$ | H2ax;H3c1;Hist2h2aa1;H4c1;H3-3;H2az2;H2bc3;H3c2;Hist1h2af |
| SIRT1 negatively regulates rRNA expression | $2.11\times10^{-9}$ | H2ax;H3c1;Hist2h2aa1;H4c1;H3-3;H2az2;H2bc3;H3c2;Hist1h2af |
| RNA Polymerase I Promoter Opening | $2.11\times10^{-9}$ | H2ax;H3c1;Hist2h2aa1;H4c1;H3-3;H2az2;H2bc3;H3c2;Hist1h2af |
| Negative epigenetic regulation of rRNA expression | $2.69\times10^{-9}$ | H2ax;H3c1;Hist2h2aa1;H4c1;H3-3;H2az2;H2bc3;H3c2;Hist1h2af |
| Activated PKN1 stimulates transcription of AR (androgen receptor) regulated genes KLK2 and KLK3 | $3.40\times10^{-9}$ | H2ax;H3c1;Hist2h2aa1;H4c1;H3-3;H2az2;H2bc3;H3c2;Hist1h2af |
| Assembly of the ORC complex at the origin of replication | $8.18\times10^{-9}$ | H2ax;H3c1;Hist2h2aa1;H4c1;H3-3;H2az2;H2bc3;H3c2;Hist1h2af |
| Transcriptional regulation by small RNAs | $1.59\times10^{-8}$ | H2ax;H3c1;Hist2h2aa1;H4c1;H3-3;H2az2;H2bc3;Q3UHK8;H3c2;Hist1h2af;Nup35 |
| PRC2 methylates histones and DNA | $1.80\times10^{-8}$ | H2ax;H3c1;Hist2h2aa1;H4c1;H3-3;H2az2;H2bc3;H3c2;Hist1h2af |
| Gene Silencing by RNA | $4.75\times10^{-8}$ | H2ax;H3c1;Hist2h2aa1;H4c1;H3-3;H2az2;H2bc3;Q3UHK8;H3c2;Hist1h2af;Nup35 |
| RUNX1 regulates genes involved in megakaryocyte differentiation and platelet function | $8.39\times10^{-8}$ | H2ax;H3c1;Hist2h2aa1;H4c1;H3-3;H2az2;H2bc3;H3c2;Hist1h2af |
| RHO GTPases activate PKNs | $1.14\times10^{-7}$ | H2ax;H3c1;Hist2h2aa1;H4c1;H3-3;H2az2;H2bc3;H3c2;Hist1h2af |
| RNA Polymerase I Promoter Escape | $2.65\times10^{-7}$ | H2ax;H3c1;Hist2h2aa1;H4c1;H3-3;H2az2;H2bc3;H3c2;Hist1h2af |
| Positive epigenetic regulation of rRNA expression | $2.65\times10^{-7}$ | H2ax;H3c1;Hist2h2aa1;H4c1;H3-3;H2az2;H2bc3;H3c2;Hist1h2af |
| B-WICH complex positively regulates rRNA expression | $2.65\times10^{-7}$ | H2ax;H3c1;Hist2h2aa1;H4c1;H3-3;H2az2;H2bc3;H3c2;Hist1h2af |
| Senescence-Associated Secretory Phenotype (SASP) | $4.44\times10^{-7}$ | H2ax;H3c1;Hist2h2aa1;H4c1;H3-3;H2az2;H2bc3;H3c2;Hist1h2af |
| Oxidative Stress Induced Senescence | $1.74\times10^{-7}$ | H2ax;H3c1;Hist2h2aa1;H4c1;H3-3;H2az2;H2bc3;H3c2;Hist1h2af |
| RNA Polymerase I Promoter Clearance | $1.93\times10^{-6}$ | H2ax;H3c1;Hist2h2aa1;H4c1;H3-3;H2az2;H2bc3;H3c2;Hist1h2af |
| Epigenetic regulation of gene expression | $2.13\times10^{-6}$ | H2ax;H3c1;Hist2h2aa1;H4c1;H3-3;H2az2;H2bc3;H3c2;Hist1h2af |
| RNA Polymerase I Transcription | $2.13\times10^{-6}$ | H2ax;H3c1;Hist2h2aa1;H4c1;H3-3;H2az2;H2bc3;H3c2;Hist1h2af |
| DNA Damage/Telomere Stress Induced Senescence | $2.96\times10^{-6}$ | H2ax;Hist2h2aa1;H4c1;Hmga1;H2az2;H1-5;H2bc3;Hist1h2af |

| | | |
|---|---|---|
| Cellular Senescence | $3.41\times10^{-6}$ | H2ax;H3c1;Hist2h2aa1;H4c1;H3-3;Hmga1;H2az2;H1-5;H2bc3;H3c2;Hist1h2af |
| Base-Excision Repair, AP Site Formation | $4.35\times10^{-6}$ | H2ax;Hist2h2aa1;H4c1;H2az2;Q6R2P8;H2bc3;Hist1h2af |
| Estrogen-dependent gene expression | $5.98\times10^{-6}$ | H2ax;H3c1;Hist2h2aa1;H4c1;H3-3;H2az2;H2bc3;H3c2;Hist1h2af |
| Assembly of the pre-replicative complex | $1.36\times10^{-5}$ | H2ax;H3c1;Hist2h2aa1;H4c1;H3-3;H2az2;H2bc3;H3c2;Hist1h2af |
| Inhibition of DNA recombination at telomere | $1.51\times10^{-5}$ | H2ax;Hist2h2aa1;H4c1;H2az2;H2bc3;Hist1h2af |
| Recognition and association of DNA glycosylase with site containing an affected purine | $1.51\times10^{-5}$ | H2ax;Hist2h2aa1;H4c1;H2az2;H2bc3;Hist1h2af |
| Depurination | $2.02\times10^{-5}$ | H2ax;Hist2h2aa1;H4c1;H2az2;H2bc3;Hist1h2af |
| Cleavage of the damaged purine | $2.02\times10^{-5}$ | H2ax;Hist2h2aa1;H4c1;H2az2;H2bc3;Hist1h2af |
| Condensation of Prophase Chromosomes | $3.05\times10^{-5}$ | H2ax;Hist2h2aa1;H4c1;H2az2;H2bc3;Hist1h2af |
| RHO GTPase Effectors | $4.21\times10^{-5}$ | H2ax;Myl9;Hist2h2aa1;H4c1;H2bc3;Myl12b;Evl;H3c1;Src;H3-3;H2az2;H3c2;Hist1h2af;Wipf3 |
| DNA Replication Pre-Initiation | $4.25\times10^{-5}$ | H2ax;H3c1;Hist2h2aa1;H4c1;H3-3;H2az2;H2bc3;H3c2;Hist1h2af |
| Base Excision Repair | $8.80\times10^{-5}$ | H2ax;Hist2h2aa1;H4c1;H2az2;Q6R2P8;H2bc3;Hist1h2af |
| ESR-mediated signaling | $8.88\times10^{-5}$ | H2ax;H3c1;Hist2h2aa1;H4c1;H3-3;Src;H2az2;H2bc3;H3c2;Hist1h2af |
| Nucleosome assembly | $1.61\times10^{-4}$ | H2ax;Hist2h2aa1;H4c1;H2az2;H2bc3;Hist1h2af |
| Deposition of new CENPA-containing nucleosomes at the centromere | $1.61\times10^{-4}$ | H2ax;Hist2h2aa1;H4c1;H2az2;H2bc3;Hist1h2af |
| HDACs deacetylate histones | $2.12\times10^{-4}$ | H3c1;Hist2h2aa1;H4c1;H2bc3;H3c2;Hist1h2af |
| Chylomicron remodeling | $3.96\times10^{-4}$ | Apoa2;Apoa4;Apoc2 |
| Chylomicron assembly | $3.96\times10^{-4}$ | Apoa2;Apoa4;Apoc2 |
| DNA Replication | $4.32\times10^{-4}$ | H2ax;H3c1;Hist2h2aa1;H4c1;H3-3;H2az2;H2bc3;H3c2;Hist1h2af |
| Mitotic Prophase | $4.78\times10^{-4}$ | H2ax;Hist2h2aa1;H4c1;H2az2;H2bc3;Hist1h2af;Nup35 |
| Transcriptional regulation by RUNX1 | $5.38\times10^{-4}$ | H2ax;H3c1;Hist2h2aa1;H4c1;H3-3;H2az2;H2bc3;H3c2;Hist1h2af |
| RMTs methylate histone arginines | $6.01\times10^{-4}$ | H3c1;Hist2h2aa1;H4c1;H3c2;Hist1h2af |
| Signaling by Nuclear Receptors | $7.81\times10^{-4}$ | H2ax;H3c1;Hist2h2aa1;H4c1;H3-3;Src;H2az2;H2bc3;H3c2;Hist1h2af |
| Telomere Maintenance | $9.49\times10^{-4}$ | H2ax;Hist2h2aa1;H4c1;H2az2;H2bc3;Hist1h2af |
| Fatty acyl-CoA biosynthesis | 0.001 | Acly;Fasn;Acaca;Elovl5 |
| Plasma lipoprotein assembly | 0.002 | Apoa2;Apoa4;Apoc2 |
| Signaling by Rho GTPases | 0.002 | H2ax;Arhgef2;Myl9;Hist2h2aa1;H4c1;H2bc3;Myl12b;Sos1;Evl;H3c1;Src;H3-3;H2az2;Tex2;Arhgap23;H3c2;Hist1h2af;Basp1;Wipf3 |
| Fatty acid metabolism | 0.003 | Ehhadh;Acot11;Acly;Fasn;Abcd1;Acaca;Acly;Elovl5 |

| | | |
|---|---|---|
| Signaling by Rho GTPases, Miro GTPases and RHOBTB3 | 0.003 | H2ax;Arhgef2;Myl9;Hist2h2aa1;H4c1;H2bc3;Myl12b;Sos1;Evl;H3c1;Src;H3-3;H2az2;Tex2;Arhgap23;H3c2;Hist1h2af;Basp1;Wipf3 |
| HATs acetylate histones | 0.003 | H3c1;H4c1;H2bc3;H3c2 |
| ChREBP activates metabolic gene expression | 0.003 | Fasn;Acaca |
| Chromosome Maintenance | 0.004 | H2ax;Hist2h2aa1;H4c1;H2az2;H2bc3;Hist1h2af |
| Metabolism of lipids | 0.004 | Hsd17b13;Fasn;Abcd1;Acaca;Lpcat1;Mgll;Ehhadh;Pla2g6;Dgat1;Acot11;Acly;Acly;Osbpl3;Elovl5;Smpd3;Acss3;Fabp2 |
| Acyl chain remodeling of DAG and TAG | 0.004 | Mgll;Dgat1 |
| Linoleic acid (LA) metabolism | 0.006 | Abcd1;Elovl5 |
| HDMs demethylate histones | 0.006 | H3c1;H4c1;H3c2 |
| Plasma lipoprotein remodeling | 0.007 | Apoa2;Apoa4;Apoc2 |
| Smooth Muscle Contraction | 0.009 | Sorbs1;Myl9;Myl12b |
| Beta-oxidation of very long chain fatty acids | 0.011 | Ehhadh;Abcd1 |
| Pyrimidine salvage | 0.011 | Uck1;Tymp |
| Formyl peptide receptors bind formyl peptides and many other ligands | 0.012 | Anxa1;Hebp1 |
| alpha-linolenic (omega3) and linoleic (omega6) acid metabolism | 0.014 | Abcd1;Elovl5 |
| alpha-linolenic acid (ALA) metabolism | 0.014 | Abcd1;Elovl5 |
| Dissolution of Fibrin Clot | 0.014 | S100a10;Anxa2 |
| Nonhomologous End-Joining (NHEJ) | 0.016 | H2ax;H4c1;H2bc3 |
| Regulation of KIT signaling | 0.017 | Src;Sos1 |
| RET signaling | 0.019 | Pdlim7;Src;Sos1 |
| GRB2:SOS provides linkage to MAPK signaling for Integrins | 0.019 | Src;Sos1 |
| Formation of Senescence-Associated Heterochromatin Foci (SAHF) | 0.019 | Hmga1;H1-5 |
| Retinoid metabolism and transport | 0.020 | Apoa2;Apoa4;Apoc2 |
| ABC transporters in lipid homeostasis | 0.024 | Abcd1;Abcd2 |
| Interleukin-7 signaling | 0.024 | H3c1;H3c2 |
| Metabolism of fat-soluble vitamins | 0.025 | Apoa2;Apoa4;Apoc2 |
| PKMTs methylate histone lysines | 0.028 | H3c1;H4c1;H3c2 |
| Class I peroxisomal membrane protein import | 0.029 | Abcd1;Abcd2 |
| Recruitment and ATM-mediated phosphorylation of repair and signaling proteins at DNA double strand breaks | 0.030 | H2ax;H4c1;H2bc3 |

| | | |
|---|---|---|
| DNA Double Strand Break Response | 0.032 | H2ax;H4c1;H2bc3 |
| RHO GTPases activate PAKs | 0.032 | Myl9;Myl12b |
| Visual phototransduction | 0.035 | Apoa2;Fntb;Apoa4;Apoc2 |
| Cellular responses to stress | 0.038 | H2ax;H3c1;Hist2h2aa1;H4c1;H3-3;Hmga1;H2az2;H1-5;H2bc3;H3c2;Hist1h2af;Nup35 |
| Cellular responses to stimuli | 0.040 | H2ax;H3c1;Hist2h2aa1;H4c1;H3-3;Hmga1;H2az2;H1-5;H2bc3;H3c2;Hist1h2af;Nup35 |
| Regulation of gap junction activity | 0.041 | Src |
| Activated NTRK3 signals through PI3K | 0.041 | Src |
| Triglyceride metabolism | 0.045 | Dgat1;Fabp2 |
| Nucleotide salvage | 0.045 | Uck1;Tymp |
| Chromatin organization | 0.045 | H3c1;Hist2h2aa1;H4c1;H2bc3;H3c2;Hist1h2af |
| Chromatin modifying enzymes | 0.045 | H3c1;Hist2h2aa1;H4c1;H2bc3;H3c2;Hist1h2af |
| Metabolism of nucleotides | 0.046 | Dnph1;Ak1;Uck1;Tymp |

The pathways listed here are altered by the significantly downregulated proteins with the control low-fat diet in C57BL6 mice presented in Supplement Table 11 using an unbiased approach. Reactome (v78) was used to generate the pathway analysis report for species *Mus musculus*. The significance (*p*-value) is calculated by the overrepresentation analysis (hypergeometric distribution).
