## Supplemental Table 14 for "Liver Protein Expression in Nash Mice on a High-Fat Diet"

**Supplement Table 14. Upregulated Proteins by an unbiased proteomic screening with Obeticholic acid (OCA) in high-fat mice**

| | | MS-NASH | | C57BL6 |
|---|---|---|---|---|
| **Proteins** | **Genes** | **OCA** | **Aquamin** | **Control** |
| Nuclear transport factor 2 | Nutf2 | 10.29±20.76 | 12.12±24.59 | 8.16±16.12 |
| Fucose mutarotase | Fuom | 10.03±19.94 | 10.28±20.61 | 8.72±17.40 |
| Glycine cleavage system H protein, mitochondrial | Gcsh | 9.05±17.88 | 7.63±14.93 | 11.09±21.41 |
| Ras-related protein Rap-1A | Rap1a | 5.83±10.73 | 5.20±9.18 | 5.66±10.21 |
| IgG receptor FcRn large subunit p51 | Fcgrt | 5.74±10.39 | 4.12±6.83 | 4.17±6.07 |
| Keratin, type II cytoskeletal 79 | Krt79 | 5.73±7.97 | 26.34±43.11 | 5.40±4.07 |
| Pigment epithelium-derived factor | Serpinf1 | 5.58±10.41 | 6.03±10.85 | 3.34±5.88 |
| SH3 domain-containing protein 21[#] | Sh3d21 | 5.43±10.12 | 8.30±16.34 | 6.05±10.62 |
| Tubulin beta-5 chain | Tubb5 | 5.06±9.36 | 5.81±10.51 | 4.90±9.25 |
| 60S ribosomal protein L36 | Rpl36 | 4.95±8.52 | 4.32±7.31 | 4.35±8.31 |
| Transmembrane protein 14C | Tmem14c | 4.82±8.80 | 5.56±10.29 | 5.97±10.43 |
| Ras-related protein R-Ras | Rras | 4.75±8.44 | 5.46±9.57 | 6.32±11.74 |
| Tubulin beta-4A chain | Tubb4a | 4.73±8.30 | 4.85±8.51 | 3.76±6.90 |
| Aquaporin-4 | Aqp4 | 3.99±3.09 | 0.92±0.56 | 0.29±0.20 |
| 60S acidic ribosomal protein P1 | Rplp1 | 3.61±5.97 | 3.53±5.59 | 4.84±8.27 |
| Splicing factor 3B subunit 4 | Sf3b4 | 3.57±5.19 | 3.03±4.20 | 3.46±4.27 |
| Galectin-related protein | Lgalsl | 3.52±5.77 | 3.26±5.14 | 2.86±4.63 |
| Proteasome subunit beta type-3 | Psmb3 | 3.50±5.41 | 3.56±5.56 | 3.27±4.30 |
| Endophilin-B1 | Sh3glb1 | 3.47±5.80 | 3.93±6.42 | 3.27±5.18 |
| ATP-dependent translocase ABCB1 | Abcb1a | 3.40±2.67 | 1.54±0.91 | 0.91±0.72 |
| Galactose-1-phosphate uridylyltransferase | Galt | 3.02±4.19 | 2.14±2.67 | 2.84±3.21 |
| U8 snoRNA-decapping enzyme | Nudt16 | 3.02±4.24 | 4.03±5.49 | 3.26±4.15 |
| 5'-AMP-activated protein kinase subunit beta-1 | Prkab1 | 3.00±3.74 | 3.05±4.30 | 1.92±2.61 |
| Alpha-1-acid glycoprotein 1 | Orm1 | 2.93±5.05 | 3.98±6.98 | 6.12±9.25 |
| Glutathione S-transferase A4 | Gsta4 | 2.90±2.09 | 1.63±1.12 | 1.95±1.06 |
| Glutaredoxin-1 | Glrx | 2.85±3.94 | 2.58±3.23 | 3.76±5.34 |
| Copper transport protein ATOX1 | Atox1 | 2.84±4.30 | 3.44±5.40 | 1.64±2.28 |
| 2-hydroxyacyl-CoA lyase 2 | Ilvbl | 2.82±4.22 | 2.99±4.38 | 2.91±4.43 |
| Coatomer subunit epsilon | Cope | 2.78±4.10 | 3.50±5.59 | 3.27±4.74 |
| Protein PAT1 homolog 1[#] | Patl1 | 2.74±2.40 | 4.99±4.23 | 3.09±2.72 |
| Glutathione S-transferase A2 | Gsta2 | 2.66±0.92* | 1.04±0.20 | 1.27±0.53 |
| All-trans-retinol dehydrogenase [NAD(+)] ADH4 | Adh4 | 2.66±2.14 | 1.16±0.72 | 1.77±1.64 |
| Putative RNA-binding protein Luc7-like 1 | Luc7l | 2.56±3.40 | 3.20±4.78 | 2.67±3.04 |

| | | | | |
|---|---|---|---|---|
| Histidine ammonia-lyase | Hal | 2.54±0.83* | 0.71±0.27 | 1.88±0.49* |
| S-methylmethionine--homocysteine S-methyltransferase BHMT2 | Bhmt2 | 2.51±3.26 | 2.70±3.60 | 2.01±2.67 |
| Peptidyl-prolyl cis-trans isomerase NIMA-interacting 1 | Pin1 | 2.51±2.90 | 2.52±2.87 | 2.43±2.54 |
| Cytochrome b-c1 complex subunit 10 | Uqcr11 | 2.45±3.24 | 2.47±3.24 | 3.40±4.36 |
| NADH-ubiquinone oxidoreductase chain 5 | Mtnd5 | 2.44±3.18 | 2.42±3.25 | 2.55±3.40 |
| Nuclear cap-binding protein subunit 1 | Ncbp1 | 2.41±3.23 | 2.54±3.24 | 2.29±2.99 |
| TIP41-like protein | Tiprl | 2.36±2.70 | 2.18±2.36 | 1.63±1.88 |
| Exportin-5 | Xpo5 | 2.32±2.68 | 1.77±1.75 | 1.82±2.36 |
| Aldo-keto reductase family 1 member C18 | Akr1c18 | 2.31±2.65 | 2.74±3.86 | 2.52±2.51 |
| NADH dehydrogenase [ubiquinone] 1 alpha subcomplex subunit 11 | Ndufa11 | 2.31±2.93 | 1.71±1.62 | 1.55±1.16 |
| Carboxypeptidase B2 | Cpb2 | 2.29±2.74 | 2.39±2.63 | 3.39±4.66 |
| Signal recognition particle 19 kDa protein | Srp19 | 2.29±3.07 | 2.46±3.39 | 3.08±4.09 |
| Tubulin beta-2A chain | Tubb2a | 2.26±2.77 | 2.30±2.80 | 1.00±1.29 |
| Aldose reductase-related protein 2 | Akr1b8 | 2.25±2.73 | 2.75±3.39 | 2.84±3.75 |
| STIP1 homology and U box-containing protein 1 | Stub1 | 2.22±2.68 | 2.07±2.40 | 2.33±2.80 |
| Small nuclear ribonucleoprotein E | Snrpe | 2.22±2.99 | 2.90±4.12 | 2.86±3.61 |
| Platelet-activating factor acetylhydrolase IB subunit gamma | Pafah1b3 | 2.21±1.34 | 2.45±1.29 | 1.80±0.90 |
| Acyl-CoA-binding protein | Dbi | 2.20±3.02 | 2.05±2.43 | 2.05±2.30 |
| Acetyl-coenzyme A transporter 1 | Slc33a1 | 2.20±2.51 | 2.13±2.62 | 2.04±2.75 |
| WD repeat-containing protein 18 | Wdr18 | 2.18±2.66 | 2.03±2.17 | 2.22±2.64 |
| Thyroid hormone-inducible hepatic protein | Thrsp | 2.18±2.04 | 1.24±0.95 | 0.70±0.46 |
| COMM domain-containing protein 8 | Commd8 | 2.16±1.99 | 2.26±2.73 | 2.63±2.14 |
| Nascent polypeptide-associated complex subunit alpha, muscle-specific form | Naca | 2.15±2.63 | 2.13±2.58 | 2.29±2.60 |
| Sulfotransferase 1A1 | Sult1a1 | 2.14±1.74 | 1.35±0.92 | 1.61±1.90 |
| ATP-binding cassette sub-family D member 4 | Abcd4 | 2.12±1.99 | 1.50±1.19 | 4.49±4.10 |
| Glutathione S-transferase theta-2 | Gstt2 | 2.12±1.22 | 1.31±0.65 | 0.91±0.45 |
| Neuroplastin | Nptn | 2.11±2.67 | 2.44±3.07 | 2.33±2.72 |
| Ubiquinol-cytochrome-c reductase complex assembly factor 3 | Uqcc3 | 2.11±1.93 | 1.82±1.44 | 1.57±1.50 |
| Histone-lysine N-methyltransferase 2A | Kmt2a | 2.08±0.33* | 1.14±0.61 | 1.61±0.32* |
| SH3-containing GRB2-like protein 3-interacting protein 1 | Sgip1 | 2.08±2.33 | 1.41±0.96 | 1.82±2.47 |
| Ig kappa chain V-V region MOPC 149 | n/a | 2.06±2.70 | 4.02±4.71 | 1.52±2.10 |
| Protein quaking | Qki | 2.06±2.46 | 2.49±3.17 | 1.99±2.52 |
| Small nuclear ribonucleoprotein Sm D1 | Snrpd1 | 2.05±2.04 | 2.03±2.00 | 2.13±1.84 |
| AP-2 complex subunit sigma | Ap2s1 | 2.05±2.57 | 2.45±3.06 | 2.61±3.59 |
| [Protein ADP-ribosylarginine] hydrolase | Adprh | 2.05±2.29 | 2.36±2.84 | 1.53±1.69 |
| 4-hydroxy-2-oxoglutarate aldolase, mitochondrial | Hoga1 | 2.05±1.87 | 1.27±0.85 | 3.81±2.81 |
| Ancient ubiquitous protein 1 | Aup1 | 2.04±2.78 | 2.00±2.73 | 2.45±3.38 |

| | | | | |
|---|---|---|---|---|
| Protein dpy-30 homolog | Dpy30 | 2.04±2.36 | 1.74±0.94 | 1.17±0.28 |
| ADP-ribosylation factor 4 | Arf4 | 2.02±2.64 | 2.31±2.87 | 1.84±2.57 |
| Phosphatidylethanolamine N-methyltransferase | Pemt | 2.02±2.17 | 1.81±1.56 | 2.60±3.30 |
| Transthyretin | Ttr | 2.02±2.15 | 1.57±0.94 | 0.98±0.41 |
| Cation channel sperm-associated protein subunit beta[#] | Catsperb | 2.00±0.90 | 1.42±0.54 | 2.55±1.47 |
| Bile salt export pump | Abcb11 | 2.00±0.35* | 1.03±0.07 | 1.96±0.57* |

These values represent average (± standard deviation) fold-change of abundance ratios for each altered protein compared to the high-fat control group (MS-NASH mice on high-fat) with a cutoff of 2-fold-change in response to OCA intervention. For each upregulated protein with OCA, corresponding values from the other two groups are shown for comparison. The liver samples (from 5 mice in each group) were individually assessed by TMT based differential proteomic expression and data were merged to get averages. Protein FDR Confidence for all proteins was ≤1% except 3 proteins ([#]≤2%). These data are also presented in Figure 3A. FDR: False Discovery Rate.
