## Supplemental Table 15 for "Liver Protein Expression in Nash Mice on a High-Fat Diet"

**Supplement Table 15. Downregulated Proteins by an unbiased proteomic screening with Obeticholic acid (OCA) in high-fat mice**

| Proteins | Genes | MS-NASH OCA | MS-NASH Aquamin | C57BL6 Control |
|---|---|---|---|---|
| 7-alpha-hydroxycholest-4-en-3-one 12-alpha-hydroxylase | Cyp8b1 | 0.268±0.134* | 0.809±0.302 | 1.012±0.462 |
| Tonsoku-like protein | Tonsl | 0.306±0.172* | 0.502±0.313 | 0.342±0.232* |
| Major urinary protein 2 | Mup2 | 0.440±0.187* | 0.586±0.229* | 10.396±5.298 |
| Serum amyloid A-1 protein | Saa1 | 0.447±0.190* | 0.728±0.404 | 3.653±4.613 |
| Major intrinsically disordered NOTCH2-binding receptor 1-like homolog# | Minar2 | 0.451±0.175* | 0.491±0.122* | 1.062±0.232 |
| Centrosomal protein of 170 kDa | Cep170 | 0.461±0.341 | 0.672±0.529 | 0.260±0.244* |
| Ornithine aminotransferase, mitochondrial | Oat | 0.485±0.124* | 0.757±0.355 | 1.681±0.551 |
| Galectin-1 | Lgals1 | 0.488±0.266* | 1.038±0.387 | 0.461±0.192* |

These values represent average (± standard deviation) fold-change of abundance ratios for each altered protein compared to the high-fat control group (MS-NASH mice on high-fat) with a cutoff of 2-fold-change in response to OCA intervention. For each downregulated protein with OCA, corresponding values from the other two groups are shown for comparison. The liver samples (from 5 mice in each group) were individually assessed by TMT based differential proteomic expression and data were merged to get averages. Protein FDR Confidence for all proteins was ≤1% except 1 protein (#≤2%). These data are also presented in Figure 3A. FDR: False Discovery Rate.
