## Supplemental Table 16 for "Liver Protein Expression in Nash Mice on a High-Fat Diet"

**Supplement Table 16. Top pathways associated with upregulated proteins altered with Obeticholic acid (OCA)**

| Pathway Name | Entities pValue | Mapped entities |
|---|---|---|
| mRNA Splicing - Minor Pathway | 0.001 | Sf3b4;Ncbp1;Snrpe;Snrpd1 |
| SLBP independent Processing of Histone Pre-mRNAs | 0.003 | Ncbp1;Snrpe |
| Recycling pathway of L1 | 0.004 | Tubb4a;Tubb2a;P62743 |
| SLBP Dependent Processing of Replication-Dependent Histone Pre-mRNAs | 0.004 | Ncbp1;Snrpe |
| Golgi-to-ER retrograde transport | 0.005 | Pafah1b3;Arf4;Tubb4a;Cope;Tubb2a |
| Microtubule-dependent trafficking of connexons from Golgi to the plasma membrane | 0.007 | Tubb4a;Tubb2a |
| COPI-independent Golgi-to-ER retrograde traffic | 0.008 | Pafah1b3;Tubb4a;Tubb2a |
| Transport of connexons to the plasma membrane | 0.008 | Tubb4a;Tubb2a |
| COPI-mediated anterograde transport | 0.008 | Arf4;Tubb4a;Cope;Tubb2a |
| COPI-dependent Golgi-to-ER retrograde traffic | 0.008 | Arf4;Tubb4a;Cope;Tubb2a |
| L1CAM interactions | 0.020 | Tubb4a;Tubb2a;P62743 |
| Intra-Golgi and retrograde Golgi-to-ER traffic | 0.020 | Pafah1b3;Arf4;Tubb4a;Cope;Tubb2a |
| Carboxyterminal post-translational modifications of tubulin | 0.020 | Tubb4a;Tubb2a |
| Sealing of the nuclear envelope (NE) by ESCRT-III | 0.020 | Tubb4a;Tubb2a |
| The role of GTSE1 in G2/M progression after G2 checkpoint | 0.023 | Tubb4a;Psmb3;Tubb2a |
| Processing of Capped Intronless Pre-mRNA | 0.023 | Ncbp1;Snrpe |
| Glyoxylate metabolism and glycine degradation | 0.027 | Hoga1;Gcsh |
| Gap junction assembly | 0.028 | Tubb4a;Tubb2a |
| Glycine degradation | 0.033 | Gcsh |
| Ion influx/efflux at host-pathogen interface | 0.033 | Atox1 |
| Aggrephagy | 0.035 | Tubb4a;Tubb2a |
| ER to Golgi Anterograde Transport | 0.035 | Arf4;Tubb4a;Cope;Tubb2a |
| Recruitment of NuMA to mitotic centrosomes | 0.039 | Tubb5;Tubb4a;Tubb2a |
| SRP-dependent cotranslational protein targeting to membrane | 0.040 | Srp19;Rplp1;Rpl36 |
| ARMS-mediated activation | 0.041 | Rap1a |
| Galactose catabolism | 0.041 | Galt |
| Nonsense Mediated Decay (NMD) independent of the Exon Junction Complex (EJC) | 0.043 | Ncbp1;Rplp1;Rpl36 |
| Metabolism of RNA | 0.046 | Sf3b4;Ncbp1;Snrpe;Psmb3;Rplp1;Patl1;Rpl36;Snrpd1;Wdr18 |
| Gap junction trafficking | 0.047 | Tubb4a;Tubb2a |
| GDP-fucose biosynthesis | 0.049 | Fuom |

The pathways listed here are altered by the significantly upregulated proteins with the intervention “OCA” presented in Supplement Table 14 using an unbiased approach. Reactome (v78) was used to generate the pathway analysis report for species *Mus musculus*. The significance (*p*-value) is calculated by the overrepresentation analysis (hypergeometric distribution).
