## Supplemental Table 17 for "Liver Protein Expression in Nash Mice on a High-Fat Diet"

**Supplement Table 17. Top pathways associated with downregulated proteins altered with Obeticholic acid (OCA)**

| Pathway name | Entities pValue | Mapped entities |
|---|---|---|
| Sterols are 12-hydroxylated by CYP8B1 | 0.002 | Cyp8b1 |
| Synthesis of Prostaglandins (PG) and Thromboxanes (TX) | 0.011 | Cyp8b1 |
| Synthesis of bile acids and bile salts via 24-hydroxycholesterol | 0.011 | Cyp8b1 |
| Glutamate and glutamine metabolism | 0.011 | Oat |
| Synthesis of bile acids and bile salts via 27-hydroxycholesterol | 0.011 | Cyp8b1 |
| Eicosanoids | 0.015 | Cyp8b1 |
| Nicotinamide salvaging | 0.017 | Cyp8b1 |
| Synthesis of bile acids and bile salts via 7alpha-hydroxycholesterol | 0.019 | Cyp8b1 |
| Endogenous sterols | 0.024 | Cyp8b1 |
| Nicotinate metabolism | 0.027 | Cyp8b1 |
| Synthesis of bile acids and bile salts | 0.028 | Cyp8b1 |
| Bile acid and bile salt metabolism | 0.037 | Cyp8b1 |
| Arachidonic acid metabolism | 0.049 | Cyp8b1 |

The pathways listed here are altered by the downregulated proteins with the intervention “Obeticholic acid (OCA)” presented in Supplement Table 15 using an unbiased approach. Reactome (v78) was used to generate the pathway analysis report for species *Mus musculus*. The significance (*p* value) is calculated by the overrepresentation analysis (hypergeometric distribution).
