## Supplemental Table 2 for "Liver Protein Expression in Nash Mice on a High-Fat Diet"

**Supplement Table 2. Common and unique upregulated proteins (at 2-fold change threshold)**

| Accession | Protein names | Gene Name |
|---|---|---|
| | **Common Proteins among Aquamin, OCA and C57BL6 (low-fat): 46** | |
| Q7TSG5 | SH3 domain-containing protein 21 | Sh3d21 |
| P62743 | AP-2 complex subunit sigma | Ap2s1 |
| Q8R2K1 | Fucose mutarotase | Fuom |
| Q60590 | Alpha-1-acid glycoprotein 1 | Agp1 |
| P99024 | Tubulin beta-5 chain | Tubb5 |
| Q9CQN6 | Transmembrane protein 14C | Tmem14c |
| P47955 | 60S acidic ribosomal protein P1 | Rplp1 |
| P03921 | NADH-ubiquinone oxidoreductase chain 5 | Mtnd5 |
| P47964 | 60S ribosomal protein L36 | Rpl36 |
| Q99J27 | Acetyl-coenzyme A transporter 1 | Acatn |
| Q9JHH6 | Carboxypeptidase B2 | Cpb2 |
| P10833 | Ras-related protein R-Ras | Rras |
| Q9R1P1 | Proteasome subunit beta type-3 | Psmb3 |
| Q9CPX8 | Cytochrome b-c1 complex subunit 10 | Uqcr11 |
| Q61559 | IgG receptor FcRn large subunit p51 | Fcgrt |
| P70295 | Lipid droplet-regulating VLDL assembly factor AUP1 | Aup1 |
| Q03249 | Galactose-1-phosphate uridylyltransferase | Galt |
| P97300 | Neuroplastin | Nptn |
| Q8K023 | Aldo-keto reductase family 1 member C18 | Akr1c18 |
| P31786 | Acyl-CoA-binding protein | Dbi |
| Q91WS4 | S-methylmethionine--homocysteine S-methyltransferase BHMT2 | Bhmt2 |
| P97298 | Pigment epithelium-derived factor | Serpinf1 |
| Q9JK48 | Endophilin-B1 | Sh3glb1 |
| Q9CYI4 | Putative RNA-binding protein Luc7-like 1 | Luc7l |
| Q9CZG3 | COMM domain-containing protein 8 | Commd8 |
| Q8VED9 | Galectin-related protein | Lgalsl |
| Q6P3D0 | U8 snoRNA-decapping enzyme (m7GpppN-mRNA hydrolase) | Nudt16 |
| Q9D7A6 | Signal recognition particle 19 kDa protein | Srp19 |
| O89079 | Coatomer subunit epsilon | Cope |
| P62835 | Ras-related protein Rap-1A | Rap1a |
| Q3TC46 | Protein PAT1 homolog 1 | Patl1 |
| Q9D6F9 | Tubulin beta-4A chain | Tubb4a |
| Q9QUH0 | Glutaredoxin-1 | Glrx |
| P45377 | Aldose reductase-related protein 2 | Akr1b8 |
| Q3UYV9 | Nuclear cap-binding protein subunit 1 | Ncbp1 |
| P61971 | Nuclear transport factor 2 | Nutf2 |
| P62305 | Small nuclear ribonucleoprotein E | Snrpe |
| Q4VBE8 | WD repeat-containing protein 18 | Wdr18 |
| Q8BU33 | 2-hydroxyacyl-CoA lyase 2 | Ilvbl |
| P70670 | Alpha-NAC, muscle-specific form | Naca |
| Q9QUR7 | Peptidyl-prolyl cis-trans isomerase NIMA-interacting 1 | Pin1 |

| | | |
|---|---|---|
| Q91WK5 | Glycine cleavage system H protein, mitochondrial | Gcsh |
| Q9WUD1 | E3 ubiquitin-protein ligase CHIP | Stub1 |
| Q8QZY9 | Splicing factor 3B subunit 4 | Sf3b4 |
| Q8VED5 | Keratin, type II cytoskeletal 79 | Krt79 |
| P62315 | Small nuclear ribonucleoprotein Sm D1 | Snrpd1 |
| | **Common Proteins between OCA and C57BL6 (low-fat): 4** | |
| Q61907 | Phosphatidylethanolamine N-methyltransferase | Pemt |
| A2RTF1 | Cation channel sperm-associated protein subunit beta | Catsperb |
| O89016 | Lysosomal cobalamin transporter ABCD4 | Abcd4 |
| Q9DCU9 | 4-hydroxy-2-oxoglutarate aldolase, mitochondrial | Hoga1 |
| | **Common Proteins between Aquamin and C57BL6 (low-fat): 12** | |
| Q6IFX2 | Keratin, type I cytoskeletal 42 | Krt42 |
| Q3TTY5 | Keratin, type II cytoskeletal 2 epidermal | Krt2 |
| Q9EPB5 | Serine hydrolase-like protein | Serhl |
| Q922U2 | Keratin, type II cytoskeletal 5 | Krt5 |
| Q9QWL7 | Keratin, type I cytoskeletal 17 | Krt17 |
| P35762 | CD81 antigen | Cd81 |
| Q99KE1 | NAD-dependent malic enzyme, mitochondrial | Me2 |
| Q61781 | Keratin, type I cytoskeletal 14 | Krt14 |
| Q3UV17 | Keratin, type II cytoskeletal 2 oral | Krt76 |
| P04104 | Keratin, type II cytoskeletal 1 | Krt1 |
| P02535 | Keratin, type I cytoskeletal 10 | Krt10 |
| Q9Z2K1 | Keratin, type I cytoskeletal 16 | Krt16 |
| | **Common Proteins between Aquamin and OCA: 9** | |
| Q61205 | Platelet-activating factor acetylhydrolase IB subunit alpha1 | Pafah1b3 |
| O08997 | Copper transport protein ATOX1 | Atox1 |
| Q7TMM9 | Tubulin beta-2A chain | Tubb2a |
| Q9R078 | 5'-AMP-activated protein kinase subunit beta-1 | Prkab1 |
| P54923 | [Protein ADP-ribosylarginine] hydrolase | Adprh |
| P61750 | ADP-ribosylation factor 4 | Arf4 |
| P01636 | Ig kappa chain V-V region MOPC 149 | n/a |
| Q8BH58 | TIP41-like protein | Tiprl |
| Q9QYS9 | Protein quaking | Qki |
| | **Unique Proteins to Aquamin: 19** | |
| P14434 | H-2 class II histocompatibility antigen, A-B alpha chain | H2-Aa |
| Q8CHW4 | Translation initiation factor eIF-2B subunit epsilon | Eif2b5 |
| Q99JW4 | LIM and senescent cell antigen-like-containing domain protein 1 | Lims1 |
| P01864 | Ig gamma-2A chain C region secreted form | n/a |
| P04441 | H-2 class II histocompatibility antigen gamma chain | Cd74 |
| P50428 | Arylsulfatase A | Arsa |
| Q61093 | Cytochrome b-245 heavy chain | Cybb |
| Q9DBC7 | cAMP-dependent protein kinase type I-alpha regulatory subunit | Prkar1a |
| Q8K1B8 | Fermitin family homolog 3 | Fermt3 |
| P42225 | Signal transducer and activator of transcription 1 | Stat1 |
| P18468 | H-2 class II histocompatibility antigen, I-A beta chain | H2-Eb1 |

| | | |
|---|---|---|
| A2AWP8 | Rho guanine nucleotide exchange factor 10-like protein | Arhgef10l |
| Q01514 | Guanylate-binding protein 1 | Gbp1 |
| P50543 | Protein S100-A11 | S100a11 |
| P28063 | Proteasome subunit beta type-8 | Psmb8 |
| P24638 | Lysosomal acid phosphatase | Acp2 |
| Q99LU0 | Charged multivesicular body protein 1b-1 | Chmp1b1 |
| Q9Z0M5 | Lysosomal acid lipase/cholesteryl ester hydrolase | Lipa |
| Q9QZ25 | Vascular non-inflammatory molecule 3 | Vnn3 |
| | **Unique Proteins to OCA: 17** | |
| P55200 | Histone-lysine N-methyltransferase 2A | Kmt2a |
| P10648 | Glutathione S-transferase A2 | Gsta2 |
| Q8VD37 | SH3-containing GRB2-like protein 3-interacting protein 1 | Sgip1 |
| P35492 | Histidine ammonia-lyase | Hal |
| P52840 | Sulfotransferase 1A1 | Sult1a1 |
| Q9QYY9 | All-trans-retinol dehydrogenase [NAD(+)] ADH4 | Adh4 |
| Q8K2T4 | Ubiquinol-cytochrome-c reductase complex assembly factor 3 | Uqcc3 |
| P21447 | ATP-dependent translocase ABCB1 | Abcb1a |
| Q61133 | Glutathione S-transferase theta-2 | Gstt2 |
| P55088 | Aquaporin-4 | Aqp4 |
| Q924C1 | Exportin-5 | Xpo5 |
| Q9QY30 | Bile salt export pump | Abcb11 |
| Q9D8B4 | NADH dehydrogenase [ubiquinone] 1 alpha subcomplex subunit 11 | Ndufa11 |
| P24472 | Glutathione S-transferase A4 | Gsta4 |
| P07309 | Transthyretin | Ttr |
| Q99LT0 | Protein dpy-30 homolog | Dpy30 |
| Q62264 | Thyroid hormone-inducible hepatic protein | Thrsp |
| | **Unique Proteins to C57BL6 (Low-fat): 120** | |
| P01869 | Ig gamma-1 chain C region, membrane-bound form | Ighg1 |
| Q8VCN5 | Cystathionine gamma-lyase | Cth |
| Q80W21 | Glutathione S-transferase Mu 7 | Gstm7 |
| Q64374 | Regucalcin | Rgn |
| P00158 | Cytochrome b | Mt-Cyb |
| Q63836 | Selenium-binding protein 2 | Selenbp2 |
| O35423 | Serine--pyruvate aminotransferase, mitochondrial | Agxt |
| Q91WS0 | CDGSH iron-sulfur domain-containing protein 1 | Cisd1 |
| Q9DCP2 | Sodium-coupled neutral amino acid transporter 3 | Slc38a3 |
| Q8BTY1 | Kynurenine--oxoglutarate transaminase 1 | Kyat1 |
| P19157 | Glutathione S-transferase P 1 | Gstp1 |
| Q9WVM8 | Kynurenine/alpha-aminoadipate aminotransferase, mitochondrial | Aadat |
| P56654 | Cytochrome P450 2C37 | Cyp2c37 |
| Q91X77 | Cytochrome P450 2C50 | Cyp2c50 |
| Q9QXZ6 | Solute carrier organic anion transporter family member 1A1 | Slco1a1 |
| Q91W64 | Cytochrome P450 2C70 | Cyp2c70 |
| Q9DBW0 | Cytochrome P450 4V2 | Cyp4v2 |
| Q8VE09 | Tetratricopeptide repeat protein 39C | Ttc39c |

| | | |
|---|---|---|
| Q91ZI0 | Cadherin EGF LAG seven-pass G-type receptor 3 | Celsr3 |
| Q9JHI5 | Isovaleryl-CoA dehydrogenase, mitochondrial | Ivd |
| Q9D906 | Ubiquitin-like modifier-activating enzyme ATG7 | Atg7 |
| Q8CIF4 | Biotinidase | Btd |
| Q8BLN5 | Lanosterol synthase | Lss |
| Q8BH35 | Complement component C8 beta chain | C8b |
| P61922 | 4-aminobutyrate aminotransferase, mitochondrial | Abat |
| Q283N4 | 2-oxo-4-hydroxy-4-carboxy-5-ureidoimidazoline decarboxylase | Urad |
| E9Q5K4 | Cytochrome P450 2C44 | Cyp2c23 |
| Q9DCT1 | 1,5-anhydro-D-fructose reductase | Akr1e2 |
| Q5SGK3 | Aldehyde oxidase 2 | Aox2 |
| Q6ZQK0 | Condensin-2 complex subunit D3 | Ncapd3 |
| Q05816 | Fatty acid-binding protein 5 | Fabp5 |
| Q8C0L9 | Glycerophosphocholine phosphodiesterase GPCPD1 | Gpcpd1 |
| Q5FW60 | Major urinary protein 20 | Mup20 |
| Q91VA0 | Acyl-coenzyme A synthetase ACSM1, mitochondrial | Acsm1 |
| Q60991 | Cytochrome P450 7B1 | Cyp7b1 |
| Q9D0J8 | Parathymosin | Ptms |
| Q63880 | Carboxylesterase 3A | Ces3a |
| O54782 | Epididymis-specific alpha-mannosidase | Man2b2 |
| Q8BZB2 | Phosphopantothenoylcysteine decarboxylase | Ppcdc |
| Q8QZR1 | Tyrosine aminotransferase | Tat |
| Q9JHU9 | Inositol-3-phosphate synthase 1 | Isyna1 |
| P01887 | Beta-2-microglobulin | B2m |
| Q8VCW8 | Medium-chain acyl-CoA ligase ACSF2, mitochondrial | Acsf2 |
| P56135 | ATP synthase subunit f, mitochondrial | Atp5mf |
| P42703 | Leukemia inhibitory factor receptor | Lifr |
| Q80XI6 | Mitogen-activated protein kinase kinase kinase 11 | Map3k11 |
| Q01768 | Nucleoside diphosphate kinase B | Nme2 |
| Q920A5 | Retinoid-inducible serine carboxypeptidase | Scpep1 |
| Q8QZR3 | Pyrethroid hydrolase Ces2a | Ces2a |
| Q920E5 | Farnesyl pyrophosphate synthase | Fdps |
| Q9WU19 | Hydroxyacid oxidase 1 | Hao1 |
| P00186 | Cytochrome P450 1A2 | Cyp1a2 |
| Q99PG0 | Arylacetamide deacetylase | Aadac |
| Q9WUR9 | Adenylate kinase 4, mitochondrial | Ak4 |
| A6BLY7 | Keratin, type I cytoskeletal 28 | Krt28 |
| Q8VC97 | Beta-ureidopropionase | Upb1 |
| O35943 | Frataxin, mitochondrial | Fxn |
| P28666 | Murinoglobulin-2 | Mug2 |
| P50429 | Arylsulfatase B | Arsb |
| Q8JZZ0 | UDP-glucuronosyltransferase 3A2 | Ugt3a2 |
| P58044 | Isopentenyl-diphosphate Delta-isomerase 1 | Idi1 |
| Q9DCY0 | Glycine N-acyltransferase-like protein Keg1 | Keg1 |
| Q9ERY9 | Ergosterol biosynthetic protein 28 homolog | Erg28 |

| | | |
|---|---|---|
| Q6XVG2 | Cytochrome P450 2C54 | Cyp2c54 |
| O08600 | Endonuclease G, mitochondrial | Endog |
| P11589 | Major urinary protein 2 | Mup2 |
| Q9CZP5 | Mitochondrial chaperone BCS1 | Bcs1l |
| P07759 | Serine protease inhibitor A3K | Serpina3k |
| Q9R1J0 | Sterol-4-alpha-carboxylate 3-dehydrogenase, decarboxylating | Nsdhl |
| Q62452 | UDP-glucuronosyltransferase 1A9 | Ugt1a9 |
| Q8VCG4 | Complement component C8 gamma chain | C8g |
| Q8K0C4 | Lanosterol 14-alpha demethylase | Cyp51a1 |
| Q8K0L9 | Zinc finger and BTB domain-containing protein 20 | Zbtb20 |
| G3X982 | Aldehyde oxidase 3 | Aox3 |
| P01878 | Ig alpha chain C region | n/a |
| Q571F8 | Glutaminase liver isoform, mitochondrial | Gls2 |
| Q9D8B6 | Protein FAM210B, mitochondrial | Fam210b |
| Q9CRA4 | Methylsterol monooxygenase 1 | Msmo1 |
| Q61694 | NADPH-dependent 3-keto-steroid reductase Hsd3b5 | Hsd3b5 |
| Q91WN4 | Kynurenine 3-monooxygenase | Kmo |
| Q61176 | Arginase-1 | Arg1 |
| Q9R008 | Mevalonate kinase | Mvk |
| P33267 | Cytochrome P450 2F2 | Cyp2f2 |
| Q71KT5 | Delta(14)-sterol reductase TM7SF2 | Tm7sf2 |
| Q8K2I4 | Beta-mannosidase | Manba |
| Q9DBE0 | Cysteine sulfinic acid decarboxylase | Csad |
| Q9Z2V4 | Phosphoenolpyruvate carboxykinase, cytosolic [GTP] | Pck1 |
| P17439 | Lysosomal acid glucosylceramidase | Gba |
| Q9DB29 | Isoamyl acetate-hydrolyzing esterase 1 homolog | Iah1 |
| Q9Z1R3 | Apolipoprotein M | Apom |
| Q6IFZ6 | Keratin, type II cytoskeletal 1b | Krt77 |
| P49935 | Pro-cathepsin H | Ctsh |
| P43024 | Cytochrome c oxidase subunit 6A1, mitochondrial | Cox6a1 |
| Q9R013 | Cathepsin F | Ctsf |
| Q91WG0 | Acylcarnitine hydrolase | Ces2c |
| Q9QWR8 | Alpha-N-acetylgalactosaminidase | Naga |
| Q9QXF8 | Glycine N-methyltransferase | Gnmt |
| P11725 | Ornithine transcarbamylase, mitochondrial | Otc |
| O88668 | Protein CREG1 | Creg1 |
| Q8C196 | Carbamoyl-phosphate synthase [ammonia], mitochondrial | Cps1 |
| Q63886 | UDP-glucuronosyltransferase 1A1 | Ugt1a1 |
| Q99K67 | Alpha-aminoadipic semialdehyde synthase, mitochondrial | Aass |
| Q01279 | Epidermal growth factor receptor | Egfr |
| Q99P30 | Peroxisomal coenzyme A diphosphatase NUDT7 | Nudt7 |
| P52843 | Sulfotransferase 2A1 | Sult2a1 |
| Q8VCU1 | Carboxylesterase 3B | Ces3b |
| Q80W22 | Threonine synthase-like 2 | Thnsl2 |
| P05366 | Serum amyloid A-1 protein | Saa1 |

| | | |
|---|---|---|
| Q9D0S9 | Histidine triad nucleotide-binding protein 2, mitochondrial | Hint2 |
| P00688 | Pancreatic alpha-amylase | Amy2 |
| Q9JLF6 | Protein-glutamine gamma-glutamyltransferase K | Tgm1 |
| Q99LB7 | Sarcosine dehydrogenase, mitochondrial | Sardh |
| Q9EP72 | ER membrane protein complex subunit 7 | Emc7 |
| Q9DCG2 | CD302 antigen | Cd302 |
| Q00898 | Alpha-1-antitrypsin 1-5 | Serpina1e |
| P16331 | Phenylalanine-4-hydroxylase | Pah |
| P52825 | Carnitine O-palmitoyltransferase 2, mitochondrial | Cpt2 |
| Q9D273 | Corrinoid adenosyltransferase | Mmab |
| P16460 | Argininosuccinate synthase | Ass1 |
| Q91XF0 | Pyridoxine-5'-phosphate oxidase | Pnpo |

The liver samples (from 5 mice in each group) were individually assessed by TMT-based differential proteomic expression and data were merged to get averages. Protein FDR Confidence for all proteins was $\leq$2%. FDR: False Discovery Rate. These altered proteins were upregulated compared to the high-fat control group (MS-NASH mice on a high-fat diet) with a 2-fold-change threshold. These data (Venn diagrams) are shown in Figure 3A.
