## Supplemental Table 3 for "Liver Protein Expression in Nash Mice on a High-Fat Diet"

**Supplement Table 3. Common and unique downregulated proteins (at 2-fold change threshold)**

| Accession | Protein names | Gene Name |
|---|---|---|
| | **Common Proteins among Aquamin, OCA and C57BL6 (low-fat): 1** | |
| Q6NZL6 | Tonsoku-like protein | Tonsl |
| | **Common Proteins between OCA and C57BL6 (low-fat): 2** | |
| Q6A065 | Centrosomal protein of 170 kDa | Cep170 |
| P16045 | Galectin-1 | Lgals1 |
| | **Common Proteins between Aquamin and C57BL6 (low-fat): 1** | |
| Q921W4 | Quinone oxidoreductase-like protein 1 | Cryzl1 |
| | **Common Proteins between Aquamin and OCA: 1** | |
| Q8C4X7 | Membrane integral NOTCH2-associated receptor 2 | Minar2 |
| | **Unique Proteins to Aquamin: 2** | |
| Q6XVG2 | Cytochrome P450 2C54 | Cyp2c54 |
| Q00898 | Alpha-1-antitrypsin 1-5 | Serpina1e |
| | **Unique Proteins to OCA: 4** | |
| P29758 | Ornithine aminotransferase, mitochondrial | Oat |
| P11589 | Major urinary protein 2 | Mup2 |
| O88962 | 7-alpha-hydroxycholest-4-en-3-one 12-alpha-hydroxylase | Cyp8b1 |
| P05366 | Serum amyloid A-1 protein | Saa1 |
| | **Unique Proteins to C57BL6 (Low-fat): 122** | |
| P62806 | Histone H4 | H4c1 |
| A2A995 | FYN-binding protein 2 | Fyb2 |
| Q9R0Y5 | Adenylate kinase isoenzyme 1 | Ak1 |
| Q8K2K6 | Arf-GAP domain and FG repeat-containing protein 1 | Agfg1 |
| Q62245 | Son of sevenless homolog 1 | Sos1 |
| Q3U4I7 | Pyridine nucleotide-disulfide oxidoreductase domain-containing protein 2 | Pyroxd2 |
| Q8CC35 | Synaptopodin | Synpo |
| Q8K4G5 | Actin-binding LIM protein 1 | Ablim1 |
| P97819 | 85/88 kDa calcium-independent phospholipase A2 | Pla2g6 |
| Q8VCR2 | 17-beta-hydroxysteroid dehydrogenase 13 | Hsd17b13 |
| Q61285 | ATP-binding cassette sub-family D member 2 | Abcd2 |
| Q2NL51 | Glycogen synthase kinase-3 alpha | Gsk3a |
| Q6NZF1 | Zinc finger CCCH domain-containing protein 11A | Zc3h11a |
| Q64475 | Histone H2B type 1-B | H2bc3 |
| Q9DBS9 | Oxysterol-binding protein-related protein 3 | Osbpl3 |
| Q8BK63 | Casein kinase I isoform alpha | Csnk1a1 |
| Q810Z1 | Epididymal-specific lipocalin-10 | Lcn10 |
| P16110 | Galectin-3 | Lgals3 |
| Q62417 | Sorbin and SH3 domain-containing protein 1 | Sorbs1 |
| Q8R4R6 | Nucleoporin NUP35 | Nup35 |
| Q8VC49 | Interferon alpha-inducible protein 27-like protein 2B | Ifi27l2b |
| Q69Z37 | Sterile alpha motif domain-containing protein 9-like | Samd9l |
| Q8BJF9 | Charged multivesicular body protein 2b | Chmp2b |
| P12790 | Cytochrome P450 2B9 | Cyp2b9 |

| | | |
|---|---|---|
| Q8K2F0 | Bromodomain-containing protein 3 | Brd3 |
| Q60875 | Rho guanine nucleotide exchange factor 2 | Arhgef2 |
| Q8CGP5 | Histone H2A type 1-F | Hist1h2af |
| Q91V92 | ATP-citrate synthase | Acly |
| Q8CDN9 | Leucine-rich repeat-containing protein 9 | Lrrc9 |
| O89053 | Coronin-1A | Coro1a |
| P19096 | Fatty acid synthase | Fasn |
| Q99LJ0 | CTTNBP2 N-terminal-like protein | Cttnbp2nl |
| Q14DH7 | Acyl-CoA synthetase short-chain family member 3, mitochondrial | Acss3 |
| Q8CIB6 | Transmembrane protein 230 | Tmem230 |
| Q9R257 | Heme-binding protein 1 | Hebp1 |
| Q9DBM2 | Peroxisomal bifunctional enzyme | Ehhadh |
| Q9Z2H5 | Band 4.1-like protein 1 | Epb41l1 |
| Q9R1Q7 | Proteolipid protein 2 | Plp2 |
| Q3TJD7 | PDZ and LIM domain protein 7 | Pdlim7 |
| Q9D0P0 | Emopamil-binding protein-like | Ebpl |
| Q9Z2A7 | Diacylglycerol O-acyltransferase 1 | Dgat1 |
| P04441 | H-2 class II histocompatibility antigen gamma chain | Cd74 |
| Q9CQ19 | Myosin regulatory light polypeptide 9 | Myl9 |
| P55050 | Fatty acid-binding protein, intestinal | Fabp2 |
| Q8R1N4 | NudC domain-containing protein 3 | Nudcd3 |
| O88492 | Perilipin-4 | Plin4 |
| Q99MQ5 | Collagen alpha-1(XXV) chain | Col25a1 |
| P56656 | Cytochrome P450 2C39 | Cyp2c39 |
| Q9D939 | Sulfotransferase 1C2 | Sult1c2 |
| Q91YR9 | Prostaglandin reductase 1 | Ptgr1 |
| Q9QZC8 | Protein ABHD1 | Abhd1 |
| Q9D312 | Keratin, type I cytoskeletal 20 | Krt20 |
| O54931 | A-kinase anchor protein 2 | Akap2 |
| P84244 | Histone H3.3 | H3-3a |
| Q6GSS7 | Histone H2A type 2-A | Hist2h2aa1 |
| Q99N42 | Thymidine phosphorylase | Tymp |
| P97315 | Cysteine and glycine-rich protein 1 | Csrp1 |
| Q9D964 | Glycine amidinotransferase, mitochondrial | Gatm |
| Q6Y7W8 | GRB10-interacting GYF protein 2 | Gigyf2 |
| P06728 | Apolipoprotein A-IV | Apoa4 |
| P35576 | Glucose-6-phosphatase catalytic subunit 1 | G6pc1 |
| Q5SWU9 | Acetyl-CoA carboxylase 1 | Acaca |
| P05480 | Neuronal proto-oncogene tyrosine-protein kinase Src | Src |
| Q9QYR9 | Acyl-coenzyme A thioesterase 2, mitochondrial | Acot2 |
| Q6PCN7 | Helicase-like transcription factor | Hltf |
| Q61753 | D-3-phosphoglycerate dehydrogenase | Phgdh |
| P13516 | Acyl-CoA desaturase 1 | Scd1 |
| P17095 | High mobility group protein HMG-I/HMG-Y | Hmga1 |
| P55088 | Aquaporin-4 | Aqp4 |

| | | |
|---|---|---|
| Q6ZPJ0 | Testis-expressed protein 2 | Tex2 |
| Q3TFD2 | Lysophosphatidylcholine acyltransferase 1 | Lpcat1 |
| Q9JJY3 | Sphingomyelin phosphodiesterase 3 | Smpd3 |
| Q62523 | Zyxin | Zyx |
| Q9Z211 | Peroxisomal membrane protein 11A | Pex11a |
| P27661 | Histone H2AX | H2ax |
| Q80VJ3 | 2'-deoxynucleoside 5'-phosphate N-hydrolase 1 | Dnph1 |
| Q8K2I1 | Protein farnesyltransferase subunit beta | Fntb |
| Q3UHK8 | Trinucleotide repeat-containing gene 6A protein | Tnrc6a |
| Q9Z1W9 | STE20/SPS1-related proline-alanine-rich protein kinase | Stk39 |
| Q3THW5 | Histone H2A.V | H2az2 |
| Q80X19 | Collagen alpha-1(XIV) chain | Col14a1 |
| P19973 | Lymphocyte-specific protein 1 | Lsp1 |
| P70429 | Ena/VASP-like protein | Evl |
| Q99P72 | Reticulon-4 | Rtn4 |
| Q9Z0G0 | PDZ domain-containing protein GIPC1 | Gipc1 |
| O35855 | Branched-chain-amino-acid aminotransferase, mitochondrial | Bcat2 |
| P84228 | Histone H3.2 | H3c2 |
| Q9QYH6 | Melanoma-associated antigen D1 | Maged1 |
| P52623 | Uridine-cytidine kinase 1 | Uck1 |
| P27546 | Microtubule-associated protein 4 | Map4 |
| P48410 | ATP-binding cassette sub-family D member 1 | Abcd1 |
| Q91XC8 | Death-associated protein 1 | Dap |
| P43276 | Histone H1.5 | H1-5 |
| P0C7L0 | WAS/WASL-interacting protein family member 3 | Wipf3 |
| O35678 | Monoglyceride lipase | Mgll |
| P08207 | Protein S100-A10 | S100a10 |
| Q8BHI7 | Elongation of very long chain fatty acids protein 5 | Elovl5 |
| Q3THE2 | Myosin regulatory light chain 12B | Myl12b |
| Q8VC30 | Triokinase/FMN cyclase | Tkfc |
| P68433 | Histone H3.1 | H3c1 |
| Q69ZH9 | Rho GTPase-activating protein 23 | Arhgap23 |
| Q62266 | Cornifin-A | Sprr1a |
| P10107 | Annexin A1 | Anxa1 |
| Q3UMT1 | Protein phosphatase 1 regulatory subunit 12C | Ppp1r12c |
| Q9D0R8 | Protein LSM12 homolog | Lsm12 |
| Q05020 | Apolipoprotein C-II | Apoc2 |
| Q91XV3 | Brain acid soluble protein 1 | Basp1 |
| Q8VHQ9 | Acyl-coenzyme A thioesterase 11 | Acot11 |
| P07356 | Annexin A2 | Anxa2 |
| Q8CFE4 | SCY1-like protein 2 | Scyl2 |
| P43883 | Perilipin-2 | Plin2 |
| Q9CQE5 | Regulator of G-protein signaling 10 | Rgs10 |
| P35459 | Lymphocyte antigen 6D | Ly6d |
| A2A7S8 | Uncharacterized protein KIAA1522 | Kiaa1522 |

| | | |
|---|---|---|
| P70296 | Phosphatidylethanolamine-binding protein 1 | Pebp1 |
| Q6R2P8 | Endonuclease 8-like 2 | Neil2 |
| P09813 | Apolipoprotein A-II | Apoa2 |
| P10637 | Microtubule-associated protein tau | Mapt |
| Q7TSI3 | Serine/threonine-protein phosphatase 6 regulatory subunit 1 | Ppp6r1 |
| P56655 | Cytochrome P450 2C38 | Cyp2c38 |
| P19324 | Serpin H1 | Serpinh1 |
| Q3V0K9 | Plastin-1 | Pls1 |

The liver samples (from 5 mice in each group) were individually assessed by TMT-based differential proteomic expression and data were merged to get averages. Protein FDR Confidence for all proteins was $\leq$2%. FDR: False Discovery Rate. These altered proteins were downregulated compared to the high-fat control group (MS-NASH mice on a high-fat diet) with a 2-fold-change threshold. These data (Venn diagrams) are shown in Figure 3B.
