## Supplemental Table 4 for "Liver Protein Expression in Nash Mice on a High-Fat Diet"

**Supplement Table 4. Significantly altered (Upregulated) proteins with Aquamin in high-fat mice**

| Proteins | Genes | MS-NASH | | C57BL6 |
|---|---|---|---|---|
| | | Aquamin | OCA | (Low-Fat) |
| UDP-N-acetylhexosamine pyrophosphorylase-like protein 1 | Uap1l1 | *1.9±0.8 | 0.9±0.4 | 1.1±0.2 |
| Biglycan | Bgn | *1.7±0.5 | 1.5±0.6 | 0.8±0.3 |
| Tax1-binding protein 3 | Tax1bp3 | *1.6±0.5 | 1.6±1.1 | 1.1±0.5 |
| Succinyl-CoA:3-ketoacid coenzyme A transferase 1, mitochondrial | Oxct1 | *1.6±0.3 | *1.4±0.2 | 1.2±0.4 |
| Glycogen phosphorylase, brain form | Pygb | *1.5±0.3 | 0.9±0.2 | 0.6±0.1 |
| Acyl-coenzyme A thioesterase 9, mitochondrial | Acot9 | *1.5±0.5 | 1.3±0.6 | 0.8±0.6 |
| CD166 antigen | Alcam | *1.5±0.4 | 1.3±0.5 | *1.2±0.2 |
| Erythrocyte band 7 integral membrane protein | Stom | *1.5±0.4 | 1.2±0.5 | 0.9±0.5 |
| RNA-binding protein 3 | Rbm3 | *1.5±0.3 | *1.4±0.4 | 1.2±0.3 |
| Angiotensinogen | Agt | *1.5±0.4 | 1.1±0.3 | *1.5±0.4 |
| Apolipoprotein A-I | Apoa1 | *1.5±0.3 | 1.0±0.3 | 0.8±0.2 |
| Heparin cofactor 2 | Serpind1 | *1.5±0.4 | 1.2±0.6 | 1.1±0.4 |
| Endonuclease/exonuclease/phosphatase family domain-containing protein 1 | Eepd1 | *1.4±0.4 | 1.4±0.5 | 0.9±0.4 |
| Galectin-3 | Lgals3 | *1.4±0.4 | 0.8±0.3 | 0.4±0.2 |
| Sterile alpha motif domain-containing protein 9-like | Samd9l | *1.4±0.1 | 1.2±0.2 | 0.4±0.1 |
| Annexin A5 | Anxa5 | *1.4±0.3 | 1.0±0.3 | 0.7±0.1 |
| Myoferlin | Myof | *1.4±0.1 | 1.0±0.2 | 0.6±0.2 |
| Charged multivesicular body protein 2b | Chmp2b | *1.4±0.2 | 1.1±0.2 | 0.4±0.2 |
| ATP-dependent 6-phosphofructokinase, platelet type | Pfkp | *1.4±0.3 | 0.9±0.3 | 0.6±0.3 |
| 55 kDa erythrocyte membrane protein | Mpp1 | *1.4±0.3 | 1.1±0.2 | 0.9±0.3 |
| Protein S100-A10 | S100a10 | *1.4±0.3 | 0.9±0.4 | 0.5±0.1 |
| Protein unc-119 homolog B | Unc119b | *1.3±0.2 | 1.1±0.2 | 1.1±0.5 |
| Glucose-6-phosphate isomerase | Gpi1 | *1.3±0.3 | *1.4±0.4 | 0.7±0.1 |
| Major vault protein | Mvp | *1.3±0.3 | 1.0±0.1 | 0.9±0.1 |
| Hexokinase-4 | Gck | *1.3±0.3 | *1.3±0.1 | 0.6±0.2 |
| Ankyrin repeat and SOCS box protein 13 | Asb13 | *1.3±0.1 | *1.5±0.2 | 0.8±0.1 |
| Serine/threonine-protein kinase TBK1 | Tbk1 | *1.3±0.0 | 1.2±0.3 | 1.3±0.2 |
| Annexin A2 | Anxa2 | *1.3±0.2 | 0.8±0.3 | 0.4±0.1 |
| Glutamate--cysteine ligase catalytic subunit | Gclc | *1.3±0.2 | 1.1±0.1 | 0.7±0.1 |
| DnaJ homolog subfamily C member 8 | Dnajc8 | *1.3±0.1 | 1.1±0.2 | 1.6±0.9 |
| Coactosin-like protein | Cotl1 | *1.3±0.2 | 1.0±0.2 | 0.5±0.2 |
| RNA polymerase II-associated factor 1 homolog | Paf1 | *1.3±0.1 | 1.2±0.2 | 1.1±0.1 |
| Phosphopantothenoylcysteine decarboxylase | Ppcdc | *1.3±0.1 | 1.4±0.4 | 2.5±2.1 |

| | | | | |
|---|---|---|---|---|
| Cytochrome c oxidase subunit 7A1, mitochondrial | Cox7a1 | *1.3±0.1 | 1.0±0.1 | 0.8±0.2 |
| Carbonyl reductase [NADPH] 1 | Cbr1 | *1.3±0.2 | *1.5±0.2 | 1.1±0.4 |
| Rho GDP-dissociation inhibitor 1 | Arhgdia | *1.3±0.0 | 1.1±0.2 | 1.0±0.1 |
| Alpha-2-HS-glycoprotein | Ahsg | *1.3±0.2 | 1.1±0.1 | 1.0±0.3 |
| COMM domain-containing protein 4 | Commd4 | *1.3±0.1 | 1.1±0.4 | 0.8±0.1 |
| Xaa-Pro dipeptidase | Pepd | *1.3±0.2 | 1.2±0.2 | 0.7±0.1 |
| Fascin | Fscn1 | *1.3±0.2 | 0.9±0.2 | 0.9±0.3 |
| Exopolyphosphatase PRUNE1 | Prune1 | *1.3±0.2 | 1.1±0.2 | 0.6±0.1 |
| Afamin | Afm | *1.3±0.2 | 1.1±0.1 | 0.9±0.4 |
| C4b-binding protein | C4bp | *1.3±0.2 | 0.8±0.2 | 0.9±0.2 |
| Transketolase | Tkt | *1.3±0.2 | *1.4±0.3 | 1.0±0.2 |
| Pyruvate kinase PKLR | Pklr | *1.3±0.2 | *1.5±0.2 | 0.8±0.2 |
| Myosin light polypeptide 6 | Myl6 | *1.3±0.2 | 1.1±0.5 | 0.6±0.3 |
| Glutathione S-transferase Mu 3 | Gstm3 | *1.3±0.2 | 1.2±0.4 | 0.8±0.3 |
| WD repeat-containing protein 1 | Wdr1 | *1.3±0.2 | 1.1±0.3 | 1.0±0.3 |
| Probable rRNA-processing protein EBP2 | Ebna1bp2 | *1.3±0.2 | *1.2±0.1 | 1.0±0.0 |
| Profilin-1 | Pfn1 | *1.2±0.2 | 1.1±0.2 | 1.0±0.1 |
| Probable tRNA N6-adenosine threonylcarbamoyltransferase | Osgep | *1.2±0.2 | 1.0±0.2 | 0.8±0.2 |
| EKC/KEOPS complex subunit Tp53rk | Trp53rk | *1.2±0.1 | 1.1±0.1 | 1.4±0.7 |
| Phospholipid scramblase 1 | Plscr1 | *1.2±0.1 | *1.2±0.1 | 0.8±0.2 |
| Protein LSM14 homolog A | Lsm14a | *1.2±0.2 | *1.2±0.2 | 0.7±0.2 |
| Pyrroline-5-carboxylate reductase 3 | Pycrl | *1.2±0.2 | 1.0±0.2 | *1.4±0.3 |
| Glutathione S-transferase A1 | Gsta1 | *1.2±0.1 | *1.5±0.1 | 0.5±0.1 |
| Striatin-3 | Strn3 | *1.2±0.1 | *1.3±0.1 | 1.0±0.2 |
| Glycogen synthase kinase-3 alpha | Gsk3a | *1.2±0.2 | 1.2±0.4 | 0.5±0.3 |
| Serum albumin | Alb | *1.2±0.1 | 1.1±0.2 | 0.8±0.2 |
| Fibulin-1 | Fbln1 | *1.2±0.1 | 0.9±0.1 | 0.9±0.3 |
| Small ubiquitin-related modifier 1 | Sumo1 | *1.2±0.1 | 1.0±0.1 | 0.9±0.1 |
| DnaJ homolog subfamily C member 10 | Dnajc10 | *1.2±0.1 | 1.0±0.2 | 1.0±0.2 |
| Cyclin-dependent kinase 6 | Cdk6 | *1.2±0.2 | 1.2±0.2 | 0.8±0.4 |
| Ragulator complex protein LAMTOR5 | Lamtor5 | *1.2±0.0 | 1.0±0.1 | *1.3±0.1 |
| Microtubule-associated protein RP/EB family member 3 | Mapre3 | *1.2±0.2 | 0.9±0.2 | 0.9±0.1 |
| Abscission/NoCut checkpoint regulator | Zfyve19 | *1.2±0.1 | 1.1±0.3 | 0.8±0.1 |
| Rho guanine nucleotide exchange factor 7 | Arhgef7 | *1.2±0.1 | 1.1±0.2 | 0.9±0.1 |
| Synaptic vesicle membrane protein VAT-1 homolog | Vat1 | *1.2±0.1 | 1.0±0.2 | 1.0±0.2 |
| Carbonyl reductase [NADPH] 3 | Cbr3 | *1.2±0.2 | 1.1±0.3 | 0.8±0.3 |
| m7GpppX diphosphatase | Dcps | *1.2±0.2 | *1.2±0.2 | *1.5±0.2 |

| | | | | |
|---|---|---|---|---|
| Pre-rRNA-processing protein TSR2 homolog | Tsr2 | *1.2±0.2 | 1.1±0.1 | 0.8±0.3 |
| Adenylyl cyclase-associated protein 1 | Cap1 | *1.2±0.1 | 1.0±0.1 | 0.8±0.1 |
| Endothelin-converting enzyme 1 | Ece1 | *1.2±0.1 | 1.1±0.2 | 1.1±0.2 |
| Cytoplasmic FMR1-interacting protein 1 | Cyfip1 | *1.2±0.2 | 1.0±0.1 | 0.8±0.2 |
| H-2 class I histocompatibility antigen, Q10 alpha chain | H2-Q10 | *1.2±0.1 | 0.9±0.1 | 0.6±0.0 |
| Nuclear autoantigenic sperm protein | Nasp | *1.2±0.1 | 1.0±0.1 | 1.2±0.4 |
| Protein C10 | Grcc10 | *1.2±0.2 | 1.4±0.9 | 1.6±0.9 |
| Zinc-alpha-2-glycoprotein | Azgp1 | *1.2±0.1 | 0.9±0.1 | 1.0±0.1 |
| Chromobox protein homolog 1 | Cbx1 | *1.2±0.2 | 1.1±0.1 | 0.8±0.2 |
| CCR4-NOT transcription complex subunit 10 | Cnot10 | *1.2±0.1 | 1.1±0.1 | 1.1±0.3 |
| TBC1 domain family member 22A | Tbc1d22a | *1.2±0.1 | 1.1±0.3 | 0.6±0.5 |
| Bis(5'-nucleosyl)-tetraphosphatase [asymmetrical] | Nudt2 | *1.2±0.2 | 1.1±0.1 | 1.0±0.2 |
| Paired amphipathic helix protein Sin3a | Sin3a | *1.2±0.1 | 1.1±0.2 | 2.0±2.1 |
| Alpha-actinin-4 | Actn4 | *1.2±0.2 | 1.0±0.2 | 0.9±0.2 |
| Ubiquitin-like modifier-activating enzyme 5 | Uba5 | *1.2±0.2 | 1.1±0.1 | 0.7±0.3 |
| Solute carrier family 23 member 1 | Slc23a1 | *1.2±0.1 | 1.0±0.1 | 0.8±0.0 |
| Leukocyte surface antigen CD47 | Cd47 | *1.2±0.1 | 1.0±0.1 | *1.3±0.3 |
| Ubiquitin thioesterase OTUB1 | Otub1 | *1.2±0.1 | 1.1±0.1 | 0.8±0.2 |
| Craniofacial development protein 1 | Cfdp1 | *1.2±0.2 | 1.1±0.1 | 0.6±0.1 |
| Osteoclast-stimulating factor 1 | Ostf1 | *1.2±0.2 | 1.0±0.1 | 0.7±0.1 |
| Protein FAM107B | Fam107b | *1.2±0.1 | 1.0±0.1 | 0.9±0.2 |
| ATP-dependent 6-phosphofructokinase, liver type | Pfkl | *1.2±0.1 | *1.2±0.1 | 0.7±0.2 |
| 39S ribosomal protein L55, mitochondrial | Mrpl55 | *1.2±0.0 | 1.0±0.0 | 1.2±0.2 |
| Heterogeneous nuclear ribonucleoprotein U-like protein 1 | Hnrnpul1 | *1.2±0.1 | 1.1±0.1 | 1.0±0.1 |
| Protein kinase C and casein kinase substrate in neurons protein 2 | Pacsin2 | *1.2±0.1 | 1.1±0.1 | 1.0±0.1 |
| ATPase inhibitor, mitochondrial | Atpif1 | *1.2±0.1 | 0.9±0.2 | 0.8±0.1 |
| Alpha-enolase | Eno1 | *1.2±0.1 | *1.2±0.2 | 0.7±0.1 |
| Alpha-actinin-1 | Actn1 | *1.2±0.1 | 1.0±0.1 | 0.8±0.1 |
| COMM domain-containing protein 1 | Commd1 | *1.2±0.1 | 0.9±0.2 | 1.2±0.3 |
| High mobility group protein B1 | Hmgb1 | *1.2±0.1 | 1.1±0.1 | 1.1±0.2 |
| Gamma-glutamylcyclotransferase | Ggct | *1.2±0.1 | 1.0±0.1 | 0.8±0.2 |
| 5'-AMP-activated protein kinase catalytic subunit alpha-2 | Prkaa2 | *1.2±0.1 | *1.4±0.2 | 0.7±0.1 |
| CTP synthase 1 | Ctps | *1.2±0.1 | 0.9±0.2 | 0.8±0.2 |
| UBX domain-containing protein 1 | Ubxn1 | *1.2±0.1 | 1.0±0.2 | 1.1±0.3 |
| DNA-(apurinic or apyrimidinic site) lyase | Apex1 | *1.2±0.1 | 1.1±0.1 | *1.3±0.1 |
| Serine/threonine-protein phosphatase 2A catalytic subunit alpha isoform | Ppp2ca | *1.2±0.1 | 1.1±0.1 | 1.1±0.1 |
| Beta-2-glycoprotein 1 | Apoh | *1.2±0.1 | 0.9±0.1 | 0.8±0.2 |

| | | | | |
|---|---|---|---|---|
| Interferon-induced, double-stranded RNA-activated protein kinase | Eif2ak2 | *1.2±0.1 | 1.0±0.2 | 1.2±0.2 |
| RNA-binding protein FUS | Fus | *1.2±0.1 | 1.0±0.2 | 1.1±0.2 |
| Mini-chromosome maintenance complex-binding protein | Mcmbp | *1.1±0.1 | 1.2±0.3 | 1.0±0.2 |
| Receptor-interacting serine/threonine-protein kinase 1 | Ripk1 | *1.1±0.1 | 1.0±0.1 | 1.1±0.3 |
| BMP-2-inducible protein kinase | Bmp2k | *1.1±0.1 | 1.2±0.3 | 0.7±0.3 |
| FACT complex subunit SPT16 | Supt16h | *1.1±0.1 | 1.0±0.1 | 1.1±0.2 |
| U6 snRNA-associated Sm-like protein LSm6 | Lsm6 | *1.1±0.1 | 1.0±0.2 | 1.1±0.2 |
| Protein S100-A4 | S100a4 | *1.1±0.0 | 0.9±0.7 | 1.0±0.4 |
| pre-mRNA 3' end processing protein WDR33 | Wdr33 | *1.1±0.0 | 1.1±0.1 | 1.5±0.7 |
| High affinity immunoglobulin gamma Fc receptor I | Fcgr1 | *1.1±0.1 | 1.0±0.3 | 0.5±0.1 |
| U6 snRNA-associated Sm-like protein LSm3 | Lsm3 | *1.1±0.1 | 0.9±0.1 | 1.1±0.2 |
| Hepatocyte nuclear factor 4-alpha | Hnf4a | *1.1±0.1 | *1.2±0.1 | 1.2±0.4 |
| Ras GTPase-activating protein-binding protein 1 | G3bp1 | *1.1±0.1 | 1.0±0.1 | 0.9±0.1 |
| Brain-specific angiogenesis inhibitor 1-associated protein 2 | Baiap2 | *1.1±0.1 | *1.3±0.2 | *1.3±0.2 |
| Neutral cholesterol ester hydrolase 1 | Nceh1 | *1.1±0.1 | 0.7±0.2 | 0.8±0.1 |
| Heterogeneous nuclear ribonucleoprotein U | Hnrnpu | *1.1±0.1 | 1.1±0.1 | 0.9±0.1 |
| Sorting nexin-4 | Nceh1 | *1.1±0.1 | 0.9±0.1 | 0.7±0.2 |
| Cell cycle and apoptosis regulator protein 2 | Ccar2 | *1.1±0.1 | 1.0±0.1 | 0.9±0.1 |
| Host cell factor 1 | Hcfc1 | *1.1±0.1 | 1.0±0.2 | 0.8±0.2 |
| UDP-glucuronic acid/UDP-N-acetylgalactosamine transporter | Slc35d1 | *1.1±0.1 | 1.0±0.1 | 0.9±0.5 |
| Fatty acid-binding protein, intestinal | Fabp2 | *1.1±0.1 | *1.2±0.1 | 0.4±0.1 |
| tRNA pseudouridine synthase A | Pus1 | *1.1±0.1 | 1.1±0.2 | 1.0±0.2 |
| Alpha-2-antiplasmin | Serpinf2 | *1.1±0.1 | 1.0±0.1 | 0.9±0.1 |
| Ras-related protein Rab-5C | Rab5c | *1.1±0.1 | 1.0±0.1 | 0.9±0.2 |
| Guanine nucleotide-binding protein G(i) subunit alpha-2 | Gnai2 | *1.1±0.1 | 0.9±0.2 | 0.8±0.2 |
| Heterogeneous nuclear ribonucleoprotein D0 | Hnrnpd | *1.1±0.1 | 1.1±0.2 | *1.1±0.1 |
| Presequence protease, mitochondrial | Pitrm1 | *1.1±0.1 | 1.0±0.1 | 1.0±0.1 |
| 14-3-3 protein epsilon | Ywhae | *1.1±0.1 | *1.1±0.1 | 1.0±0.1 |
| Dynactin subunit 5 | Dctn5 | *1.1±0.0 | 0.9±0.1 | 0.8±0.3 |
| Actin, cytoplasmic 1 | Actb | *1.1±0.1 | 1.0±0.2 | 0.8±0.3 |
| Signal transducer and activator of transcription 5B | Stat5b | *1.1±0.1 | 1.1±0.2 | 0.8±0.2 |
| Phosphatidylinositol transfer protein alpha isoform | Pitpna | *1.1±0.1 | 1.0±0.1 | 0.9±0.1 |
| Septin-2 | Sept2 | *1.1±0.1 | 1.0±0.1 | 0.9±0.1 |
| L-fucose kinase | Fuk | *1.1±0.1 | 1.1±0.2 | 0.8±0.3 |
| 6-phosphogluconate dehydrogenase, decarboxylating | Pgd | *1.1±0.1 | 1.1±0.1 | 0.6±0.1 |
| Wiskott-Aldrich syndrome protein family member 2 | Wasf2 | *1.1±0.1 | 0.9±0.1 | 0.6±0.1 |
| Ubiquitin-conjugating enzyme E2 H | Ube2h | *1.1±0.1 | 1.0±0.1 | 0.9±0.1 |

| | | | | |
|---|---|---|---|---|
| Calcyclin-binding protein | Cacybp | *1.1±0.1 | *1.1±0.1 | 1.0±0.1 |
| BAG family molecular chaperone regulator 1 | Bag1 | *1.1±0.1 | 1.0±0.1 | 0.8±0.1 |
| RNA-binding Raly-like protein | Ralyl | *1.1±0.0 | 0.9±0.1 | 1.1±0.1 |
| Tubulin alpha-1C chain | Tuba1c | *1.1±0.1 | 1.0±0.1 | 0.7±0.2 |
| Pre-mRNA-splicing factor ATP-dependent RNA helicase DHX15 | Dhx15 | *1.1±0.1 | *1.1±0.1 | 1.0±0.2 |
| NEDD8-activating enzyme E1 catalytic subunit | Uba3 | *1.1±0.1 | 1.0±0.1 | 0.8±0.1 |
| Tumor protein D54 | Tpd52l2 | *1.1±0.1 | 1.0±0.1 | 0.9±0.1 |
| Ras-related C3 botulinum toxin substrate 1 | Rac1 | *1.1±0.1 | 1.0±0.1 | 1.0±0.1 |
| Desmoglein-2 | Dsg2 | *1.1±0.1 | 1.1±0.1 | 1.0±0.1 |
| Serine/threonine-protein phosphatase PP1-alpha catalytic subunit | Ppp1ca | *1.1±0.1 | 1.1±0.1 | 1.0±0.1 |
| Paxillin | Pxn | *1.1±0.1 | *1.1±0.0 | 0.8±0.1 |
| Heat shock 70 kDa protein 4 | Hspa4 | *1.1±0.1 | *1.1±0.0 | 1.0±0.1 |
| Hsc70-interacting protein | St13 | *1.1±0.1 | 1.0±0.1 | 1.1±0.2 |
| Spliceosome RNA helicase Ddx39b | Ddx39b | *1.1±0.1 | 1.0±0.1 | 0.9±0.2 |
| Carboxy-terminal domain RNA polymerase II polypeptide A small phosphatase 1 | Ctdsp1 | *1.1±0.1 | 1.0±0.1 | 1.0±0.1 |
| Soluble calcium-activated nucleotidase 1 | Cant1 | *1.1±0.0 | 0.9±0.1 | 0.9±0.0 |
| Coagulation factor XIII B chain | F13b | *1.1±0.1 | 1.0±0.2 | 0.8±0.2 |
| Proteasome subunit beta type-1 | Psmb1 | *1.1±0.1 | 1.0±0.1 | 1.3±0.3 |
| Nck-associated protein 1 | Nckap1 | *1.1±0.1 | 1.0±0.0 | 0.8±0.1 |
| Platelet-activating factor acetylhydrolase IB subunit alpha | Pafah1b1 | *1.1±0.1 | 1.0±0.1 | 0.9±0.1 |
| SUMO-conjugating enzyme UBC9 | Ube2i | *1.1±0.1 | 1.0±0.1 | 0.8±0.1 |
| Proteasome subunit alpha type-3 | Psma3 | *1.1±0.1 | 1.0±0.1 | 1.2±0.3 |
| Dystrophin | Dmd | *1.1±0.1 | 1.0±0.2 | 1.0±0.2 |
| Threonine--tRNA ligase, mitochondrial | Tars2 | *1.1±0.0 | 0.9±0.2 | 1.2±0.5 |
| GTPase NRas | Nras | *1.1±0.0 | 1.1±0.1 | 0.9±0.2 |
| Acylamino-acid-releasing enzyme | Apeh | *1.1±0.1 | *1.3±0.1 | 1.1±0.3 |
| Basigin | Bsg | *1.1±0.0 | 1.0±0.1 | 0.8±0.1 |
| E3 ubiquitin-protein ligase HUWE1 | Huwe1 | *1.1±0.0 | 1.0±0.0 | 0.9±0.2 |
| Kinesin-1 heavy chain | Kif5b | *1.1±0.1 | 1.1±0.1 | 0.9±0.1 |
| TP53-regulated inhibitor of apoptosis 1 | Triap1 | *1.1±0.0 | 1.0±0.2 | 1.1±0.1 |
| CapZ-interacting protein | Rcsd1 | *1.1±0.1 | 0.9±0.1 | 0.9±0.3 |
| Tripartite motif-containing protein 14 | Trim14 | *1.1±0.0 | *1.2±0.2 | 0.9±0.2 |
| Filamin-B | Flnb | *1.1±0.1 | 0.9±0.1 | 0.8±0.1 |
| Conserved oligomeric Golgi complex subunit 8 | Cog8 | *1.1±0.0 | 0.9±0.1 | 1.2±0.3 |
| Trafficking protein particle complex subunit 11 | Trappc11 | *1.1±0.0 | 1.0±0.1 | 0.9±0.2 |
| Transmembrane 9 superfamily member 2 | Tm9sf2 | *1.1±0.1 | 0.9±0.0 | 1.1±0.2 |
| RuvB-like 2 | Ruvbl2 | *1.1±0.1 | 1.0±0.1 | 0.9±0.2 |

| | | | | |
|---|---|---|---|---|
| Probable ATP-dependent RNA helicase DDX5 | Ddx5 | *1.1±0.1 | 1.0±0.1 | 1.0±0.2 |
| Exportin-1 | Xpo1 | *1.1±0.0 | 1.0±0.0 | 0.8±0.2 |

These values represent average (± standard deviation) fold-change of abundance ratios for each altered (upregulated) protein compared to the high-fat control group (MS-NASH mice on a high-fat diet) with a 1.1-fold change threshold in response to Aquamin intervention and are significant with a p-value <0.05 (*). For each upregulated protein with Aquamin, corresponding values from the other two groups are shown for comparison. These liver samples (from 5 mice in each group) were individually assessed by TMT-based differential proteomic expression and data were merged to get averages. These data are also presented in Figure 4.
