## Supplemental Table 5 for "Liver Protein Expression in Nash Mice on a High-Fat Diet"

**Supplement Table 5. Top pathways associated with significantly upregulated proteins (shown in S Table 4) altered with Aquamin**

| Pathway name | Entities pValue | Mapped entities |
|---|---|---|
| RHO GTPases Activate WASPs and WAVEs | $1.01\times10^{-4}$ | Rac1;Wasf2;Cyfip1;Nckap1;Baiap2;Actb |
| Neutrophil degranulation | $3.99\times10^{-4}$ | Ahsg;Stom;Mvp;Gpi1;Pygb;Cd47;Cant1;Rab5c;Ostf1;Vat1;Cap1;Rac1;Pfkl;Hmgb1;Cyfip1; Lgals3;Huwe1;Cotl1;H2-Q10;Apeh;Anxa2;Nras;Psmb1 |
| VEGFA-VEGFR2 Pathway | $6.09\times10^{-4}$ | Rac1;Wasf2;Cyfip1;Nckap1;Baiap2;Pxn;Nras;Actb |
| Platelet degranulation | $8.98\times10^{-4}$ | Ahsg;Serpinf2;Actn1;Alb;Apoh;Anxa5;Apoa1;Actn4;Wdr1 |
| Signaling by VEGF | 0.001 | Rac1;Wasf2;Cyfip1;Nckap1;Baiap2;Pxn;Nras;Actb |
| Response to elevated platelet cytosolic Ca2+ | 0.001 | Ahsg;Serpinf2;Actn1;Alb;Apoh;Anxa5;Apoa1;Actn4;Wdr1 |
| Glycolysis | 0.002 | Pfkl;Gpi1;Eno1;Gck;Pfkp;Pklr |
| RHO GTPase Effectors | 0.002 | Wasf2;Tax1bp3;Nckap1;Kif5b;Baiap2;Tuba1c;Rac1;Cyfip1;Ppp2ca;Myl6;Pafah1b1;Pfn1;Ywhae; Actb |
| Dissolution of Fibrin Clot | 0.002 | Serpinf2;S100a10;Anxa2 |
| PTK6 Regulates RHO GTPases, RAS GTPase and MAP kinases | 0.003 | Rac1;Pxn;Nras |
| SUMOylation of DNA methylation proteins | 0.003 | Sumo1;Ube2i |
| mRNA Splicing - Major Pathway | 0.004 | Wdr33;Ddx5;Lsm3;Hnrnpd;Dhx15;Dnajc8;Fus;Hnrnpu;Hnrnpul1;Lsm6 |
| SUMO is transferred from E1 to E2 (UBE2I, UBC9) | 0.005 | Sumo1;Ube2i |
| mRNA Splicing | 0.005 | Wdr33;Ddx5;Lsm3;Hnrnpd;Dhx15;Dnajc8;Fus;Hnrnpu;Hnrnpul1;Lsm6 |
| Cell-extracellular matrix interactions | 0.006 | Actn1;Pxn;Actb |
| Innate Immune System | 0.006 | Ahsg;Stom;Psma3;Mvp;Nckap1;Baiap2;Cd47;Rab5c;Tbk1;Ostf1;Vat1;Cap1;Rac1;Uba3;Pfkl; Cyfip1;Huwe1;H2-Q10;Apeh;Anxa2;Psmb1;Actb;Wasf2;Gpi1;Fcgr1;Ripk1;Pygb;Cant1; Hmgb1;Lgals3;Ppp2ca;Cotl1;Nras |
| Glucose metabolism | 0.007 | Pfkl;Gpi1;Eno1;Gck;Pfkp;Pklr |
| SUMOylation of transcription cofactors | 0.008 | Sin3a;Sumo1;Ddx5;Ube2i |
| Nuclear Envelope (NE) Reassembly | 0.008 | Sumo1;Ppp2ca;Ube2i;Tuba1c;Chmp2b |
| Processing of Capped Intron-Containing Pre-mRNA | 0.010 | Wdr33;Ddx39b;Ddx5;Lsm3;Hnrnpd;Dhx15;Dnajc8;Fus;Hnrnpu;Hnrnpul1;Lsm6 |
| Regulation of cytoskeletal remodeling and cell spreading by IPP complex components | 0.012 | Actn1;Pxn |
| Regulation of HSF1-mediated heat shock response | 0.013 | Hspa4;St13;Bag1;Ccar2;Ywhae |

| | | |
|---|---|---|
| Platelet activation, signaling and aggregation | 0.014 | Ahsg;Rac1;Serpinf2;Actn1;Alb;Apoh;Anxa5;Apoa1;Actn4;Wdr1;Gnai2 |
| HDL remodeling | 0.014 | Alb;Apoa1 |
| Negative regulation of activity of TFAP2 (AP-2) family transcription factors | 0.014 | Sumo1;Ube2i |
| Processing and activation of SUMO | 0.018 | Sumo1;Ube2i |
| Hemostasis | 0.018 | Ahsg;Serpinf2;Actn1;S100a10;Alb;Kif5b;Cd47;Anxa5;Bsg;Tuba1c;Serpind1;Rac1;Apoh;F13b; Apoa1;Actn4;Ppp2ca;Anxa2;Nras;Wdr1;Gnai2 |
| SUMOylation of immune response proteins | 0.021 | Sumo1;Ube2i |
| RHO GTPases activate KTN1 | 0.021 | Rac1;Kif5b |
| RHO GTPases activate IQGAPs | 0.025 | Rac1;Actb |
| Glutathione synthesis and recycling | 0.025 | Ggct;Gclc |
| Deadenylation-dependent mRNA decay | 0.026 | Dcps;Lsm3;Cnot10;Lsm6 |
| Pentose phosphate pathway | 0.029 | Tkt;Pgd |
| Glutathione conjugation | 0.030 | Ggct;Gclc;Gstm3 |
| RAC2 GTPase cycle | 0.032 | Dsg2;Wasf2;Cyfip1;Nckap1;Arhgdia |
| mRNA decay by 5' to 3' exoribonuclease | 0.033 | Lsm3;Lsm6 |
| Cellular response to heat stress | 0.033 | Hspa4;St13;Bag1;Ccar2;Ywhae |
| SUMOylation of transcription factors | 0.037 | Sumo1;Ube2i |
| RAC3 GTPase cycle | 0.039 | Dsg2;Wasf2;Cyfip1;Nckap1;Baiap2 |
| Abasic sugar-phosphate removal via the single-nucleotide replacement pathway | 0.039 | Apex1 |
| Insulin effects increased synthesis of Xylulose-5-Phosphate | 0.039 | Tkt |
| Interleukin-37 signaling | 0.039 | Tbk1 |
| Signaling by Rho GTPases | 0.047 | Arhgef7;Wasf2;Stom;Actn1;Ddx39b;Tax1bp3;Nckap1;Kif5b;Baiap2;Arhgdia;Tuba1c;Dsg2;Rac1; Cyfip1;Ppp2ca;Myl6;Pafah1b1;Pfn1;Ywhae;Actb |

The pathways listed here are altered by the significantly upregulated proteins with the intervention “Aquamin” presented in Supplement Table 4. Reactome (v78) was used to generate the pathway analysis report for species *Mus musculus.* The significance (p-value) is calculated by the overrepresentation analysis (hypergeometric distribution).
