## Supplemental Table 6 for "Liver Protein Expression in Nash Mice on a High-Fat Diet"

**Supplement Table 6. Significantly altered (Downregulated) proteins with Aquamin in high-fat mice**

| Proteins | Genes | MS-NASH Aquamin | MS-NASH OCA | C57BL6 (Low-Fat) |
|---|---|---|---|---|
| Cytochrome P450 2C54 | Cyp2c54 | *0.48±0.14 | *0.61±0.19 | 2.72±0.49 |
| Major intrinsically disordered NOTCH2-binding receptor 1-like homolog | Minar2 | *0.49±0.12 | *0.45±0.18 | 1.06±0.23 |
| Alpha-1-antitrypsin 1-5 | Serpina1e | *0.52±0.27 | 0.70±0.35 | 37.05±11.87 |
| Protein CASC3 | Casc3 | *0.57±0.13 | *0.55±0.16 | *0.60±0.22 |
| Major urinary protein 2 | Mup2 | *0.59±0.23 | *0.44±0.19 | 10.40±5.30 |
| Cysteine dioxygenase type 1 | Cdo1 | *0.59±0.18 | *0.54±0.11 | 1.94±0.66 |
| Sodium-coupled neutral amino acid transporter 4 | Slc38a4 | *0.60±0.27 | *0.62±0.27 | 1.30±0.37 |
| Ferritin light chain 1 | Ftl1 | *0.62±0.23 | 0.74±0.27 | 1.18±0.80 |
| 40S ribosomal protein S30 | Fau | *0.63±0.33 | 0.71±0.26 | 0.80±0.42 |
| Sodium-coupled neutral amino acid transporter 3 | Slc38a3 | *0.63±0.13 | 0.71±0.18 | 2.60±1.10 |
| 2-oxo-4-hydroxy-4-carboxy-5-ureidoimidazoline decarboxylase | Urad | *0.64±0.20 | 0.94±0.20 | 3.19±2.07 |
| Cytochrome P450 2C50 | Cyp2c50 | *0.65±0.22 | 0.79±0.40 | 4.82±2.53 |
| 5'-nucleotidase | Nt5e | *0.66±0.28 | *0.67±0.19 | 1.33±0.30 |
| Ileal sodium/bile acid cotransporter | Slc10a2 | *0.67±0.24 | 0.77±0.22 | *0.54±0.27 |
| Ethanolamine-phosphate phospho-lyase | Etnppl | *0.69±0.26 | 1.55±0.67 | 1.37±0.38 |
| Cytochrome P450 2C29 | Cyp2c29 | *0.69±0.25 | *0.74±0.24 | 1.68±0.81 |
| Leucine-rich repeat-containing protein 9 | Lrrc9 | *0.71±0.17 | 0.98±0.23 | *0.05±0.01 |
| Histidine ammonia-lyase | Hal | *0.71±0.27 | 2.54±0.83 | 1.88±0.49 |
| Carboxylesterase 3B | Ces3b | *0.71±0.16 | 1.34±0.32 | 5.56±2.65 |
| Magnesium-dependent phosphatase 1 | Mdp1 | *0.72±0.21 | 0.83±0.18 | 0.96±0.13 |
| Hepatocyte nuclear factor 1-alpha | Hnf1a | *0.72±0.21 | 0.80±0.30 | 1.18±0.45 |
| Sodium/bile acid cotransporter | Slc10a1 | *0.72±0.21 | 1.21±0.37 | 1.88±1.04 |
| Aquaporin-9 | Aqp9 | *0.73±0.23 | *0.63±0.09 | 1.17±0.48 |
| Cytochrome P450 2C37 | Cyp2c37 | *0.73±0.19 | *0.71±0.19 | 2.55±1.14 |
| Dolichyl-phosphate beta-glucosyltransferase | Alg5 | *0.73±0.21 | *0.68±0.20 | *0.66±0.19 |
| Golgi SNAP receptor complex member 2 | Gosr2 | *0.73±0.21 | *0.75±0.16 | 0.82±0.29 |
| Cytochrome c oxidase subunit 7A-related protein, mitochondrial | Cox7a2l | *0.74±0.10 | *0.73±0.03 | 1.32±0.06 |
| Isovaleryl-CoA dehydrogenase, mitochondrial | Ivd | *0.74±0.10 | 0.89±0.12 | 2.57±0.87 |
| Cytochrome P450 7B1 | Cyp7b1 | *0.74±0.21 | *0.63±0.31 | 15.31±8.01 |
| Cancer-related nucleoside-triphosphatase homolog | Ntpcr | *0.74±0.19 | *0.81±0.18 | *0.61±0.16 |
| Cytochrome P450 4A12A | Cyp4a12a | *0.75±0.22 | 0.75±0.45 | 0.70±0.32 |
| Cytochrome P450 2C44 | Cyp2c23 | *0.76±0.12 | 1.09±0.27 | 2.50±0.46 |
| Thioredoxin domain-containing protein 15 | Txndc15 | *0.76±0.11 | *0.76±0.14 | 1.17±0.24 |

| | | | | |
|---|---|---|---|---|
| Hermansky-Pudlak syndrome 1 protein homolog | Hps1 | *0.76±0.17 | 0.78±0.43 | *0.56±0.15 |
| Vitamin K epoxide reductase complex subunit 1 | Vkorc1 | *0.76±0.06 | *0.86±0.08 | 1.05±0.14 |
| Homocysteine-responsive endoplasmic reticulum-resident ubiquitin-like domain member 2 protein | Herpud2 | *0.76±0.03 | 0.93±0.05 | 1.00±0.13 |
| Serine/threonine-protein phosphatase 6 regulatory ankyrin repeat subunit A | Ankrd28 | *0.77±0.04 | *0.80±0.09 | *0.84±0.01 |
| Multiple inositol polyphosphate phosphatase 1 | Minpp1 | *0.77±0.09 | 0.87±0.22 | *0.75±0.19 |
| Peroxisomal membrane protein 2 | Pxmp2 | *0.77±0.13 | 1.03±0.30 | 1.01±0.28 |
| Calcium uptake protein 1, mitochondrial | Micu1 | *0.77±0.02 | 0.95±0.02 | 1.06±0.21 |
| Sideroflexin-2 | Sfxn2 | *0.77±0.11 | 0.90±0.11 | 1.81±0.63 |
| Tetratricopeptide repeat protein 36 | Ttc36 | *0.78±0.09 | 1.02±0.10 | 1.19±0.16 |
| Delta(14)-sterol reductase TM7SF2 | Tm7sf2 | *0.78±0.20 | 0.85±0.18 | 2.28±0.76 |
| Hydroxyproline dehydrogenase | Prodh2 | *0.78±0.08 | 0.93±0.16 | 1.06±0.15 |
| Canalicular multispecific organic anion transporter 1 | Abcc2 | *0.78±0.19 | 1.04±0.24 | 1.03±0.26 |
| 28S ribosomal protein S25, mitochondrial | Mrps25 | *0.79±0.18 | *0.74±0.18 | 0.97±0.20 |
| Cytochrome P450 2D11 | Cyp2d11 | *0.79±0.14 | 0.88±0.31 | 1.17±0.42 |
| Transmembrane protein 19 | Tmem19 | *0.79±0.11 | *0.83±0.12 | 0.90±0.24 |
| Methionine-R-sulfoxide reductase B1 | Msrb1 | *0.79±0.12 | 0.94±0.13 | 1.20±0.10 |
| Bifunctional UDP-N-acetylglucosamine 2-epimerase/N-acetylmannosamine kinase | Gne | *0.79±0.14 | 1.00±0.20 | 1.38±0.30 |
| Serine protease inhibitor A3K | Serpina3k | *0.80±0.13 | 0.88±0.16 | 3.28±0.96 |
| Surfeit locus protein 4 | Surf4 | *0.80±0.17 | *0.70±0.15 | *0.67±0.24 |
| Lipoamide acyltransferase component of branched-chain alpha-keto acid dehydrogenase complex, mitochondrial | Dbt | *0.80±0.14 | 1.04±0.19 | 1.50±0.25 |
| 28S ribosomal protein S12, mitochondrial | Mrps12 | *0.80±0.09 | 0.89±0.18 | 1.12±0.11 |
| Receptor-type tyrosine-protein phosphatase delta | Ptprd | *0.81±0.08 | 1.19±0.13 | 1.67±0.47 |
| Tubulin gamma-1 chain | Tubg1 | *0.81±0.14 | 1.05±0.32 | *0.72±0.17 |
| NADH dehydrogenase [ubiquinone] iron-sulfur protein 5 | Ndufs5 | *0.81±0.15 | 0.88±0.17 | 0.91±0.13 |
| Dolichol-phosphate mannosyltransferase subunit 1 | Dpm1 | *0.81±0.02 | *0.80±0.07 | *0.81±0.17 |
| Apolipoprotein A-V | Apoa5 | *0.81±0.17 | 1.08±0.23 | 0.83±0.20 |
| Actin-related protein 2/3 complex subunit 1A | Arpc1a | *0.82±0.12 | 1.13±0.08 | 1.32±0.13 |
| Kinesin-like protein KIF16B | Kif16b | *0.82±0.04 | 1.09±0.49 | 1.08±0.50 |
| Tensin-2 | Tns2 | *0.82±0.11 | 0.88±0.12 | 0.95±0.33 |
| Ferrochelatase, mitochondrial | Fech | *0.82±0.15 | 0.95±0.13 | 1.22±0.19 |
| Mitochondrial 2-oxodicarboxylate carrier | Slc25a21 | *0.82±0.06 | 0.95±0.12 | 0.81±0.16 |
| ATP-binding cassette sub-family B member 10, mitochondrial | Abcb10 | *0.82±0.08 | 1.02±0.13 | 1.28±0.13 |
| Probable N-acetyltransferase CML1 | Cml1 | *0.82±0.06 | 1.28±0.28 | 1.19±0.35 |
| Short/branched chain specific acyl-CoA dehydrogenase, mitochondrial | Acadsb | *0.82±0.06 | 1.02±0.15 | 1.70±0.44 |

| | | | | |
|---|---|---|---|---|
| Sulfite oxidase, mitochondrial | Suox | *0.82±0.10 | 1.07±0.20 | 1.26±0.29 |
| NADPH-dependent 3-keto-steroid reductase Hsd3b5 | Hsd3b5 | *0.83±0.12 | 0.78±0.26 | 19.97±21.84 |
| Serine--pyruvate aminotransferase, mitochondrial | Agxt | *0.83±0.15 | 1.08±0.11 | 2.55±1.07 |
| Mannose-binding protein C | Mbl2 | *0.83±0.12 | 0.95±0.07 | 1.24±0.19 |
| Hypoxia up-regulated protein 1 | Hyou1 | *0.83±0.14 | *0.81±0.10 | 1.40±0.27 |
| Inhibin beta C chain | Inhbc | *0.83±0.08 | *0.80±0.10 | *0.79±0.13 |
| Prolactin regulatory element-binding protein | Preb | *0.83±0.16 | *0.79±0.18 | 0.87±0.23 |
| Calcium load-activated calcium channel | Tmco1 | *0.83±0.16 | *0.80±0.18 | *0.69±0.23 |
| ER membrane protein complex subunit 6 | Emc6 | *0.84±0.04 | 0.84±0.28 | *0.71±0.06 |
| Kinesin-like protein KIF13A | Kif13a | *0.84±0.15 | 1.23±0.53 | 0.98±0.29 |
| Growth factor receptor-bound protein 7 | Grb7 | *0.84±0.12 | *0.91±0.07 | 0.98±0.26 |
| Probable D-lactate dehydrogenase, mitochondrial | Ldhd | *0.84±0.11 | *0.75±0.09 | 1.38±0.47 |
| Protein disulfide isomerase Creld1 | Creld1 | *0.84±0.05 | *0.81±0.05 | 1.50±0.43 |
| Transmembrane protein 106B | Tmem106b | *0.84±0.08 | 0.80±0.14 | *0.67±0.08 |
| Cytochrome b5 type B | Cyb5b | *0.84±0.08 | *0.71±0.17 | 1.36±0.63 |
| Sulfotransferase family cytosolic 1B member 1 | Sult1b1 | *0.85±0.07 | 1.32±0.18 | 0.90±0.22 |
| Atlastin-2 | Atl2 | *0.85±0.13 | *0.81±0.08 | *0.79±0.10 |
| ADP-ribosylation factor-like protein 6-interacting protein 1 | Arl6ip1 | *0.85±0.09 | *0.70±0.09 | 0.86±0.15 |
| Ethanolamine kinase 2 | Etnk2 | *0.85±0.06 | 1.08±0.19 | 1.38±0.39 |
| Long-chain fatty acid transport protein 4 | Slc27a4 | *0.85±0.11 | *0.63±0.09 | 0.76±0.31 |
| Monoacylglycerol lipase ABHD6 | Abhd6 | *0.86±0.05 | *0.73±0.10 | *0.76±0.14 |
| 39S ribosomal protein L20, mitochondrial | Mrpl20 | *0.86±0.09 | 0.96±0.27 | 1.53±0.70 |
| Methylcrotonoyl-CoA carboxylase beta chain, mitochondrial | Mccc2 | *0.86±0.07 | 1.00±0.06 | 1.45±0.15 |
| ATP synthase protein 8 | ATP8 | *0.86±0.12 | 1.01±0.06 | 0.95±0.27 |
| Probable ATP-dependent RNA helicase DHX58 | Dhx58 | *0.86±0.06 | 0.84±0.23 | 0.67±0.24 |
| Perilipin-2 | Plin2 | *0.86±0.11 | *0.56±0.10 | *0.18±0.01 |
| Methylcrotonoyl-CoA carboxylase subunit alpha, mitochondrial | Mccc1 | *0.86±0.09 | 1.02±0.13 | 1.30±0.28 |
| Serine beta-lactamase-like protein LACTB, mitochondrial | Lactb | *0.87±0.07 | *0.88±0.06 | 1.12±0.13 |
| Emopamil-binding protein-like | Ebpl | *0.87±0.12 | *0.72±0.11 | *0.38±0.21 |
| Sideroflexin-5 | Sfxn5 | *0.87±0.07 | 0.96±0.10 | *0.71±0.14 |
| Secretory carrier-associated membrane protein 3 | Scamp3 | *0.87±0.11 | 0.94±0.16 | 1.13±0.50 |
| Adenosylhomocysteinase | Ahcy | *0.87±0.12 | 1.21±0.18 | 1.33±0.35 |
| Methylmalonic aciduria type A homolog, mitochondrial | Mmaa | *0.87±0.12 | 1.00±0.22 | 0.97±0.26 |
| Histidine triad nucleotide-binding protein 3 | Hint3 | *0.87±0.05 | *0.87±0.13 | 1.02±0.06 |
| Transmembrane protein 143 | Tmem143 | *0.87±0.08 | 0.91±0.11 | *0.72±0.18 |
| Protein jagunal homolog 1 | Jagn1 | *0.87±0.08 | *0.86±0.06 | 0.91±0.20 |
| ATP synthase subunit epsilon, mitochondrial | Atp5e | *0.87±0.07 | 0.90±0.13 | 0.89±0.11 |

| | | | | |
|---|---|---|---|---|
| Protein phosphatase methylesterase 1 | Ppme1 | *0.87±0.09 | 0.97±0.12 | 0.60±0.19 |
| Transmembrane protein 205 | Tmem205 | *0.87±0.05 | *0.83±0.14 | 1.52±0.33 |
| Methionine aminopeptidase 1 | Metap1 | *0.88±0.08 | 1.01±0.19 | *0.90±0.05 |
| Acyl-coenzyme A synthetase ACSM5, mitochondrial | Acsm5 | *0.88±0.09 | 1.01±0.12 | 1.51±0.29 |
| Putative peptidyl-tRNA hydrolase PTRHD1 | Ptrhd1 | *0.88±0.07 | 0.95±0.20 | *0.63±0.28 |
| 60S ribosomal protein L11 | Rpl11 | *0.88±0.10 | *0.86±0.12 | 0.99±0.30 |
| Dolichol-phosphate mannosyltransferase subunit 3 | Dpm3 | *0.88±0.08 | *0.85±0.09 | 0.91±0.12 |
| Syndecan-1 | Sdc1 | *0.88±0.11 | 0.92±0.17 | *0.69±0.16 |
| MICOS complex subunit Mic10 | Minos1 | *0.89±0.08 | *0.83±0.14 | 1.13±0.33 |
| Equilibrative nucleoside transporter 1 | Slc29a1 | *0.89±0.06 | 1.05±0.15 | 1.59±0.34 |
| Growth hormone receptor | Ghr | *0.89±0.07 | 1.01±0.16 | 0.89±0.06 |
| 28S ribosomal protein S6, mitochondrial | Mrps6 | *0.89±0.06 | 1.00±0.05 | 1.13±0.22 |
| Glutaredoxin-related protein 5, mitochondrial | Glrx5 | *0.89±0.08 | 0.98±0.13 | 0.96±0.15 |
| Peptidyl-prolyl cis-trans isomerase F, mitochondrial | Ppif | *0.89±0.05 | 1.02±0.10 | 1.17±0.21 |
| Complex I intermediate-associated protein 30, mitochondrial | Ndufaf1 | *0.90±0.05 | 1.08±0.14 | 1.39±0.12 |
| Protein disulfide-isomerase A4 | Pdia4 | *0.90±0.10 | *0.78±0.03 | 1.33±0.12 |
| Succinate--CoA ligase [ADP-forming] subunit beta, mitochondrial | Sucla2 | *0.90±0.05 | 1.01±0.06 | 0.95±0.24 |
| FERM, ARHGEF and pleckstrin domain-containing protein 2 | Farp2 | *0.90±0.04 | 0.95±0.11 | 1.04±0.21 |
| Elongation factor G, mitochondrial | Gfm1 | *0.90±0.09 | 1.01±0.10 | 0.98±0.12 |
| Adenylosuccinate synthetase isozyme 1 | Adssl1 | *0.90±0.07 | 1.41±0.17 | *0.79±0.06 |
| Mitochondrial import inner membrane translocase subunit Tim8 B | Timm8b | *0.90±0.06 | *0.77±0.10 | 1.09±0.14 |
| Lipid droplet-associated hydrolase | Ldah | *0.91±0.06 | *0.75±0.08 | *0.75±0.03 |
| Inositol 1,4,5-trisphosphate receptor type 2 | Itpr2 | *0.91±0.08 | *0.87±0.06 | 0.88±0.25 |
| DnaJ homolog subfamily A member 1 | Dnaja1 | *0.91±0.07 | *0.91±0.06 | 1.04±0.10 |
| Peptidyl-tRNA hydrolase 2, mitochondrial | Ptrh2 | *0.91±0.05 | *0.89±0.07 | 0.98±0.18 |
| Serine/threonine-protein kinase TAO1 | Taok1 | *0.91±0.06 | *0.90±0.07 | *0.80±0.08 |

These values represent average (± standard deviation) fold-change of abundance ratios for each altered (downregulated) protein compared to the high-fat control group (MS-NASH mice on a high-fat diet) with a 1.1-fold change threshold in response to Aquamin intervention and are significant with a p-value <0.05 (*). For each upregulated protein with Aquamin, corresponding values from the other two groups are shown for comparison. These liver samples (from 5 mice in each group) were individually assessed by TMT-based differential proteomic expression and data were merged to get averages. These data are also presented in Figure 4.
