## Supplemental Table 7 for "Liver Protein Expression in Nash Mice on a High-Fat Diet"

**Supplement Table 7. Top pathways associated with significantly downregulated proteins (shown in S Table 5) altered with Aquamin**

| Pathway name | Entities pValue | Mapped entities |
|---|---|---|
| Branched-chain amino acid catabolism | $1.20 \times 10^{-5}$ | Acadsb;Mccc1;Mccc2;Dbt;Ivd |
| Metabolism of amino acids and derivatives | $2.88 \times 10^{-4}$ | Hal;Suox;Acadsb;Agxt;Mccc1;Slc25a21;Mccc2;Prodh2;Dbt;Cdo1;Ivd;Ahcy |
| Synthesis of dolichyl-phosphate mannose | $8.95 \times 10^{-4}$ | Dpm3;Dpm1 |
| Metabolism | 0.003 | Cyp4a12a;Ces3b;Slc25a21;Slc10a1;Abcc2;Ndufs5;Acsm5;Minpp1;Acadsb;Suox;Agxt;Mccc1;Etnk2;Cdo1;Atp5e;Fech;Slc10a2;ATP8;Nt5e;Mccc2;Adssl1;Prodh2;Dbt;Mmaa;Cyb5b;Ivd;Itpr2;Ndufaf1;Cox7a2l;Cyp7b1;Ahcy;Hal;Etnppl;Sucla2;Tm7sf2;Sdc1;Glrx5;Vkorc1;Sult1b1 |
| Sulfur amino acid metabolism | 0.005 | Suox;Cdo1;Ahcy |
| Mitochondrial translation elongation | 0.008 | Mrps25;Mrps6;Mrpl20;Mrps12;Gfm1 |
| Glyoxylate metabolism and glycine degradation | 0.010 | Agxt;Prodh2;Dbt |
| Mitochondrial translation | 0.010 | Mrps25;Mrps6;Mrpl20;Mrps12;Gfm1 |
| Biotin transport and metabolism | 0.011 | Mccc1;Mccc2 |
| Synthesis of substrates in N-glycan biosythesis | 0.012 | Alg5;Dpm3;Dpm1;Gne |
| Synthesis of PE | 0.013 | Etnppl;Etnk2 |
| Degradation of cysteine and homocysteine | 0.013 | Suox;Cdo1 |
| Synthesis of IPs in the ER lumen | 0.014 | Minpp1 |
| Biosynthesis of the N-glycan precursor (dolichol lipid-linked oligosaccharide, LLO) and transfer to a nascent protein | 0.021 | Alg5;Dpm3;Dpm1;Gne |
| Bile acid and bile salt metabolism | 0.023 | Slc10a2;Slc10a1;Cyp7b1 |
| Recycling of bile acids and salts | 0.025 | Slc10a2;Slc10a1 |
| Formation of ATP by chemiosmotic coupling | 0.028 | ATP8;Atp5e |
| Cristae formation | 0.028 | ATP8;Atp5e |
| Respiratory electron transport, ATP synthesis by chemiosmotic coupling, and heat production by uncoupling proteins | 0.028 | ATP8;Atp5e;Ndufaf1;Ndufs5;Cox7a2l |

| | | |
|---|---|---|
| Transport of glycerol from adipocytes to the liver by Aquaporins | 0.028 | Aqp9 |
| Synthesis of dolichyl-phosphate-glucose | 0.028 | Alg5 |
| Biological oxidations | 0.033 | Cyp4a12a;Ces3b;Cyb5b;Cyp7b1;Acsm5;Ahcy;Sult1b1 |
| Metabolism of vitamins and cofactors | 0.035 | Nt5e;Mccc1;Mccc2;Sdc1;Mmaa;Vkorc1 |
| The citric acid (TCA) cycle and respiratory electron transport | 0.037 | ATP8;Sucla2;Atp5e;Ndufs5;Ndufaf1;Cox7a2l |
| Mitochondrial translation termination | 0.039 | Mrps25;Mrps6;Mrpl20;Mrps12 |
| Metabolism of vitamin K | 0.042 | Vkorc1 |
| Proline catabolism | 0.042 | Prodh2 |

The pathways listed here are altered by the significantly downregulated proteins with the intervention “Aquamin” presented in Supplement Table 5. Reactome (v78) was used to generate the pathway analysis report for species *Mus musculus.* The significance (p-value) is calculated by the overrepresentation analysis (hypergeometric distribution).
