## Supplemental Table 10 for "Liver Protein Expression in Nash Mice on a High-Fat Diet"

**Supplement Table 10. Upregulated Proteins by an unbiased proteomic screening of C57BL6 mice on low-fat diet**

| | | C57BL6 | MS-NASH | |
|---|---|---|---|---|
| **Proteins** | **Genes** | **Control** | **OCA** | **Aquamin** |
| Alpha-1-antitrypsin 1-5 | Serpina1e | 37.05±11.87* | 0.70±0.35 | 0.52±0.27 |
| NADPH-dependent 3-keto-steroid reductase Hsd3b5 | Hsd3b5 | 19.97±21.84 | 0.78±0.26 | 0.83±0.12 |
| Solute carrier organic anion transporter family member 1A1 | Slco1a1 | 18.33±10.59* | 1.03±0.73 | 1.05±0.66 |
| Cytochrome P450 7B1 | Cyp7b1 | 15.31±8.01* | 0.63±0.31 | 0.74±0.21 |
| Glycine cleavage system H protein, mitochondrial | Gcsh | 11.09±21.41 | 9.05±17.88 | 7.63±14.93 |
| Major urinary protein 20 | Mup20 | 10.78±14.91 | 1.02±0.37 | 0.90±0.50 |
| Major urinary protein 2 | Mup2 | 10.40±5.30* | 0.44±0.19 | 0.59±0.23 |
| Fucose mutarotase | Fuom | 8.72±17.40 | 10.03±19.94 | 10.28±20.61 |
| UDP-glucuronosyltransferase 3A2 | Ugt3a2 | 8.39±3.65* | 0.85±0.07 | 0.96±0.13 |
| Nuclear transport factor 2 | Nutf2 | 8.16±16.12 | 10.29±20.76 | 12.12±24.59 |
| Ras-related protein R-Ras | Rras | 6.32±11.74 | 4.75±8.44 | 5.46±9.57 |
| Glutathione S-transferase P 1 | Gstp1 | 6.24±2.43* | 0.92±0.14 | 1.14±0.15 |
| Epidermal growth factor receptor | Egfr | 6.21±1.51* | 0.79±0.19* | 0.87±0.18 |
| Alpha-1-acid glycoprotein 1 | Orm1 | 6.12±9.25 | 2.93±5.05 | 3.98±6.98 |
| SH3 domain-containing protein 21[#] | Sh3d21 | 6.05±10.62 | 5.43±10.12 | 8.30±16.34 |
| Transmembrane protein 14C | Tmem14c | 5.97±10.43 | 4.82±8.80 | 5.56±10.29 |
| Keratin, type II cytoskeletal 2 oral | Krt76 | 5.89±7.01 | 1.71±2.42 | 8.07±14.08 |
| Keratin, type I cytoskeletal 14 | Krt14 | 5.88±7.13 | 0.70±0.43 | 5.58±9.45 |
| 1,5-anhydro-D-fructose reductase | Akr1e2 | 5.75±1.76* | 1.65±1.00 | 1.24±0.50 |
| Ras-related protein Rap-1A | Rap1a | 5.66±10.21 | 5.83±10.73 | 5.20±9.18 |
| Selenium-binding protein 2 | Selenbp2 | 5.63±0.99* | 1.24±0.31 | 1.07±0.23 |
| Carboxylesterase 3B | Ces3b | 5.56±2.65* | 1.34±0.32* | 0.71±0.16 |
| Isopentenyl-diphosphate Delta-isomerase 1 | Idi1 | 5.50±4.41 | 1.62±0.95 | 0.98±0.35 |
| Keratin, type II cytoskeletal 79 | Krt79 | 5.40±4.07 | 5.73±7.97 | 26.34±43.11 |
| Tubulin beta-5 chain | Tubb5 | 4.90±9.25 | 5.06±9.36 | 5.81±10.51 |
| 60S acidic ribosomal protein P1 | Rplp1 | 4.84±8.27 | 3.61±5.97 | 3.53±5.59 |
| Cytochrome P450 2C50 | Cyp2c50 | 4.82±2.53* | 0.79±0.40 | 0.65±0.22 |
| Methylsterol monooxygenase 1 | Msmo1 | 4.54±1.38* | 1.38±0.45 | 1.15±0.54 |
| Isoamyl acetate-hydrolyzing esterase 1 homolog | Iah1 | 4.50±1.12* | 1.00±0.24 | 0.98±0.27 |
| ATP-binding cassette sub-family D member 4 | Abcd4 | 4.49±4.10 | 2.12±1.99 | 1.50±1.19 |
| Adenylate kinase 4, mitochondrial | Ak4 | 4.42±1.10* | 1.75±0.21* | 1.00±0.13 |
| 60S ribosomal protein L36 | Rpl36 | 4.35±8.31 | 4.95±8.52 | 4.32±7.31 |
| Inositol-3-phosphate synthase 1 | Isyna1 | 4.32±5.70 | 0.79±0.23 | 0.90±0.24 |

| | | | | |
|---|---|---|---|---|
| Lanosterol 14-alpha demethylase | Cyp51a1 | 4.21±2.01* | 1.38±0.72 | 0.99±0.30 |
| IgG receptor FcRn large subunit p51 | Fcgrt | 4.17±6.07 | 5.74±10.39 | 4.12±6.83 |
| Keratin, type I cytoskeletal 10 | Krt10 | 4.10±6.43 | 1.99±3.15 | 4.86±8.83 |
| Ig alpha chain C region | n/a | 4.06±1.96* | 0.96±0.40 | 1.03±0.48 |
| Pyrethroid hydrolase Ces2a | Ces2a | 4.00±1.62* | 1.49±0.51 | 0.96±0.27 |
| Keratin, type II cytoskeletal 1 | Krt1 | 3.96±3.36 | 1.80±2.51 | 10.38±20.44 |
| 4-hydroxy-2-oxoglutarate aldolase, mitochondrial | Hoga1 | 3.81±2.81 | 2.05±1.87 | 1.27±0.85 |
| UDP-glucuronosyltransferase 1-9 | Ugt1a9 | 3.78±4.76 | 1.73±1.63 | 1.68±1.68 |
| Glutaredoxin-1 | Glrx | 3.76±5.34 | 2.85±3.94 | 2.58±3.23 |
| Tubulin beta-4A chain | Tubb4a | 3.76±6.90 | 4.73±8.30 | 4.85±8.51 |
| Keratin, type I cytoskeletal 42 | Krt42 | 3.74±3.98 | 1.26±1.63 | 7.91±13.57 |
| Aldehyde oxidase 2 | Aox2 | 3.70±2.76 | 1.86±0.89 | 1.22±0.36 |
| Serum amyloid A-1 protein | Saa1 | 3.65±4.61 | 0.45±0.19 | 0.73±0.40 |
| Carbamoyl-phosphate synthase [ammonia], mitochondrial | Cps1 | 3.62±2.04* | 1.39±0.35* | 0.90±0.33 |
| Bile salt sulfotransferase 1 | Sult2a1 | 3.60±2.35* | 0.98±0.50 | 0.79±0.23 |
| Glutaminase liver isoform, mitochondrial | Gls2 | 3.56±1.50* | 1.53±0.29* | 0.83±0.31 |
| Keratin, type II cytoskeletal 5 | Krt5 | 3.56±4.11 | 1.60±2.45 | 11.14±22.14 |
| Frataxin, mitochondrial | Fxn | 3.54±3.31 | 1.80±1.67 | 1.97±2.15 |
| Keratin, type I cytoskeletal 17 | Krt17 | 3.50±2.69 | 1.23±1.59 | 10.62±21.47 |
| Peroxisomal coenzyme A diphosphatase NUDT7 | Nudt7 | 3.48±2.23* | 1.14±0.80 | 1.12±0.72 |
| Splicing factor 3B subunit 4 | Sf3b4 | 3.46±4.27 | 3.57±5.19 | 3.03±4.20 |
| Cytochrome b-c1 complex subunit 10 | Uqcr11 | 3.40±4.36 | 2.45±3.24 | 2.47±3.24 |
| Carboxypeptidase B2 | Cpb2 | 3.39±4.66 | 2.29±2.74 | 2.39±2.63 |
| Complement component C8 beta chain | C8b | 3.39±1.08* | 1.07±0.18 | 0.87±0.30 |
| Beta-2-microglobulin | B2m | 3.35±3.11 | 0.95±0.24 | 1.31±0.42 |
| Pigment epithelium-derived factor | Serpinf1 | 3.34±5.88 | 5.58±10.41 | 6.03±10.85 |
| Lysosomal acid glucosylceramidase | Gba | 3.31±0.46* | 1.54±0.52 | 1.33±0.31 |
| Serine protease inhibitor A3K | Serpina3k | 3.28±0.96* | 0.88±0.16 | 0.80±0.13 |
| Coatomer subunit epsilon | Cope | 3.27±4.74 | 2.78±4.10 | 3.50±5.59 |
| Proteasome subunit beta type-3 | Psmb3 | 3.27±4.30 | 3.50±5.41 | 3.56±5.56 |
| Endophilin-B1 | Sh3glb1 | 3.27±5.18 | 3.47±5.80 | 3.93±6.42 |
| U8 snoRNA-decapping enzyme | Nudt16 | 3.26±4.15 | 3.02±4.24 | 4.03±5.49 |
| Glycine N-acyltransferase-like protein Keg1 | Keg1 | 3.25±0.94* | 1.06±0.23 | 0.88±0.21 |
| Mevalonate kinase | Mvk | 3.22±1.45* | 1.24±0.47 | 1.09±0.35 |
| 2-oxo-4-hydroxy-4-carboxy-5-ureidoimidazoline decarboxylase | Urad | 3.19±2.07* | 0.94±0.20 | 0.64±0.20 |
| Beta-ureidopropionase | Upb1 | 3.19±0.80* | 1.34±0.51 | 1.05±0.25 |
| Biotinidase | Btd | 3.10±1.91* | 1.31±0.72 | 1.35±0.99 |

| | | | | |
|---|---|---|---|---|
| Protein PAT1 homolog 1[#] | Patl1 | 3.09±2.72 | 2.74±2.40 | 4.99±4.23 |
| Signal recognition particle 19 kDa protein | Srp19 | 3.08±4.09 | 2.29±3.07 | 2.46±3.39 |
| Aldehyde oxidase 3 | Aox3 | 3.00±1.58* | 1.59±0.49* | 1.16±0.25 |
| Epididymis-specific alpha-mannosidase | Man2b2 | 2.96±1.97 | 1.64±1.54 | 1.82±1.75 |
| Cysteine sulfinic acid decarboxylase | Csad | 2.91±1.74* | 0.50±0.24 | 1.88±1.14 |
| 2-hydroxyacyl-CoA lyase 2 | Ilvbl | 2.91±4.43 | 2.82±4.22 | 2.99±4.38 |
| Pancreatic alpha-amylase | Amy2 | 2.89±3.48 | 0.67±0.54 | 0.85±0.69 |
| Small nuclear ribonucleoprotein E | Snrpe | 2.86±3.61 | 2.22±2.99 | 2.90±4.12 |
| Galectin-related protein | Lgalsl | 2.86±4.63 | 3.52±5.77 | 3.26±5.14 |
| Fatty acid-binding protein 5 | Fabp5 | 2.85±1.63* | 0.91±0.48 | 1.11±0.33 |
| Galactose-1-phosphate uridylyltransferase | Galt | 2.84±3.21 | 3.02±4.19 | 2.14±2.67 |
| Aldose reductase-related protein 2 | Akr1b8 | 2.84±3.75 | 2.25±2.73 | 2.75±3.39 |
| Cathepsin F | Ctsf | 2.84±1.10* | 1.10±0.40 | 1.11±0.44 |
| Phosphoenolpyruvate carboxykinase, cytosolic [GTP] | Pck1 | 2.83±1.39* | 1.41±0.33* | 0.74±0.29 |
| Kynurenine/alpha-aminoadipate aminotransferase, mitochondrial | Aadat | 2.82±0.74* | 1.28±0.39 | 0.91±0.29 |
| Acyl-coenzyme A synthetase ACSM1, mitochondrial | Acsm1 | 2.82±0.86* | 1.16±0.32 | 1.02±0.31 |
| Kynurenine--oxoglutarate transaminase 1 | Kyat1 | 2.76±0.86* | 1.00±0.21 | 0.99±0.20 |
| Keratin, type II cytoskeletal 1b | Krt77 | 2.74±2.78 | 0.67±0.42 | 0.82±0.66 |
| Farnesyl pyrophosphate synthase | Fdps | 2.73±1.05* | 1.41±0.60 | 1.07±0.43 |
| Cytochrome P450 2C54 | Cyp2c54 | 2.72±0.49* | 0.61±0.19 | 0.48±0.14 |
| Arylacetamide deacetylase | Aadac | 2.70±2.02 | 1.45±1.19 | 1.34±1.01 |
| Putative RNA-binding protein Luc7-like 1 | Luc7l | 2.67±3.04 | 2.56±3.40 | 3.20±4.78 |
| Cytochrome b | Mt-Cyb | 2.66±2.80 | 1.83±1.81 | 1.48±1.03 |
| COMM domain-containing protein 8 | Commd8 | 2.63±2.14 | 2.16±1.99 | 2.26±2.73 |
| Cystathionine gamma-lyase | Cth | 2.63±0.81* | 1.50±0.32* | 0.92±0.22 |
| Leukemia inhibitory factor receptor | Lifr | 2.63±1.40 | 1.03±0.23 | 1.13±0.15 |
| Threonine synthase-like 2 | Thnsl2 | 2.63±0.21* | 1.08±0.46 | 1.13±0.57 |
| Carboxylesterase 3A | Ces3a | 2.61±1.04* | 0.65±0.14 | 1.01±0.33 |
| AP-2 complex subunit sigma | Ap2s1 | 2.61±3.59 | 2.05±2.57 | 2.45±3.06 |
| Sodium-coupled neutral amino acid transporter 3 | Slc38a3 | 2.60±1.10* | 0.71±0.18 | 0.63±0.13 |
| Phosphatidylethanolamine N-methyltransferase | Pemt | 2.60±3.30 | 2.02±2.17 | 1.81±1.56 |
| Alpha-aminoadipic semialdehyde synthase, mitochondrial | Aass | 2.60±2.27 | 1.33±0.59 | 0.87±0.29 |
| Ornithine carbamoyltransferase, mitochondrial | Otc | 2.59±0.85* | 1.47±0.52 | 1.07±0.35 |
| Cytochrome P450 2C70 | Cyp2c70 | 2.58±2.08 | 1.69±0.78 | 1.13±0.59 |
| Isovaleryl-CoA dehydrogenase, mitochondrial | Ivd | 2.57±0.87* | 0.89±0.12 | 0.74±0.10 |
| Complement component C8 gamma chain | C8g | 2.57±2.50 | 1.29±1.05 | 1.34±1.25 |
| Cation channel sperm-associated protein subunit beta[#] | Catsperb | 2.55±1.47 | 2.00±0.90 | 1.42±0.54 |

| | | | | |
|---|---|---|---|---|
| Serine--pyruvate aminotransferase, mitochondrial | Agxt | 2.55±1.07* | 1.08±0.11 | 0.83±0.15 |
| Cytochrome P450 2C37 | Cyp2c37 | 2.55±1.14* | 0.71±0.19 | 0.73±0.19 |
| NADH-ubiquinone oxidoreductase chain 5 | Mtnd5 | 2.55±3.40 | 2.44±3.18 | 2.42±3.25 |
| Apolipoprotein M# | Apom | 2.55±0.25* | 0.75±0.12 | 0.95±0.21 |
| Retinoid-inducible serine carboxypeptidase | Scpep1 | 2.53±0.55* | 0.80±0.12 | 0.92±0.21 |
| Aldo-keto reductase family 1 member C18 | Akr1c18 | 2.52±2.51 | 2.31±2.65 | 2.74±3.86 |
| Zinc finger and BTB domain-containing protein 20 | Zbtb20 | 2.52±2.14 | 1.82±1.49 | 1.33±0.71 |
| Nucleoside diphosphate kinase B | Nme2 | 2.51±1.07* | 1.03±0.20 | 0.98±0.19 |
| Ig gamma-1 chain C region, membrane-bound form | Ighg1 | 2.50±2.04 | 1.28±0.47 | 1.43±0.90 |
| Phosphopantothenoylcysteine decarboxylase | Ppcdc | 2.50±2.07 | 1.37±0.38 | 1.30±0.07* |
| Cytochrome P450 2C44 | Cyp2c23 | 2.50±0.46* | 1.09±0.27 | 0.76±0.12 |
| Cytochrome P450 1A2 | Cyp1a2 | 2.50±0.63* | 0.78±0.08 | 0.95±0.18 |
| Tyrosine aminotransferase | Tat | 2.46±0.62* | 0.87±0.24 | 0.72±0.27 |
| 4-aminobutyrate aminotransferase, mitochondrial | Abat | 2.46±1.39* | 1.38±0.75 | 0.93±0.43 |
| Ancient ubiquitous protein 1 | Aup1 | 2.45±3.38 | 2.04±2.78 | 2.00±2.73 |
| Protein CREG1 | Creg1 | 2.45±0.80* | 0.96±0.39 | 1.54±0.79 |
| Ergosterol biosynthetic protein 28 homolog | Erg28 | 2.44±1.88 | 1.61±1.46 | 1.42±1.19 |
| Peptidyl-prolyl cis-trans isomerase NIMA-interacting 1 | Pin1 | 2.43±2.54 | 2.51±2.90 | 2.52±2.87 |
| Lanosterol synthase | Lss | 2.41±1.16* | 1.40±0.98 | 1.16±0.38 |
| Sterol-4-alpha-carboxylate 3-dehydrogenase, decarboxylating | Nsdhl | 2.40±1.33* | 1.33±0.85 | 1.06±0.29 |
| Glycine N-methyltransferase | Gnmt | 2.40±1.09* | 1.31±0.45 | 0.85±0.37 |
| Tetratricopeptide repeat protein 39C | Ttc39c | 2.40±0.77* | 0.86±0.15 | 0.88±0.13 |
| Arylsulfatase B | Arsb | 2.39±0.65* | 0.71±0.05 | 1.03±0.08 |
| Carnitine O-palmitoyltransferase 2, mitochondrial | Cpt2 | 2.36±0.54* | 1.11±0.25 | 1.12±0.30 |
| CD81 antigen | Cd81 | 2.36±2.50 | 1.67±2.05 | 2.10±2.42 |
| Corrinoid adenosyltransferase | Mmab | 2.35±0.52* | 1.15±0.23 | 0.86±0.16 |
| STIP1 homology and U box-containing protein 1 | Stub1 | 2.33±2.80 | 2.22±2.68 | 2.07±2.40 |
| Neuroplastin | Nptn | 2.33±2.72 | 2.11±2.67 | 2.44±3.07 |
| Cytochrome P450 4V2 | Cyp4v3 | 2.32±0.97* | 1.05±0.39 | 1.04±0.39 |
| Protein FAM210B, mitochondrial | Fam210b | 2.32±1.67 | 0.92±0.21 | 0.96±0.17 |
| Condensin-2 complex subunit D3# | Ncapd3 | 2.31±1.46 | 1.31±0.95 | 1.69±0.86 |
| Cytochrome c oxidase subunit 6A1, mitochondrial | Cox6a1 | 2.30±3.14 | 1.96±2.32 | 1.93±2.32 |
| Nuclear cap-binding protein subunit 1 | Ncbp1 | 2.29±2.99 | 2.41±3.23 | 2.54±3.24 |
| Nascent polypeptide-associated complex subunit alpha, muscle-specific form | Naca | 2.29±2.60 | 2.15±2.63 | 2.13±2.58 |
| Mitogen-activated protein kinase kinase kinase 11# | Map3k11 | 2.28±0.43* | 1.17±0.31 | 1.30±0.35 |
| Delta(14)-sterol reductase TM7SF2 | Tm7sf2 | 2.28±0.76* | 0.85±0.18 | 0.78±0.20 |
| CDGSH iron-sulfur domain-containing protein 1 | Cisd1 | 2.27±1.41 | 1.61±1.32 | 1.45±1.04 |

| | | | | |
|---|---|---|---|---|
| Pro-cathepsin H | Ctsh | 2.27±0.47* | 1.09±0.25 | 1.41±0.50 |
| NAD-dependent malic enzyme, mitochondrial | Me2 | 2.24±2.73 | 1.95±2.35 | 2.54±3.24 |
| Cytochrome P450 2F2 | Cyp2f2 | 2.23±1.27 | 1.64±0.98 | 1.18±0.74 |
| WD repeat-containing protein 18 | Wdr18 | 2.22±2.64 | 2.18±2.66 | 2.03±2.17 |
| Cadherin EGF LAG seven-pass G-type receptor 3[#] | Celsr3 | 2.21±1.48 | 1.78±1.60 | 1.65±1.18 |
| Endonuclease G, mitochondrial | Endog | 2.21±1.64 | 1.56±1.14 | 1.40±1.03 |
| Keratin, type I cytoskeletal 28 | Krt28 | 2.21±3.17 | 0.81±0.55 | 1.74±1.37 |
| Acylcarnitine hydrolase | Ces2c | 2.20±0.95* | 0.93±0.09 | 0.92±0.09 |
| Sarcosine dehydrogenase, mitochondrial | Sardh | 2.19±0.59* | 1.45±0.33* | 0.95±0.24 |
| ER membrane protein complex subunit 7 | Emc7 | 2.17±2.81 | 1.88±2.23 | 1.92±2.37 |
| Beta-mannosidase | Manba | 2.16±1.19 | 0.71±0.09 | 0.86±0.30 |
| Alpha-N-acetylgalactosaminidase | Naga | 2.16±0.36* | 0.89±0.07 | 1.06±0.17 |
| Keratin, type II cytoskeletal 2 epidermal | Krt2 | 2.15±1.84 | 1.47±1.86 | 4.58±8.23 |
| Kynurenine 3-monooxygenase | Kmo | 2.14±2.47 | 1.74±1.64 | 1.34±1.22 |
| Glycerophosphocholine phosphodiesterase GPCPD1 | Gpcpd1 | 2.14±1.59 | 0.79±0.39 | 0.70±0.35 |
| ATP synthase subunit f, mitochondrial | Atp5mf | 2.13±2.20 | 1.94±1.94 | 1.80±2.03 |
| Small nuclear ribonucleoprotein Sm D1 | Snrpd1 | 2.13±1.84 | 2.05±2.04 | 2.03±2.00 |
| CD302 antigen | Cd302 | 2.12±1.72 | 0.86±0.49 | 1.08±0.91 |
| Parathymosin | Ptms | 2.12±1.66 | 1.92±1.69 | 1.68±1.45 |
| Hydroxyacid oxidase 1 | Hao1 | 2.11±0.90* | 0.99±0.45 | 1.15±0.34 |
| Histidine triad nucleotide-binding protein 2, mitochondrial | Hint2 | 2.10±0.35* | 0.99±0.14 | 0.93±0.18 |
| Pyridoxine-5'-phosphate oxidase | Pnpo | 2.10±0.67* | 1.23±0.16* | 1.00±0.04 |
| Protein-glutamine gamma-glutamyltransferase K | Tgm1 | 2.10±0.62* | 0.82±0.27 | 0.91±0.20 |
| Serine hydrolase-like protein[#] | Serhl | 2.10±2.09 | 1.94±1.62 | 2.05±1.88 |
| Mitochondrial chaperone BCS1 | Bcs1l | 2.09±1.82 | 1.65±1.32 | 1.54±1.28 |
| Ubiquitin-like modifier-activating enzyme ATG7 | Atg7 | 2.07±1.35 | 1.38±0.73 | 1.41±0.88 |
| Argininosuccinate synthase | Ass1 | 2.05±1.64 | 1.92±0.75* | 0.90±0.38 |
| Acyl-CoA-binding protein | Dbi | 2.05±2.30 | 2.20±3.02 | 2.05±2.43 |
| UDP-glucuronosyltransferase 1-1 | Ugt1a1 | 2.05±1.31 | 1.34±0.88 | 1.32±0.83 |
| Acetyl-coenzyme A transporter 1 | Slc33a1 | 2.04±2.75 | 2.20±2.51 | 2.13±2.62 |
| Phenylalanine-4-hydroxylase | Pah | 2.03±1.32 | 1.42±0.79 | 1.21±0.73 |
| Regucalcin | Rgn | 2.03±0.80* | 1.08±0.59 | 0.82±0.31 |
| Arginase-1 | Arg1 | 2.02±0.50* | 1.21±0.25 | 0.97±0.23 |
| Keratin, type I cytoskeletal 16 | Krt16 | 2.02±1.83 | 0.80±0.58 | 4.30±6.06 |
| S-methylmethionine--homocysteine S-methyltransferase BHMT2 | Bhmt2 | 2.01±2.67 | 2.51±3.26 | 2.70±3.60 |
| Medium-chain acyl-CoA ligase ACSF2, mitochondrial | Acsf2 | 2.01±0.59* | 0.89±0.14 | 1.05±0.21 |
| Glutathione S-transferase Mu 7 | Gstm7 | 2.00±1.36 | 1.79±1.50 | 1.47±1.11 |

| | | | | |
|---|---|---|---|---|
| Mitochondrial import receptor subunit TOM5 homolog | Tomm5 | 2.00±2.27 | 1.97±2.01 | 1.93±2.13 |

These values represent average (± standard deviation) fold-change of abundance ratios for each altered protein in C57BL6 mice on low-fat compared to the high-fat control group (MS-NASH mice on high-fat) with a cutoff of 2-fold-change. For each upregulated protein, corresponding values from the other two groups (on high-fat diet) are shown for comparison. The liver samples (from 5 mice in each group) were individually assessed by TMT based differential proteomic expression and data were merged to get averages. Protein FDR Confidence for all proteins was ≤1% except 8 proteins (#≤2%). These data are also presented in Figure 3A. FDR: False Discovery Rate.
