## Supplemental Table 11 for "Liver Protein Expression in Nash Mice on a High-Fat Diet"

**Supplement Table 11. Downregulated Proteins by an unbiased proteomic screening of C57BL6 mice on low-fat diet**

| | | C57BL6 | MS-NASH | |
|---|---|---|---|---|
| **Proteins** | **Genes** | **Control** | **OCA** | **Aquamin** |
| Leucine-rich repeat-containing protein 9[#] | Lrrc9 | 0.046±0.013* | 0.978±0.235 | 0.705±0.168* |
| Histone H2A type 2-A | Hist2h2aa1 | 0.181±0.229* | 0.878±0.500 | 0.941±0.590 |
| Cytochrome P450 2B9 | Cyp2b9 | 0.182±0.068* | 0.751±0.219* | 1.076±0.412 |
| Perilipin-2 | Plin2 | 0.184±0.013* | 0.559±0.096* | 0.864±0.107* |
| Cornifin-A | Sprr1a | 0.207±0.148* | 0.920±0.570 | 1.341±0.634 |
| Thymidine phosphorylase | Tymp | 0.216±0.038* | 1.633±0.324* | 0.939±0.160 |
| Lymphocyte antigen 6D | Ly6d | 0.221±0.094* | 1.116±0.828 | 1.693±0.866 |
| Histone H2B type 1-B | H2bc3 | 0.227±0.251* | 0.840±0.449 | 0.931±0.614 |
| SCY1-like protein 2 | Scyl2 | 0.231±0.131* | 1.032±0.144 | 1.028±0.080 |
| Son of sevenless homolog 1[#] | Sos1 | 0.233±0.096* | 0.605±0.115* | 1.015±0.134 |
| Keratin, type I cytoskeletal 20 | Krt20 | 0.236±0.167* | 0.925±0.343 | 0.971±0.256 |
| PDZ and LIM domain protein 7 | Pdlim7 | 0.241±0.121* | 0.804±0.342 | 1.123±0.265 |
| Branched-chain-amino-acid aminotransferase, mitochondrial | Bcat2 | 0.250±0.211* | 0.604±0.594 | 0.656±0.549 |
| Oxysterol-binding protein-related protein 3 | Osbpl3 | 0.255±0.087* | 0.627±0.467 | 1.297±0.707 |
| Histone H2AX | H2ax | 0.257±0.253* | 0.931±0.515 | 1.056±0.712 |
| Centrosomal protein of 170 kDa | Cep170 | 0.260±0.244* | 0.461±0.341 | 0.672±0.529 |
| Perilipin-4 | Plin4 | 0.270±0.170* | 0.502±0.264* | 1.445±1.157 |
| Lymphocyte-specific protein 1 | Lsp1 | 0.272±0.136* | 0.798±0.420 | 1.091±0.551 |
| Myosin regulatory light polypeptide 9 | Myl9 | 0.274±0.056* | 0.936±0.603 | 0.939±0.406 |
| Aquaporin-4 | Aqp4 | 0.285±0.197* | 3.992±3.087 | 0.921±0.561 |
| Cysteine and glycine-rich protein 1 | Csrp1 | 0.292±0.208* | 0.695±0.207* | 1.097±0.307 |
| Histone H2A.V | H2az2 | 0.295±0.264* | 0.993±0.330 | 1.076±0.663 |
| Acyl-CoA desaturase 1 | Scd1 | 0.296±0.141* | 0.720±0.322 | 0.828±0.212 |
| Trinucleotide repeat-containing gene 6A protein[#] | Tnrc6a | 0.304±0.193* | 0.663±0.455 | 0.809±0.607 |
| Cytochrome P450 2C38 | Cyp2c38 | 0.305±0.177* | 1.332±0.317 | 1.084±0.283 |
| Apolipoprotein C-II | Apoc2 | 0.311±0.147* | 1.155±0.593 | 1.180±0.433 |
| Elongation of very long chain fatty acids protein 5 | Elovl5 | 0.311±0.087* | 1.058±0.222 | 0.965±0.239 |
| Interferon alpha-inducible protein 27-like protein 2B | Ifi27l2b | 0.333±0.070* | 0.909±0.194 | 1.124±0.310 |
| Tonsoku-like protein | Tonsl | 0.342±0.232* | 0.306±0.172* | 0.512±0.333 |
| 2'-deoxynucleoside 5'-phosphate N-hydrolase 1 | Dnph1 | 0.344±0.214* | 1.428±0.860 | 0.897±0.489 |
| Brain acid soluble protein 1 | Basp1 | 0.347±0.101* | 0.864±0.223 | 1.184±0.333 |
| Histone H2A type 1-F | Hist1h2af | 0.355±0.388* | 0.647±0.432 | 0.818±0.511 |
| Fatty acid synthase | Fasn | 0.356±0.145* | 1.501±0.692 | 1.171±0.425 |

| | | | | |
|---|---|---|---|---|
| Protein ABHD1 | Abhd1 | 0.356±0.146* | 0.995±0.116 | 0.981±0.165 |
| Uridine-cytidine kinase 1 | Uck1 | 0.362±0.083* | 0.695±0.077* | 0.928±0.119 |
| Casein kinase I isoform alpha | Csnk1a1 | 0.364±0.271* | 0.972±0.207 | 0.892±0.140 |
| Protein phosphatase 1 regulatory subunit 12C | Ppp1r12c | 0.368±0.297* | 0.870±0.098 | 0.908±0.144 |
| Synaptopodin[#] | Synpo | 0.368±0.036* | 0.812±0.351 | 1.250±0.587 |
| Regulator of G-protein signaling 10 | Rgs10 | 0.369±0.196* | 0.856±0.278 | 1.296±0.198 |
| Cytochrome P450 2C39 | Cyp2c39 | 0.373±0.132* | 1.365±0.175 | 1.029±0.185 |
| Emopamil-binding protein-like | Ebpl | 0.376±0.212* | 0.724±0.108* | 0.867±0.125* |
| Histone H4 | H4c1 | 0.378±0.410* | 0.851±0.341 | 0.936±0.620 |
| Plastin-1 | Pls1 | 0.380±0.345* | 1.401±0.599 | 1.398±0.404 |
| Glycine amidinotransferase, mitochondrial | Gatm | 0.387±0.211* | 0.732±0.431 | 0.876±0.486 |
| Death-associated protein 1 | Dap | 0.389±0.146* | 1.013±0.459 | 0.924±0.470 |
| Annexin A2 | Anxa2 | 0.390±0.119* | 0.826±0.272 | 1.313±0.171 |
| Diacylglycerol O-acyltransferase 1 | Dgat1 | 0.392±0.319* | 0.835±0.266 | 0.999±0.428 |
| Histone H3.1 | H3c1 | 0.392±0.418* | 0.791±0.422 | 0.936±0.741 |
| Monoglyceride lipase | Mgll | 0.394±0.079* | 0.767±0.123* | 0.942±0.151 |
| Protein LSM12 homolog | Lsm12 | 0.394±0.257* | 0.954±0.108 | 1.058±0.202 |
| Ena/VASP-like protein | Evl | 0.397±0.194* | 0.927±0.705 | 1.309±0.750 |
| STE20/SPS1-related proline-alanine-rich protein kinase | Stk39 | 0.398±0.315* | 0.885±0.634 | 1.204±0.786 |
| Heme-binding protein 1 | Hebp1 | 0.398±0.064* | 1.110±0.264 | 0.989±0.193 |
| Histone H3.2 | H3c2 | 0.399±0.280* | 0.884±0.193 | 0.960±0.032 |
| Acyl-coenzyme A thioesterase 2, mitochondrial | Acot2 | 0.400±0.123* | 1.022±0.290 | 1.289±0.436 |
| Collagen alpha-1(XXV) chain[#] | Col25a1 | 0.402±0.221* | 0.915±0.334 | 0.903±0.369 |
| Annexin A1 | Anxa1 | 0.403±0.156* | 0.716±0.208* | 1.037±0.381 |
| Triokinase/FMN cyclase | Tkfc | 0.404±0.099* | 1.472±0.355 | 1.213±0.317 |
| Sorbin and SH3 domain-containing protein 1 | Sorbs1 | 0.410±0.092* | 0.895±0.123 | 0.923±0.226 |
| Acyl-coenzyme A thioesterase 11 | Acot11 | 0.412±0.058* | 0.924±0.133 | 0.980±0.240 |
| 17-beta-hydroxysteroid dehydrogenase 13 | Hsd17b13 | 0.412±0.177* | 0.916±0.286 | 1.256±0.304 |
| ATP-citrate synthase | Acly | 0.413±0.098* | 1.367±0.295 | 1.065±0.253 |
| Charged multivesicular body protein 2b | Chmp2b | 0.416±0.219* | 1.072±0.156 | 1.378±0.229 |
| Pyridine nucleotide-disulfide oxidoreductase domain-containing protein 2 | Pyroxd2 | 0.420±0.324* | 1.061±0.076 | 0.950±0.025* |
| Acetyl-CoA carboxylase 1 | Acaca | 0.420±0.111* | 1.152±0.300 | 1.096±0.188 |
| Fatty acid-binding protein, intestinal | Fabp2 | 0.421±0.136* | 1.233±0.062 | 1.128±0.067 |
| Apolipoprotein A-IV | Apoa4 | 0.423±0.125* | 1.238±0.432 | 1.706±0.733 |
| Quinone oxidoreductase-like protein 1 | Cryzl1 | 0.423±0.299* | 0.650±0.491 | 0.514±0.430 |
| Sterile alpha motif domain-containing protein 9-like | Samd9l | 0.426±0.077* | 1.178±0.168 | 1.415±0.144* |
| High mobility group protein HMG-I/HMG-Y | Hmga1 | 0.426±0.209* | 1.460±0.331 | 1.105±0.231 |

| | | | | |
|---|---|---|---|---|
| Rho guanine nucleotide exchange factor 2 | Arhgef2 | 0.427±0.135* | 0.744±0.358 | 1.205±0.350 |
| Acyl-CoA synthetase short-chain family member 3, mitochondrial | Acss3 | 0.429±0.111* | 1.082±0.234 | 1.083±0.400 |
| Histone H1.5 | H1-5 | 0.430±0.145* | 1.010±0.181 | 1.211±0.266 |
| ATP-binding cassette sub-family D member 1 | Abcd1 | 0.430±0.161* | 0.942±0.180 | 0.934±0.212 |
| Transmembrane protein 230 | Tmem230 | 0.431±0.199* | 0.877±0.186 | 0.867±0.280 |
| Peroxisomal bifunctional enzyme | Ehhadh | 0.431±0.168* | 0.858±0.240 | 1.110±0.395 |
| Galectin-3 | Lgals3 | 0.434±0.161* | 0.840±0.280 | 1.430±0.391 |
| Zyxin | Zyx | 0.435±0.053* | 1.089±0.210 | 1.076±0.150 |
| FYN-binding protein 2 | Fyb2 | 0.436±0.158* | 1.275±0.175 | 1.055±0.305 |
| Lysophosphatidylcholine acyltransferase 1 | Lpcat1 | 0.437±0.221* | 1.094±0.292 | 1.053±0.332 |
| NudC domain-containing protein 3 | Nudcd3 | 0.439±0.150* | 0.913±0.092 | 1.076±0.147 |
| D-3-phosphoglycerate dehydrogenase | Phgdh | 0.440±0.375 | 0.551±0.464 | 0.690±0.600 |
| Melanoma-associated antigen D1 | Maged1 | 0.441±0.184* | 1.002±0.422 | 1.043±0.639 |
| Prostaglandin reductase 1 | Ptgr1 | 0.443±0.093* | 0.847±0.153 | 1.123±0.187 |
| Epididymal-specific lipocalin-10[#] | Lcn10 | 0.444±0.466 | 0.918±0.787 | 0.745±0.695 |
| Helicase-like transcription factor | Hltf | 0.445±0.168* | 0.946±0.458 | 0.890±0.079 |
| Histone H3.3 | H3-3a | 0.449±0.505* | 0.875±0.463 | 0.894±0.586 |
| Myosin regulatory light chain 12B | Myl12b | 0.450±0.127* | 1.029±0.255 | 1.131±0.215 |
| CTTNBP2 N-terminal-like protein | Cttnbp2nl | 0.453±0.151* | 0.832±0.154 | 1.035±0.263 |
| Bromodomain-containing protein 3 | Brd3 | 0.453±0.068* | 1.041±0.254 | 0.856±0.349 |
| WAS/WASL-interacting protein family member 3 | Wipf3 | 0.454±0.107* | 1.289±0.481 | 1.201±0.559 |
| Endonuclease 8-like 2 | Neil2 | 0.455±0.255* | 1.209±0.434 | 1.231±0.724 |
| Sulfotransferase 1C2 | Sult1c2 | 0.455±0.135* | 0.984±0.371 | 1.092±0.407 |
| Reticulon-4 | Rtn4 | 0.456±0.107* | 0.793±0.107* | 1.108±0.178 |
| A-kinase anchor protein 2 | Akap2 | 0.456±0.068* | 0.833±0.065* | 1.004±0.197 |
| Sphingomyelin phosphodiesterase 3 | Smpd3 | 0.459±0.153* | 1.224±0.148* | 0.972±0.253 |
| Apolipoprotein A-II | Apoa2 | 0.459±0.258* | 0.651±0.183* | 1.074±0.327 |
| Galectin-1 | Lgals1 | 0.461±0.192* | 0.488±0.266* | 1.038±0.387 |
| Phosphatidylethanolamine-binding protein 1 | Pebp1 | 0.461±0.204* | 1.263±0.771 | 1.378±0.930 |
| Serine/threonine-protein phosphatase 6 regulatory subunit 1 | Ppp6r1 | 0.462±0.225* | 1.016±0.279 | 0.975±0.220 |
| Microtubule-associated protein 4 | Map4 | 0.462±0.115* | 0.955±0.142 | 1.120±0.275 |
| Rho GTPase-activating protein 23 | Arhgap23 | 0.463±0.096* | 0.767±0.135* | 1.155±0.152 |
| 85/88 kDa calcium-independent phospholipase A2 | Pla2g6 | 0.464±0.115* | 1.045±0.221 | 0.908±0.333 |
| PDZ domain-containing protein GIPC1 | Gipc1 | 0.466±0.110* | 0.904±0.140 | 1.009±0.210 |
| Collagen alpha-1(XIV) chain | Col14a1 | 0.466±0.036* | 1.155±0.543 | 1.815±0.964 |
| Adenylate kinase isoenzyme 1 | Ak1 | 0.467±0.392 | 0.656±0.522 | 0.812±0.649 |
| H-2 class II histocompatibility antigen gamma chain | Cd74 | 0.470±0.182* | 0.971±0.774 | 2.733±3.743 |

| | | | | |
|---|---|---|---|---|
| Zinc finger CCCH domain-containing protein 11A | Zc3h11a | 0.471±0.214* | 0.939±0.278 | 1.075±0.352 |
| Uncharacterized protein KIAA1522 | Kiaa1522 | 0.474±0.279* | 0.994±0.278 | 1.207±0.614 |
| Microtubule-associated protein tau | Mapt | 0.475±0.132* | 0.960±0.315 | 1.776±0.814 |
| Protein S100-A10 | S100a10 | 0.476±0.073* | 0.857±0.361 | 1.353±0.302 |
| Glucose-6-phosphatase | G6pc | 0.476±0.098* | 0.801±0.156* | 0.893±0.173 |
| Arf-GAP domain and FG repeat-containing protein 1 | Agfg1 | 0.478±0.260* | 0.957±0.291 | 0.951±0.309 |
| GRB10-interacting GYF protein 2 | Gigyf2 | 0.479±0.148* | 0.979±0.225 | 0.972±0.339 |
| Actin-binding LIM protein 1 | Ablim1 | 0.480±0.137* | 0.987±0.071 | 0.990±0.125 |
| Testis-expressed protein 2 | Tex2 | 0.482±0.124* | 0.956±0.113 | 0.933±0.158 |
| Band 4.1-like protein 1 | Epb41l1 | 0.485±0.163* | 1.628±0.540 | 1.236±0.474 |
| Neuronal proto-oncogene tyrosine-protein kinase Src | Src | 0.486±0.515 | 1.418±1.088 | 1.441±0.757 |
| Glycogen synthase kinase-3 alpha | Gsk3a | 0.489±0.280* | 1.191±0.427 | 1.233±0.208 |
| Nucleoporin NUP35 | Nup35 | 0.492±0.205* | 0.930±0.359 | 0.900±0.226 |
| Protein farnesyltransferase subunit beta | Fntb | 0.493±0.362 | 0.684±0.446 | 0.616±0.428 |
| Peroxisomal membrane protein 11A | Pex11a | 0.493±0.078* | 0.657±0.092* | 0.957±0.034* |
| Coronin-1A | Coro1a | 0.494±0.210* | 0.798±0.389 | 1.131±0.466 |
| ATP-binding cassette sub-family D member 2 | Abcd2 | 0.495±0.065* | 0.924±0.148 | 1.236±0.287 |
| Serpin H1 | Serpinh1 | 0.497±0.208* | 0.819±0.171* | 1.141±0.224 |
| Proteolipid protein 2[#] | Plp2 | 0.498±0.393* | 0.644±0.364 | 1.007±0.478 |

These values represent average (± standard deviation) fold-change of abundance ratios for each altered protein in C57BL6 mice on low-fat compared to the high-fat control group (MS-NASH mice on high-fat) with a cutoff of 2-fold-change. For each downregulated protein, corresponding values from the other two groups (on high-fat diet) are shown for comparison. The liver samples (from 5 mice in each group) were individually assessed by TMT based differential proteomic expression and data were merged to get averages. Protein FDR Confidence for all proteins was ≤1% except 7 proteins ([#]≤2%). These data are also presented in Figure 3A. FDR: False Discovery Rate
