## Supplemental Table 12 for "Liver Protein Expression in Nash Mice on a High-Fat Diet"

**Supplement Table 12. Top pathways associated with upregulated proteins altered with low-fat diet in C57BL6 mice**

| Pathway name | Entities pValue | Mapped entities |
|---|---|---|
| Metabolism | $4.84\times10^{-10}$ | Apom;Ces3b;Otc;Kmo;Ak4;Hao1;Nsdhl;Aass;Nme2;Lss;Glrx;Ass1;Rap1a;Akr1b8;Ugt3a2;Cps1;Gnmt;Cyp1a2;Fdps;Cyp2f2;Mmab;Gba;Ivd;Ugt1a9;Gls2;Cyp51a1;Cyp7b1;Mvk;Me2;Hoga1;Msmo1;Isyna1;Arsb;Aadat;Abcd4;Pck1;Cox6a1;Kyat1;Aadac;Acsf2;Nudt16;Cyp4v2;Mtnd5;Atp5mf;Sardh;Gstm7;Psmb3;Pnpo;Gstp1;Arg1;Man2b2;Fxn;Agxt;Idi1;Galt;Ugt1a1;Ppcdc;Upb1;Cpt2;Manba;Gcsh;Cth;Dbi;Acsm1;Fabp5;Pah;Btd;Tm7sf2;Mt-Cyb;Pemt;Bhmt2;Tat |
| Metabolism of amino acids and derivatives | $4.63\times10^{-8}$ | Gnmt;Sardh;Kmo;Otc;Psmb3;Hao1;Gcsh;Ivd;Arg1;Aass;Gls2;Cth;Hoga1;Agxt;Pah;Aadat;Ass1;Bhmt2;Tat;Kyat1;Cps1 |
| Cholesterol biosynthesis | $5.63\times10^{-7}$ | Msmo1; Lss;Fdps;Idi1;Tm7sf2;Nsdhl;Cyp51a1;Mvk |
| Formation of the cornified envelope | $2.61\times10^{-7}$ | Krt2; Krt1;Krt79;Krt16;Tgm1;Krt5;Krt76;Krt17;Krt10; Krt14;Krt77;Krt28 |
| Urea cycle | $5.62\times10^{-5}$ | Ass1;Otc;Arg1;Cps1 |
| Metabolism of steroids | $1.58\times10^{-4}$ | Msmo1; Lss;Fdps;Idi1;Tm7sf2;Akr1b8;Nsdhl;Cyp51a1;Cyp7b1;Mvk |
| Keratinization | $2.03\times10^{-4}$ | Krt2; Krt1;Krt79;Krt16;Tgm1;Krt5;Krt76;Krt17;Krt10; Krt14;Krt77;Krt28 |
| Biological oxidations | $3.84\times10^{-4}$ | Cyp1a2;Ugt1a1;Cyp4v2;Ces3b;Gstm7;Cyp2f2;Gstp1;Ugt1a9;Cyp51a1;Cyp7b1;Acsm1;Ugt3a2;Aadac |
| Glyoxylate metabolism and glycine degradation | $3.87\times10^{-4}$ | Gnmt;Hoga1;Agxt;Hao1;Gcsh |
| Phenylalanine and tyrosine metabolism | 0.002 | Pah;Tat;Kyat1 |
| Tryptophan catabolism | 0.002 | Aadat;Kmo;Kyat1 |
| Lysosomal oligosaccharide catabolism | 0.003 | Man2b2;Manba |
| Phase II - Conjugation of compounds | 0.004 | Cyp1a2;Ugt1a1;Gstm7;Gstp1;Ugt1a9;Ugt3a2;Acsm1 |
| Phenylalanine metabolism | 0.007 | Pah;Kyat1 |
| Phase I - Functionalization of compounds | 0.007 | Cyp1a2;Cyp4v2;Ces3b;Cyp2f2;Cyp51a1;Aadac;Cyp7b1 |
| Glucuronidation | 0.010 | Ugt1a1;Ugt1a9;Ugt3a2 |
| Terminal pathway of complement | 0.012 | C8b;C8g |
| Cytochrome P450 - arranged by substrate type | 0.016 | Cyp1a2;Cyp4v2;Cyp2f2;Cyp51a1;Cyp7b1 |
| SLBP independent Processing of Histone Pre-mRNAs | 0.018 | Ncbp1;Snrpe |
| Mitochondrial protein import | 0.018 | Fxn;Otc |
| Endogenous sterols | 0.018 | Cyp4v2;Cyp51a1;Cyp7b1 |
| Interconversion of nucleotide di- and triphosphates | 0.019 | Nme2;Glrx;Ak4 |
| Aromatic amines can be N-hydroxylated or N-dealkylated by CYP1A2 | 0.020 | Cyp1a2 |

| | | |
|---|---|---|
| SLBP Dependent Processing of Replication-Dependent Histone Pre-mRNAs | 0.021 | Ncbp1;Snrpe |
| Type I hemidesmosome assembly | 0.021 | Krt5; Krt14 |
| mRNA Splicing - Minor Pathway | 0.022 | Sf3b4;Ncbp1;Snrpe;Snrpd1 |
| Lysine catabolism | 0.025 | Aadat;Aass |
| Cobalamin (Cbl, vitamin B12) transport and metabolism | 0.025 | Mmab;Abcd4 |
| Glutamate and glutamine metabolism | 0.029 | Gls2;Kyat1 |
| Conjugation of phenylacetate with glutamine | 0.039 | Acsm1 |
| Cysteine formation from homocysteine | 0.039 | Cth |
| Degradation of GABA | 0.039 | Abat |
| Developmental Biology | 0.045 | Krt1;Krt16;Tgm1;Rras;Krt76; Krt14;Krt77;Krt2;Egfr;Krt79;Krt5;Tubb4a;Krt17;Krt10;Krt28;Ap2s1 |
| Metabolism of nucleotides | 0.046 | Nudt16;Nme2;Glrx;Upb1;Ak4 |

The pathways listed here are altered by the significantly upregulated proteins with the control low-fat diet in C57BL6 mice presented in Supplement Table 10 using an unbiased approach. Reactome (v78) was used to generate the pathway analysis report for species *Mus musculus*. The significance (*p*-value) is calculated by the overrepresentation analysis (hypergeometric distribution).
